# Beyond power: Multivariate discovery, replication, and interpretation of pleiotropic loci using summary association statistics

**DOI:** 10.1101/022269

**Authors:** Zheng Ning, Yakov A. Tsepilov, Sodbo Zh. Sharapov, Alexander K. Grishenko, Xiao Feng, Masoud Shirali, Peter K. Joshi, James F. Wilson, Yudi Pawitan, Chris S. Haley, Yurii S. Aulchenko, Xia Shen

## Abstract

The ever-growing genome-wide association studies (GWAS) have revealed widespread pleiotropy. To exploit this, various methods which consider variant association with multiple traits jointly have been developed. However, most effort has been put on improving discovery power: how to replicate and interpret these discovered pleiotropic loci using multivariate methods has yet to be discussed fully. Using only multiple publicly available single-trait GWAS summary statistics, we develop a fast and flexible multi-trait framework that contains modules for (i) multi-trait genetic discovery, (ii) replication of locus pleiotropic profile, and (iii) multi-trait conditional analysis. The procedure is able to handle any level of sample overlap. As an empirical example, we discovered and replicated 23 novel pleiotropic loci for human anthropometry and evaluated their pleiotropic effects on other traits. By applying conditional multivariate analysis on the 23 loci, we discovered and replicated two additional multi-trait associated SNPs. Our results provide empirical evidence that multi-trait analysis allows detection of additional, replicable, highly pleiotropic genetic associations without genotyping additional individuals. The methods are implemented in a free and open source R package MultiABEL.

**Author summary:** By analyzing large-scale genomic data, geneticists have revealed widespread pleiotropy, i.e. single genetic variation can affect a wide range of complex traits. Methods have been developed to discover such genetic variants. However, we still lack insights into the relevant genetic architecture - What more can we learn from knowing the effects of these genetic variants?

Here, we develop a fast and flexible statistical analysis procedure that includes discovery, replication, and interpretation of pleiotropic effects. The whole analysis pipeline only requires established genetic association study results. We also provide the mathematical theory behind the pleiotropic genetic effects testing.

Most importantly, we show how a replication study can be essential to reveal new biology rather than solely increasing sample size in current genomic studies. For instance, we show that, using our proposed replication strategy, we can detect the difference in genetic effects between studies of different geographical origins.

We applied the method to the GIANT consortium anthropometric traits to discover new genetic associations, replicated in the UK Biobank, and provided important new insights into growth and obesity.

Our pipeline is implemented in an open-source R package MultiABEL, sufficiently efficient that allows researchers to immediately apply on personal computers in minutes.

## Introduction

During the past decade, single-trait genome-wide association studies (GWASs) have successfully identified many genetic variants underlying complex traits [1]. However, there are many issues in the current GWAS procedure. For example, the effects of the genetic variants, such as single-nucleotide polymorphisms (SNPs), on complex traits are usually very small. This directly limits the discovery power in most GWASs. On the other hand, multiple GWASs suggest that pleiotropy is widespread for complex diseases and traits [2]. However, in standard single-trait GWAS, pleiotropy is not directly taken into account. Aiming to address these problems, many multi-trait analyses methods have been developed in recent years to jointly analyze multiple correlated phenotypes. At the early stage, most multi-trait tools were based on individual-level data. For example, mv-plink implements canonical correlation analysis (CCA) to identify the association between each SNP and linear combinations of traits [3]; MultiPhen [4] performs a reversed regression with SNP as outcome and phenotypes as predictors and combined-PC [5] where a principle components analysis is done on the phenotype data to improve statistical power. A simulation study [6] demonstrated that the statistical power of these methods is very similar to the power of the standard Multivariate Analysis of Variance (MANOVA) for multiple phenotypes on each common SNP.

As more and more GWASs have been done in different study populations, given the difficulty of sharing individual-level data, multi-trait methods based on summary-level data have become popular. In order to combine any set of single-trait GWAS summary-level data, the method should be able to (1) efficiently meta-analyze an arbitrary number of phenotypes, (2) combine any phenotypic distributions including quantitative and case-control outcomes, (3) handle any level of sample overlap between studies, and (4) do not rely on known sample size knowledge or on strong assumptions. Desired features (1) and (2) are computationally challenging for most multivariate methods, and more importantly, (3) and (4) have to be achieved to take full advantage of all the established GWAS results. There are several methods fulfilling most of these requirements. For example, Stephens (2013) outlined a unified multivariate analysis framework based on Bayesian model comparison [7]; Zhu et al. (2015) introduced two test statistics *S_Hom_* and *S_Het_* to improve statistical power under different assumptions of effect sizes [8], and suggested seven new loci by jointly analyzing the summary statistics of three traits from GIANT [9]; Multi-trait analysis of GWAS (MTAG) was developed to integrate the GWAS summary results of several related traits and improve the inference in each single-trait GWAS [10].

Although most summary-level multi-trait methods can boost discovery power, the replication strategy and interpretation of the loci discovered in multi-trait analysis have yet to be developed and agreed upon. When a locus is significant in multi-trait analysis, a trivial approach to replication is to replicate associations trait by trait. However, in this way, the overall association pattern between a SNP and multiple phenotypes is not replicated. Another straightforward way for replication is performing the multi-trait test in replication sample, and asking for the overall association (omnibus p-value) to be significant. Although this strategy is at multivariate level, it does not replicate the discovered locus pleiotropic profile either, as the consistency of the effect sizes and effect directions across samples are neglected. Even if the effect sizes and directions of effects are completely different in replication sample, the multivariate test may still reject the null hypothesis of no association between the SNP and any phenotypes. Therefore, neither of these two trivial replication strategies replicates the discovered locus pleiotropic profile properly.

Furthermore, loci detected in multi-trait analysis is usually interpreted at the single-trait level. Given many phenotypes are defined with numerous underlying biological factors, O’Reilly et al. [4] point out that linear combinations of phenotypes can be defined as new traits. Then the association between a SNP and a linear combination of phenotypes can be interpreted as an association between the SNP and a hidden phenotype which integrates the relevant underlying factors. Cichonska et al. [11] get CCA results using summary-level data, but how to interpret and replicate CCA results are not thoroughly discussed. Based on CCA, using individual-level data, we introduced a combined phenotype score to assess the genetic effect so that the replication can be meaningful [12]. More importantly, what geneticists want to acquire from multi-trait analyses is additional knowledge about pleiotropy, while such an aim is hardly reached by current multivariate methods.

Besides multi-trait analysis, various summary-level methods have been developed based on many classical statistical methods. For example, the conditional and joint multi-variant analysis (GCTA-COJO) [13] has been successful in discovering additional association signals within detected loci. In order to identify other trait-associated SNPs in linkage disequilibrium (LD) with the top SNP, GCTA-COJO performs a secondary association analysis conditioned on discovered top variants. For loci detected in multi-trait analysis, it will be helpful to perform similar conditional analysis to detect additional multi-trait associations instead of only reporting the top SNPs. Although several methods [14–16] have been developed to include multiple traits as covariates, only one trait is allowed to be dependent variable in these methods.

Here, using GWAS meta-analysis summary statistics, we develop a multi-trait analysis framework that integrates the discovery, replication and conditional analyses. We first develop a computationally fast method MVA (represents multivariate analysis) to get MANOVA results based on multiple GWAS results for different traits. We then introduce two replication strategies: MVA-score replication and Monte-Carlo (MC) based correlation replication to replicate the underlying pattern of associations between genotype and phenotypes. The MVA-score replication is related to CCA thus helpful for interpretation; and the correlation replication directly suggests that the relative effects and direction of associations are stable between discovery and replication samples. These two replication methods can be used to gain knowledge of pleiotropy from different aspects. We also develop and implement conditional multivariate analysis (cMVA) that performs a conditional analysis analog to GCTA-COJO [13] after MVA. All these procedures are solely based on summary association statistics from the discovery and replication studies. To empirically demonstrate the utility of our methods, we apply the methods on the publicly available GWAS summary-level data for human anthropometry, and replicate in the UK Biobank.

## Materials and Methods

### Summary of the methods

#### MVA

MVA is developed to get MANOVA results using summary statistics. To simplify the formulae, we assume the phenotypes are standardized to have mean zero and variance one, and genotypes are centered to have mean zero. *k* traits ***Y***_1_,…***Y***_*k*_ are dependent variables in MANOVA. We first focus on the analysis of one SNP. If we denote the true marginal effects on the k traits by ***β***, then the null hypothesis in MANOVA is *H*_0_: ***β*** = **0**. Let **t** = [*t*_1_,…,*t_k_*]′ be the vector of single-trait t-test statistics across the *k* phenotypes on the SNP **g**, and **R**\* Cor(t) = Var(t). If **R**\* is available, the test statistic

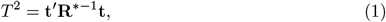

which asymptotically follows a *χ*^2^ distribution with *k* degrees of freedom under the null hypothesis.

#### Estimation of R*

Let **R** represent the phenotypic correlation matrix of the *k* phenotypes. According to Zhu et al. [8], **R**\* = **R** when the phenotypes are measured on the same set of individuals. If the individuals for trait *j* and those for trait *j*′ partially overlap, we denote the number of overlapping individuals as *n*_0_, those with trait *j* but not trait *j*′ as *n*_1_, and those with trait *j*′ but not trait *j* as *n*_2_. Then we have

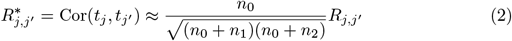

(S1 Appendix). Therefore, the correlation of t-statistics is a shrinkage version of the phenotypic correlation, with a factor determined by the level of overlap. Because (2) holds for all SNPs, an unbiased estimate of the correlation matrix **R**\* can be obtained by selecting a large number of independent variants from the meta-GWAS summary statistics and calculating their correlation coefficients (S1 Appendix).

#### cMVA

To detect additional associated SNPs at loci discovered in MVA, we introduce cMVA to perform conditional analysis. When *p* SNPs *G* = (*G*_1_,…,*G_p_*) and *k* traits are involved, for SNP *i*, we denote its t-test statistics conditional on the other *p* − 1 SNPs as 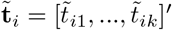. Both 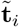 and its correlation matrix 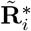 can be obtained by using summary statistics and a reference sample used to approximate the LD matrix Cor(*G*) (a full derivation is provided in the S1 Appendix). Therefore, similar to (1), we can use the test statistic

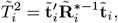

which asymptotically follows a *χ*^2^ distribution with *k* degrees of freedom under the null hypothesis.

#### Correlation replication

Aiming to replicate the locus pleiotropic profile in the replication sample, we develop this MC-based correlation replication strategy. The key idea is to evaluate the similarity of marginal effects across samples considering taking the uncertainty in estimates into account. When *p* SNPs and *k* traits are analyzed jointly, let 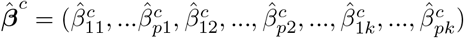 be the vector of estimated partial regression coefficients. Then for cMVA,

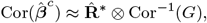

where ⊗ represents Kronecker product (S1 Appendix). Specifically, for MVA where *p* =1, 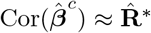. Because the variances of 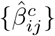 can be obtained from MVA or cMVA, we can get 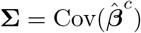. This allows us to draw 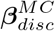 and 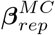 from 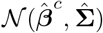 based on 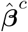 and 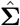 for the discovery sample and replication sample respectively. Then we can compute their correlation coefficients. Here we compute Kendall’s rank correlation coefficient

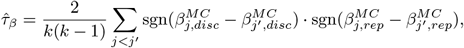

which measures the ordinal association between 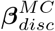 and 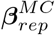. We choose Kendall’s correlation instead of Pearson’s correlation so that the correlation is not dominated by those traits with especially large effects. By performing parametric bootstrap simulations, we can get an estimated distribution of *τ_β_*. The parametric bootstrap confidence intervals (CI) based on this distribution can be used for inference.

#### MVA-score replication

If individual-level data are available, given a SNP, we can use CCA to get its most associated linear combination of traits. This linear combination of traits, which we name as MVA-score, can be seen as a new phenotype which is helpful for investigating the association between the SNP and the group of traits. It has been shown [4] that the coefficients in CCA are equivalent to the estimate of b in this reversed multiple regression

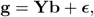

where g and **Y** represent genotypes and phenotypes respectively. Assuming Hardy-Weinberg equilibrium (HWE), 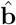 can be obtained by

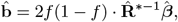

where *f* is the coding allele frequency of the SNP (S1 Appendix). Therefore, we can get 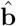 based on summary statistics and construct the MVA-score 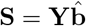. Taking **S** as a new phenotype and denoting the effect of the SNP on **S** as *β_s_*, we can estimate and test *β_s_* in the discovery and replication populations (S1 Appendix). If 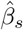 is significantly different from 0 and having the same sign in both populations, then we consider the association between the SNP and the MVA-score is replicated.

### Discovery cohort: GIANT

We downloaded the summary association statistics of six sex-stratified anthropometric traits meta-GWAS by the GIANT consortium from: https://www.broadinstitute.org/collaboration/giant/index.php/GIANT_consortium_data_files. We used six anthropometric traits: BMI, height, weight, hip circumference (denoted here as HIP), waist circumference (WC) and waist-to-hip ratio (WHR). We used two datasets with different sample size and meta-analysis date denoted as GIANT2013 and GIANT2015. For GIANT2013 data for all six traits from Randall et al. [17] was used. As sex stratified data was available, for each trait we computed the summary statistics by meta-analyzing the effects and standard errors of the two genders. For GIANT2015 we used several different data: height from Wood et al. [18]; HIP, WC and WHR from Shungin et al. [19], BMI from Locke et al. [20]. There wasn’t available summary statistics for weight later then 2013, therefore for GIANT2015 we have used the same weight GWAS results as for GIANT2013.

As HapMap II allele frequencies were reported in the meta-GWAS instead of pooled allele frequencies across all the cohorts, we excluded SNPs with sample size less than 40,000 and MAF<0.01 for GIANT2013 and sample size less than 70,000 and MAF<0.01 for GIANT2015. SNPs with missing allele frequencies were also excluded. All SNPs were merged with genome positions (GRCh37) and filtered for autosomals only and position missings. Then we selected only SNPs that were presented both in GIANT2013 and GIANT2015. In total we ended with 2,352,481 SNPs.

### Novel loci discovery and clumping

For comparison of MVA loci and UVA loci on GIANT2013 we used all SNPs above the threshold (p-value < 5 × 10^−8^). All SNPs were clumped to loci and compared with each other. We used position based clumping: (SNPs that are less than 500K distant from most significant SNP are considered as one locus).

For multivariate analysis of GIANT2015 we excluded all significant UVA SNPs with nearby region (p-value < 5 × 10^−8^). In total for GIANT2015 for six traits we found 28,658 significant SNPs. We removed these SNPs as well as all SNPs 500kb around (1Mb window): in total 618,873 SNPs were removed. All other SNPs were used for discovery multivariate analysis.

### Replication cohort: UK Biobank

UKB participants were recruited from the general UK population across 22 centers between 2006-2010. Subjects were aged 40-69 at baseline, underwent extensive phenotyping by questionnaire and clinic measurements, and provided a blood sample. Genotyping is in progress, with a wave 1 public release in June/July 2015. Data access to UKB was granted under MAF 8304. Phenotypes and genotypes were downloaded direct from UKB. In total 502,664 subjects had phenotypic information available, of whom 152,732 had been genotyped, of these 120,286 were identified as genetically British by UK Biobank, of which 118,182 (55,842 men) had complete phenotyping. These subjects were taken forward for analysis. Participants provided full informed consent to participate in UK Biobank. This study was covered by the generic ethical approval for UK Biobank studies from the NHS National Research Ethics Service (approval letter dated 17th June 2011, Ref 11/NW/0382). The authors in this study were completed blinded to the individual-level data collection and preparation. The phenotypes involved in this study were adjusted for age, sex and batch before being standardized to have mean zero and variance one.

### Reference cohort: 1000 Genomes

The 1000 Genomes Project was launched in 2008. The latest phases 3 data includes 2,504 individuals from 26 populations in Africa, East Asia, Europe, South Asia, and the Americas [21]. Both whole-genome sequencing and targeted exome sequencing have been done for all individuals. Among the sequenced individuals, 503 were identified as European ancestry samples. The genotypes of these subjects were used to approximate LD matrix in this study.

### Correlation matrix estimation

For correlation matrix estimation, we used previously proposed approach based on correlation of Z-statistics between independent unassociated SNPs [8]. We filtered SNPs based on given criteria: MAF > 0.1; high imputations quality as indicated either by INFO > 0.99 (for UKB) or by *N_e_/N* > 0.9, where *N_e_* = 1/(2*pq*(*se*)^2^), for GIANT; *abs*(*Z*) < 2; the sample of 200,000 independent (LD pruned) SNPs to compute correlation matrix (the list of SNPs was obtained by “–prune” option for PLINK using 1000 Genomes data). In total 128,670 independent SNPs were used to estimate correlation matrix for GIANT, and 166,000 SNPs for UKB. All estimated correlation matrices can be found in S3 Table.

### Multivariate trait construction

Together with six univariate traits we constructed six multivariate phenotypes based on these six anthropometric traits: all-six-traits (denoted as multi6 or m6), waist+hip+height+weight (denoted as multi4 or m4), waist+hip+whr (denoted as multi.shape3 or msh3), waist+hip (denoted as multi.shape2 or msh2), height+weight+bmi (denoted as multi.size3 or msz3), height+weight (denoted as multi.size2 or msz2).

### Application of cMVA at loci discovered in MVA

For each locus suggested in MVA, we set a 1-Mb window centred at the variant reported in Table 1 as the genomic locus to be analyzed. We perform cMVA to select the associated variants for each locus using the following stepwise selection strategy:

1. Take intersection of available variants in GIANT and 1000 Genomes European-ancestry samples.
2. Estimate LD correlations using an individual-level genotype data in 1000 Genomes.
3. Start with setting the remaining variants set as all the variants, and the selected variants set as empty.
4. Calculate the multivariate p-values of all the SNPs in the remaining set conditional on the SNPs in the selected set.
5. If the minimum multivariate conditional p-value is below a cutoff p-value, such as 5 × 10^−8^, then the corresponding SNP enters the selected set.
6. Calculate the conditional multivariate p-values of all the SNPs in the selected set. Those selected variants with p-values larger than the cutoff p-value are dropped.
7. To avoid collinearity, we need to filter the remaining variants. If the regression *R*^2^ between the selected SNPs and a remaining SNP is larger than 0.5, we remove the SNP from the remaining set.
8. Repeat 4-7 until the remaining and selected sets can no longer be changed.

**Table 1.**
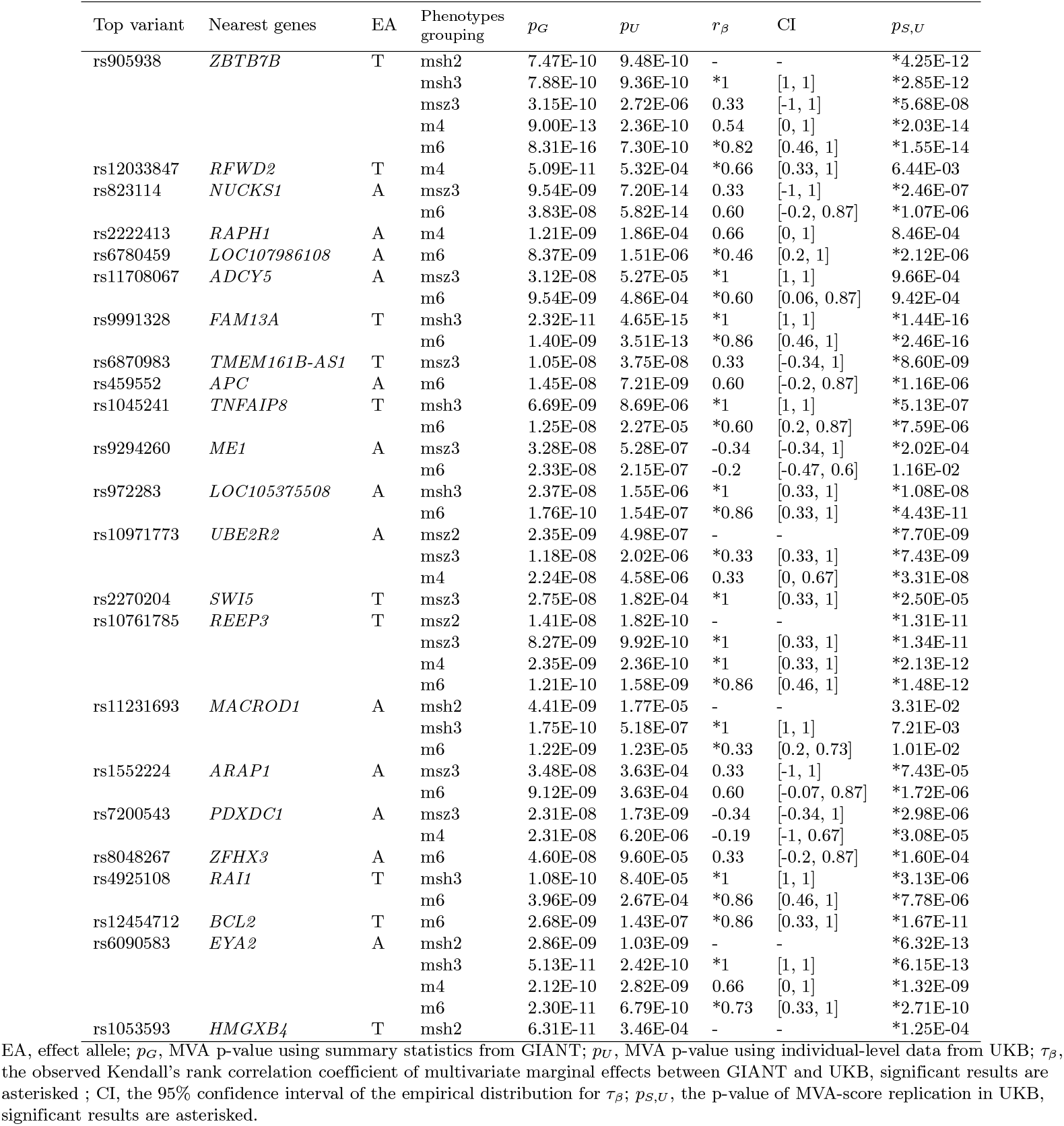
Summary of 23 loci detected and replicated by MVA for six anthropometric traits.

## Results

### Multivariate analysis of published results allows robust loci discovery

In order to test whether MVA of already published results allows for robust and fruitful discovery of new loci, we applied MVA to GWAS results published by the GIANT consortium in the previous wave of meta-analyses (GIANT2013) and used the latest, bigger analyses (GIANT2015) to validate our findings.

Using MVA, we have re-analyzed summary statistics for six traits in GIANT2013 data [17]: height (*n* = 133, 724), weight (*n* = 125, 946), body mass index (BMI, *n* = 126, 623), hip (*n* = 73, 209), waist (*n* = 85, 635), and waist-to-hip ratio (*n* = 77, 369). From these traits, we have constructed six multi-trait combinations related to size (SZ2: height and weight, SZ3: height, weight, BMI), shape (SH2: waist and hip; SH3: waist, hip, and WHR), and all parameters (M4: height, weight, waist, and hip; M6: the same as M4 and BMI and WHR). In contrast with MVA, single trait GWAS is referred as univariate analysis (UVA) in this study. In UVA, associations having nominal p-value < 5 × 10^−8^ were considered significant; we have used the same threshold for MVA.

Re-analysis of GIANT2013 data identified 72 new loci with significant MVA p-value for at least one multi-trait combination (S1 Table). In single-trait analysis, only 14 of these loci demonstrated suggestive (5 × 10^−8^ < *p* < 5 × 10^−7^) significance, and majority (47) had *p* > 1 × 10^−6^. The 72 loci were checked in the GIANT2015 analyses, that had roughly double sample sizes for all traits except for weight. The data used included GWAS for height [18] (*n* = 253, 108), body mass index (BMI [20], *n* = 233, 963), hip [19] (*n* = 145, 432), waist [19] (*n* = 153, 927), and waist-to-hip ratio [19] (*n* = 144, 578). We observed that most of loci (54 out of 72) discovered with MVA in GIANT2013 had genome-wide significant p-value < 5 × 10^−8^ with at least one of the traits in GIANT2015.

These results indicate that MVA re-analysis of published GWAS results may be a promising way to utilize already existing data and improve statistical power.

### Discovery of new anthropometric loci using published summary statistics

Given indications that MVA of summary-level data may allow for fruitful and robust loci discovery, we have set off to re-analyze the latest GWAS results published in GIANT2015. Re-analysis has led to identification of 49 new loci (Fig 1, S2 Table). We also quantified and analyzed the established associations of these loci using PhenoScanner [22] at a false discovery rate threshold of 0.05 (Fig 1, S4 Table, S3 Appendix, S4 Appendix). It is interesting to note that the yield of new loci is lower than that we had for the preliminary round of analyses, which we attribute to the fact that the overlap between samples is lower and hence the added value of MVA is lower in the latest GIANT data (e.g. see S3 Table for trait correlations estimated using three data sets used in this work).

**Fig 1.**
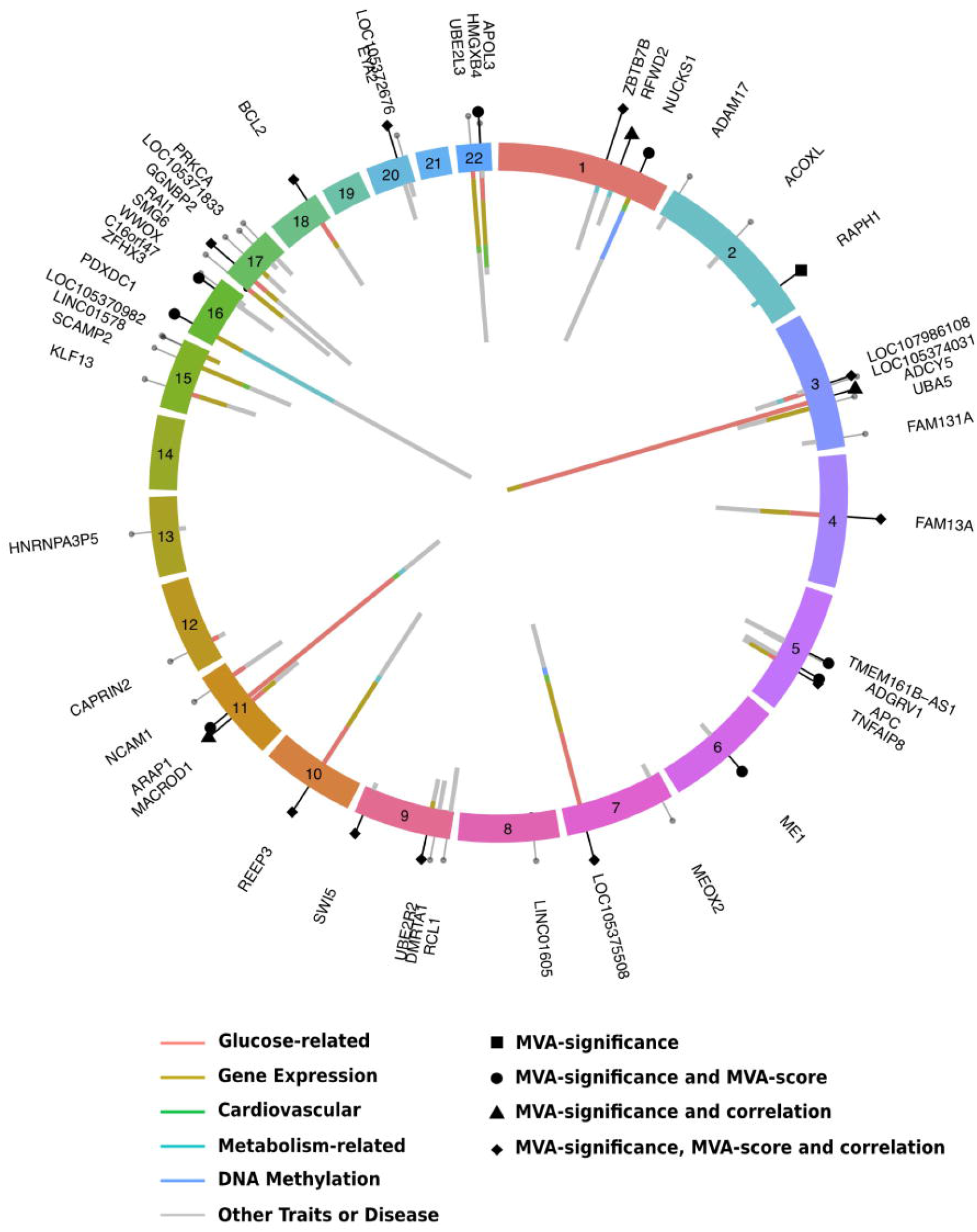
Novel associations discovered by the multi-trait analysis. Different chromosomes are displayed as circular chunks. The outside ring shows the newly detected loci, where the height of the bars are proportional to the multi-trait GWAS −log10 p-values for the most significant multi-trait combination. The 23 MVA replicated loci are represented as different shapes depending on replication strategies, and the rest are shown as small gray dots. The nearest genes of the top associated variants at these loci are labeled. The inside ring shows the amount of shared pleiotropic effects with other phenotypes in PhenoScanner at a 5% false discovery rate threshold.

### Replication of new anthropometric loci in the UK Biobank

We next attempted to replicate 49 new loci using the UK Biobank (UKB) interim release data (118,182 ethnically British, genetically Caucasian participants having all anthropometric measurements and genotypic data). For each locus, we defined the top SNP of a locus as the SNP with the smallest MVA p-value across all six multi-trait combinations. In the following replication, we used the top SNPs to represent loci.

There are various ways to replicate multivariate results. To begin with, we used two straightforward replication strategies: single-trait replication and MVA-significance replication. In the single-trait replication, a locus was replicated if there was at least one trait which was significantly associated with the top SNP in UKB, and the sign of association for this specific trait was consistent with that observed in GIANT2015. Because we have six traits, we set the single-trait replication p-value threshold as 0.05/(6 × 49) = 1.7 × 10^−4^. Using this criterion, we saw replication for 21 loci (S2 Table). Next, we used MVA-significance replication, where a locus was considered to be replicated if it has at least one replicable multi-trait combination. More specifically, a multi-trait combination is replicable if its MVA results are significant in both GIANT2015 and UKB. The replication p-value threshold of MVA-significance test was set to be 0.05/85 = 5.9 × 10^-4^. We adjusted the p-values by dividing 85 because for the 49 loci discovered in GIANT2015, there are in total 85 MVA-significant multi-trait combinations. Using this criterion, we replicated 23 loci (Table 1), three of which were not replicated in single-trait replication (S1 Fig). For some loci, there were multiple replicable multi-trait combinations. Although easy to implement, these two replication strategies have obvious drawbacks: the single-trait replication lacks power, and neither of these two strategies guarantees the genetic effects have similar sizes and directions as in the discovery population.

For the 23 loci replicated by the MVA-significance test, to test the consistency of pleiotropic effects, we implemented two novel replication strategies as follow-up methods of the MVA-significance replication: MVA-score replication, which we previously developed for individual-level data analysis [12], and MC-based correlation replication (S1 Fig). The MVA-score replication integrates the multivariate genetic effects into the genetic effect on MVA-score, and aims to replicate the discovered locus pleiotropic profile by testing this integrated effect; while the MC-based correlation replication tests the consistency of multivariate genetic effects directly. We firstly demonstrate the MVA-score replication, which can also be used for interpretation. For each replicable multi-trait combination of the 23 loci, we found the optimal linear combination of traits and computed its MVA-score (described in the Materials and Methods) in GIANT2015. For example, for six-traits MVA of rs905938,

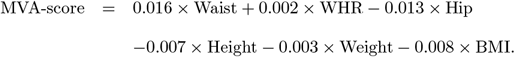

Then we used the same linear combination coefficients in UKB to get an MVA-score phenotype, estimated and tested the genetic effect *β_s_* of the SNP on the MVA-score in UKB. In this procedure, we consider results to be replicated when the association p-value is < 0.05/85 = 5.9 × 10^−4^ in UKB and the estimated *β_s_* has the same direction as in GIANT. In this way, we replicated 19 out of 23 loci (Table 1).

Next, for each replicable multi-trait combination of the 23 loci, we performed our MC-based correlation replication to test its consistency of pleiotropic association patterns between the discovery and replication populations. Based on the estimated marginal effect sizes and their variance in discovery sample and replication sample, we got a parametric bootstrap distribution of *τ_β_*, which is a measurement of the consistency of marginal effect sizes (described in the Materials and Methods). The CI of *τ_β_* can be used to evaluate the consistency of pleiotropic effects between discovery and replication samples. If 0 is not included in the 95% CI of the parametric bootstrap distribution of *τ_β_*, we considered it suggests there is consistency. By this criterion, 14 out of 23 loci have at least one multi-trait combination with consistent pleiotropic association pattern (Table 1). To visualize the comparison between consistent patterns and inconsistent ones, we looked into the 16 loci which are replicated by six-traits MVA (Fig 2). For example, both rs10761785 and rs9294260 can be replicated using six-traits MVA-significance replication. However, the 95% CI of *τ_β_* is [0.46,1] for rs10761785, and [−0.47,0.6] for rs9294260. This means the multi-trait marginal effects across samples are similar for rs10761785 but not for rs9294260. The contrast indicates that the underlying six-traits pleiotropic pattern is more plausible at the locus around rs10761785. For the locus around rs9294260, although it can be detected and replicated by MVA, the SNP may be only associated with a small subset of the six traits instead of most of them. To detect which traits tend to be irrelevant for a SNP, we can compare the discordance generated by each trait. For a SNP, if a trait introduces lots of inconsistency into the pleiotropic pattern, then the association between the SNP and that trait is more suspicious. For example, the marginal effect of height severely reduces the consistency for rs11231693, which indicates height is more likely to be irrelevant to rs11231693 among the six-traits. This detection is verified by its non-significance in height GWAS (p-value is 0.91 in GIANT2015 and 0.96 in UKB).

**Fig 2.**
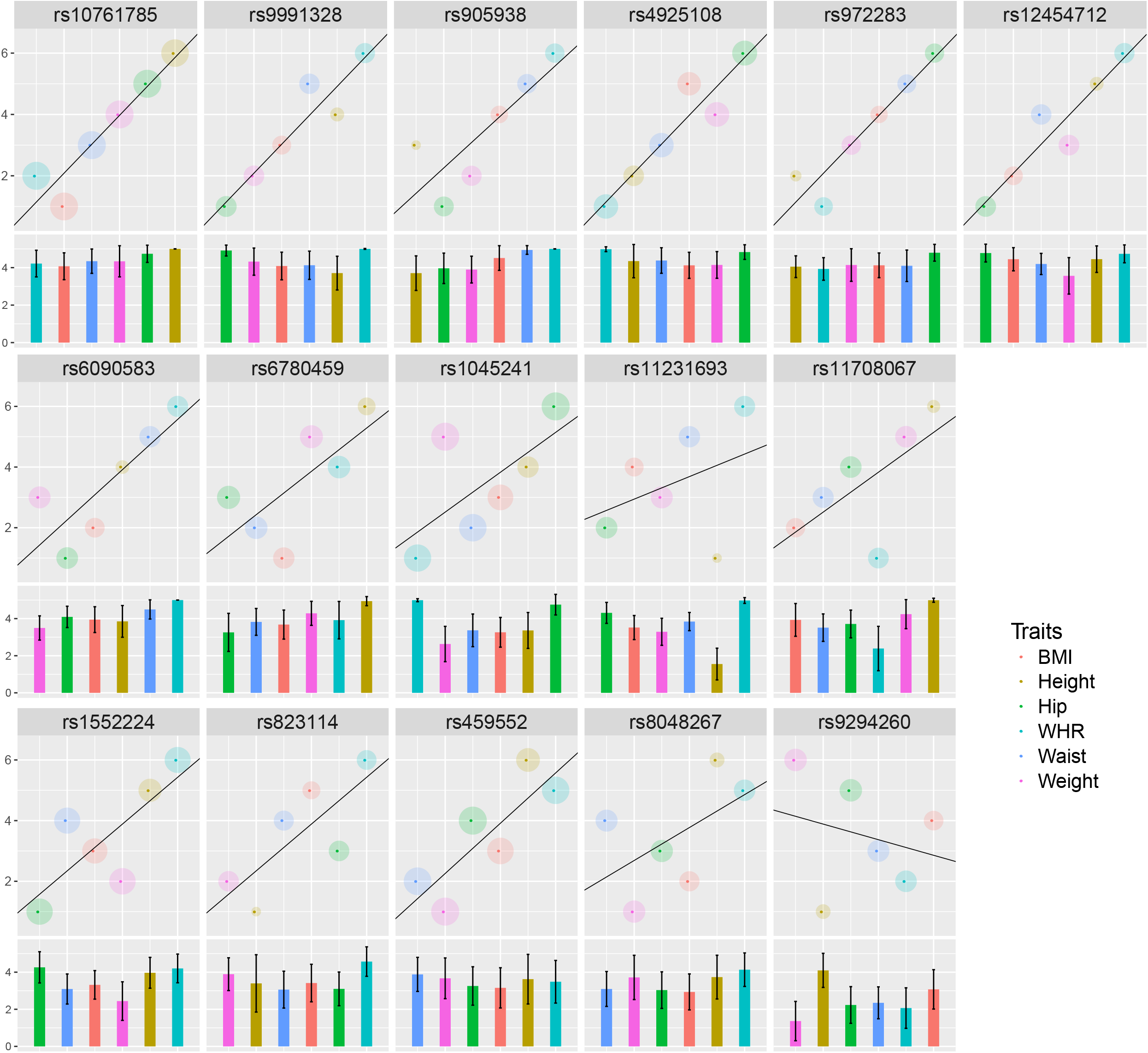
The correlations of the estimated marginal effects from GIANT and UKB at 16 loci which are replicated by six-traits MVA. The panels are reordered in ascending order according to the lower bounds of their 95% CI in correlation replication. The 11 loci in the first two rows are replicated by correlation replication. Each color represents one trait. There are two parts in each panel. In both parts, the x axis is the ranks of estimated marginal effect sizes in ascending order from GIANT. For the upper part, the y axis is the ranks from UKB. Therefore each dot represents the rank in GIANT and UKB for one trait. The radius of shade around a dot is proportional to the standard error of the estimated marginal effect. The standard errors are computed with variances in GIANT and UKB using inverse variance weights. To facilitate visualization, a regression line is added. Its slope equals to the Spearman’s correlation. The lower part shows the results based on 10,000 times Monte-Carlo simulations (described in the Materials and Methods). The y axis is the mean number of concordant pairs generated by a trait. If a trait has a very low bar, it means the trait disturbs the consistency. The whiskers represents ± 1 times the standard deviation about the mean.

Consequently, these two new replication strategies facilitate the investigation of pleiotropic architecture at each MVA-discovered locus, which is helpful for interpreting multi-trait association results.

### Conditional multivariate analysis suggests additional SNPs at new loci

Finally, as a secondary association analysis of MVA, we applied cMVA at the 23 new loci discovered and replicated in MVA. In this study, we took 1000 Genomes European-ancestry samples (*n* = 503) as reference sample to approximate regional LD matrix (described in the Materials and Methods). When all the six traits are analyzed together, cMVA identified 2 additional SNPs at these 23 novel loci with p-value lower than 5 × 10^−8^ in conditional multivariate analysis (Table 2). These two signals can both be replicated in UKB using either cMVA-significance replication (*p* < 0.05) or correlation replication (95% CI).

**Table 2.** MVA and cMVA results at two loci with additional hits suggested by cMVA.

| SNP | EA | $r$ | Single-SNP analysis | | Conditional analysis | | $\tau_\beta$ | CI |
| --- | --- | --- | --- | --- | --- | --- | --- | --- |
| | | | $p_G$ | $p_U$ | $p_G$ | $p_U$ | | |
| rs12033847 | T | 1 | 5.0E-10 | 2.1E-03 | 3.3E-10 | 1.2E-03 | *0.76 | (0.42, 0.88) |
| rs12138008 | T | 0.65 | 4.7E-08 | 5.2E-02 | 1.8E-09 | 3.1E-02 | - | - |
| rs4646404 | A | 1 | 1.1E-10 | 1.5E-07 | 1.7E-09 | 2.6E-05 | *0.64 | (0.21, 0.85) |
| rs7946 | T | 0.37 | 8.7E-08 | 2.7E-04 | 4.5E-08 | 2.8E-02 | - | - |

EA, effect allele; $r$ , LD correlation between a SNP and the top SNP at the locus in MVA; $p_G$ , cMVA p-value using summary statistics from GIANT; $p_U$ , cMVA p-value using individual-level data from UKB; $\tau_\beta$ , the observed correlation of multivariate marginal effects between GIANT and UKB, significant results are asterisked; CI, the parametric bootstrap distribution 95% confidence interval of $\tau_\beta$ .

## Discussion

We developed and implemented an analysis framework consisting of a series of methods to discover and replicate pleiotropic loci using GWAS summary statistics. Additionally, with a reference sample, we demonstrated how to perform conditional analysis to detect other traits-associated SNPs at loci discovered by MVA. We have shown that the analysis of multi-trait is not only powerful but also informative for evaluation of pleiotropy.

When individual-level data are available, our methods are equivalent to their correspondent individual-level versions (S2 Fig, S3 Fig). In practice, multi-trait analyses based on summary-level data usually have larger statistical power compared to those based on individual-level data for two reasons. Firstly, summary-level data from meta-anlyses have larger sample sizes in general. Secondly, when there are few individuals overlapping across traits, individual-level multivariate methods will lose power substantially by removing individuals with incomplete phenotypes. In the extreme case, when the samples are completely different for two traits, which means no individual gets both phenotypes measured, individual-level multivariate methods can not be implemented. Here, inspired by Zhu et al.’s work [8], we derive the detailed fundamental math for various test statistics when samples partially overlap. Therefore, methods in this study can account for the sample overlap properly by using summary statistics.

An essential issue of a multi-trait analysis is about deciding optimal trait sets, so that the multivariate test has larger power to capture known loci and discover new signals. The definition of the 6 sets used in this study depends on their relevance to body size or shape. If we compare the power of multi-trait and single-trait analyses under different pleiotropic architectures (S4 Fig), the power gain of the multi-trait analysis is determined by both the level of correlations among the phenotypes and by the effect directions of the genotype on the phenotypes. More specifically, power gain is achieved when the correlation between traits is positive and the genetic effects are opposite in their direction, or when the correlations between phenotypes is negative and genetic effect directions are same. Such scenarios may suggest an interesting biological basis of the phenotypes that can be missed in single-trait analysis.

We developed two statistical methods for replicating multi-trait signals using single-trait summary statistics. The replication results from these two methods can provide additional evidence for the existence of pleiotropy. The first method is MVA-score replication [12], which aims to replicate the association between SNP and the optimal linear combination of traits. This method is closely related to CCA (S2 Fig). If the association between MVA-score and a SNP is replicable, then the coefficients in the linear combination can be used to interpret the roles of traits in this association.

The second replication method is correlation replication, which evaluates the similarity of marginal effects across traits between samples by computing their Kendall’s correlation (S5 Fig). Since Kendall’s *τ* computes correlation using ranks of estimated marginal effects, any factor disturbing ranks weakens the correlation. For example, if a SNP does not have effects on all traits, then its estimated marginal effects on those irrelevant traits will be randomly ranked around zero, which reduces Kendall’s correlation (S6 Fig). An extreme case is when a SNP is only associated with one trait. In this case, there will be almost no consistency of the estimated effect sizes rank between samples. Therefore, if a SNP can be replicated by correlation replication, it has to be associated with more than one trait. On the other hand, when a SNP affects only a small subset of traits, it may not be replicated in correlation replication although pleiotropy exists. To solve this problem, we can compute the concordant pairs generated by each trait and identify traits weakening the correlation. Then a subset of traits could be taken and used to perform MVA and correlation replication again. Nevertheless, when the total number of traits is limited such as two or three, the correlation replication is less meaningful because the correlation is based on too few data points.

Although theoretically cMVA can be implemented on all the variants across the genome, we applied cMVA locus by locus for three reasons. Firstly, because SNPs separated by large genetic distance are usually independent to each other, regional conditional analysis is equivalent to genome-wise conditional analysis in most cases. The second reason is for accuracy. As in GCTA-COJO, we need a reference sample to approximate the LD correlations of the population where the meta-analysis sample is taken from. However, there is a mismatch between the LD structure of reference sample and that of meta-analysis sample. When more and more SNPs are selected, the mismatch accumulates and may disturb the results. To limit the error caused by the LD mismatch, we implemented cMVA locally so that not many SNPs are involved. The third reason is for power and computation. In some stepwise selection procedures including GCTA-COJO, the residuals from the regression given selected variants are used as new phenotype to search the next variant. This “adjusted-outcome” procedure is computationally fast because it performs univariate regression at each step. However, comparing to the standard multiple linear regression, where the original phenotype is the outcome and the selected variants are the covariates, the “adjusted-outcome” procedure has lower power when a new variant is correlated with the selected variants [23]. Since most of the additional traits-associated SNPs are in LD with the detected SNPs, we choose to use the standard multiple linear regression in cMVA. As a consequence, as more and more SNPs are selected, we have more and more covariates in the regression, which slows down the procedure. By implementing cMVA regionally, the analysis can be done quickly at each locus.

Based on GIANT2015 summary association statistics, our multi-trait analysis revealed 49 novel loci. Most of the replicated loci show pleiotropic effects beyond the six analyzed anthropometric traits. In particular, we see that some of newly discovered multivariate anthropometric loci is likely extending their effects onto metabolic (glucose and lipid levels) and life history (age at menarche, birth weight) phenotypes (S3 Appendix). According to results based on PhenoScanner [22], the loci with consistent pleiotropic effects on the six anthropometric traits are associated with more traits in general (S2 Appendix). As the MVA idetified loci are pleiotropic, a locus-specific test was also used to identify shared genetic basis between complex traits, including prediction of candidate genes according to expression quantitative trait loci (eQTL) analysis (S4 Appendix). This identifies, for some loci, that the pleiotropic effects are due to shared genetic causes instead of different linked causal variants.

Although only for a limited number of new loci, the existing mouse phenotyping database did suggest that the loci detected via multivariate analysis have functional relevance (S3 Appendix). Additional animal experimental validation is beyond the scope of this *in silico* paper. We suggest that future molecular studies should include such pleiotropic loci from multivariate analysis into investigation.

The developed pleiotropic analysis is implemented and freely available in the R package **MultiABEL** (The **GenABEL** project packages URL: https://r-forge.r-project.org/R/?group_id=505). The internal genome-wide screening module is implemented using Fortran 95 to gain computational speed.

With our results, we emphasize the value of combining multiple related phenotypes in large-scale genomic studies. We also emphasize the value of replication study for multi-trait analysis. Our results suggest that proper multivariate analysis may substantially enhance our understanding of shared genetic architecture between complex traits and disease and reveal more interesting biological knowledge.

## Supporting information

**S1 Appendix. Complete methods and derivations.**

**S2 Appendix. Pleiotropic effects of MVA loci.**

**S3 Appendix. Fine-mapping and candidate genes investigation.**

**S4 Appendix. Functional annotation via SMR-HEIDI.**

**S1 Fig. The number of loci replicated by different replication strategies.** Each circle in the Venn diagram represents one replication strategy. The four replication strategies are: single-trait GWAS replication (UVA), MVA-significance replication (MVA), MVA-score replication (Score) and correlation replication (Correlation).

**S2 Fig. Plot of coefficients from MVA-score against those from CCA.** We randomly sampled 50,000 individuals and 100 SNPs from UKB chromosome 22. Then for each SNP, we generated six marginal effects and traits. The estimated shrinkage phenotypic correlation matrix from GIANT was used as the phenotypic correlation matrix of simulated traits. Each SNP explains 0.01% variance of each trait. The x-axis represents the coefficients in reverse regression estimated from CCA based on individual-level data. The y-axis represents the coefficients estimated by MVA-score using summary statistics. In the left panel, true phenotypic correlation matrix was used in MVA-score. In the right panel, the phenotypic correlation matrix was estimated using the t-statistics from 10,000 simulated SNPs without effect. In both cases, MVA-score are almost equivalent to CCA.

**S3 Fig. Plot of −log_10_ p-values from cMVA against those from multiple regression based on individual-level data.** We randomly sampled 50,000 individuals from UKB and took their genotypes of 103 snps around SNP rs132622 as an example. Among the 103 snps, we randomly picked two as causal variants. Then the phenotypes were simulated as: *Y*_1_ = 0.3*X*_1_ + 0.1*X*_2_ + *ϵ*_1_, *Y*_2_ = 0.3*X*_1_ − 0.1*X*_2_ + *ϵ*_2_, where Var(*ϵ*_1_) = Var(*ϵ*_2_) = 2Cov(*ϵ*_1_,*ϵ*_2_) = 100. 40,000 individuals among 50,000 were used to perform GWAS and to generate summary statistics. A subset of the rest 10,000 individuals is used as reference sample to approximate LD matrix. The phenotypic correlation matrix in cMVA was estimated using the t-statistics from 10,000 simulated SNPs without effect. The reference sample used in cMVA is the original GWAS sample (left panel), an independent sample with 4,000 individuals (middle panel) or an independent sample with 500 individuals (right panel).

**S4 Fig. Power comparison of multi-trait and single-trait analyses.** Cor: correlation coefficient between each pair of phenotypes; *β_i_*: the genetic effect on the *i*-th phenotype; Half-overlap: each pair of the phenotypes share only 50% of the genotyped samples.

**S5 Fig. The rejection rate of correlation test under different scenarios.** In correlation test, the null hypothesis is rejected if the lower bound of MC-based CI is larger than 0. The lines represent the change of rejection rate when the percentile used for the lower bound is set to be different values. For example, percentile threshold = 0.05 means the test is based on whether 0 is above or below the 5th percentile of MC-based distribution for *τ_β_*. The first step is to simulate 12 traits in two samples. We firstly simulated 1,000 groups of marginal effects. In each group, 12 pairs of coefficients were drawn from 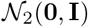, which are the marginal effects of a SNP on 12 traits in discovery and replication sample. Those groups with *τ_β_* = 0 or 0.15 or 0.3 were saved for next step. We then simulated a SNP for 10,000 individuals. The SNP explains 0.1% variance of each trait. After this, we sampled one group of coefficients from the saved groups and simulated phenotypes. The phenotypic correlation matrix of the 12 simulated traits is set as a block diagonal matrix, where the first 6 × 6 is the estimated shrinkage phenotypic correlation matrix from GIANT and the second 6 × 6 is the phenotypic correlation matrix from UKB. Then we performed the replication test and got the parametric bootstrap distribution of *τ_β_*. Each dot is based on 1,000 simulations. This figure shows that the test has no inflation or deflation when the true *τ_β_* = 0. The power of this test increases as the true *τ_β_* becomes larger.

**S6 Fig. The performance of correlation replication when zero effect sizes exist.** In this simulation, we set the marginal effects of a SNP on 12 traits in discovery and replication sample to be same. We firstly simulated 1,000 groups of marginal effects. In each group, 12 coefficients were drawn from 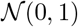, which are the marginal effects of a SNP on 12 traits in discovery and replication sample. Because the effect sizes for each trait are same across two samples, the true *τ_β_* = 1. To simulate the impact of zero effect sizes on the MC-based distribution of *τ_β_* in correlation test, we set the first several effect sizes as zero. In this case, the true *τ_β_* = 1 still, but the MC-based distribution of *τ_β_* would change. We then simulated a SNP for 10,000 individuals. The SNP explains 0.1% variance of each trait. The phenotypic correlation matrix of the 12 simulated traits is set as a block diagonal matrix, where the first 6 × 6 is the estimated shrinkage phenotypic correlation matrix from GIANT and the second 6 × 6 is the phenotypic correlation matrix from UKB. After this, we sampled one group of coefficients from the 1,0 groups and simulated phenotypes. Then we performed the replication test and got the parametric bootstrap distribution of *τ_β_*. The x-axis represents the number of traits on which the SNP has non-zero effect. The y-axis is the 5th percentile of the MC-based distribution of *τ_β_*.

**S7 Fig. The pleiotropic effects of six-traits MVA-only and UVA-only loci in GIANT2015 across different PhenoScanner p-value threshold.** (A) The x-axis represents p-value threshold in PhenoScanner; the y-axis is the natural logarithm of the number of associated traits plus one. (B) The parametric bootstrap distribution is computed for each locus based on the summary statistics of the six anthropometric traits in GIANT2015 and UKB. In each panel, the x-axis represents the lower bound of 95% CI of Kendall’s tau. The y-axis is the number of associated traits. To facilitate visualization, loci with the same lower bound values are clustered. At each lower bound value, only the median of the number of associated traits is plotted for each method.

The proportion of each cluster is represented by diameter of dots, which is computed separately for MVA and UVA. The curves are based on LOESS fit using data without clustering.

**S8 Fig. Effects of ELK4, ARAP1, and YDJC knock-out in mice across different phenotypes. The figure data were extracted from the International Mouse Phenotyping Consortium.**

**S9 Fig. ARAP1 and YDJC knock-out mice show difference in body mass and caudal vertebrae. The figure data were extracted from the International Mouse Phenotyping Consortium.**

**S10-S54 Fig. Visualization of the SMR-HEIDI test results for a single shared causal variant for different traits pairs.**

**S1 Table. MV analysis results for GIANT2013 data with replication on GIANT2015 data, UKB data, and GIANT2015+UKB meta-analysis data.**

**S2 Table. Extended results for 49 SNPs discovered using MV-only approach in GIANT2015 data.**

**S3 Table. Correlation matrix for GIANT 2013, GIANT2015 and UKB.**

**S4 Table. Pleiotropy database records with the anthropometric traits according to PhenoScanner (FDR < 5%) for the 49 novel loci.**

**S5 Table. SMR-HEIDI test results for the prediction of candidate gene and detection of shared genetic basis across complex traits.**

## Acknowledgments

We thank the global consortia, including GIANT, GLGC, MAGIC, CARDIOGRAM, DIAGRAM, CARDIOGRAMplusC4D, PGC, EGG, IBD, ADGC, RAGC, for making their reported summary-level data freely available online.

We thank the UK Biobank (Project No. 14302) for the data resource used in our replication analysis.

We thank the Swedish Twin Registry for allowing us extract the names of a set of independent SNPs to estimate the phenotypic shrinkage correlation coefficients.

## References

1. Visscher PM, Wray NR, Zhang Q, Sklar P, McCarthy MI, Brown MA, et al. 10 years of GWAS discovery: biology, function, and translation. The American Journal of Human Genetics. 2017;101(1):5–22.

2. Sivakumaran S, Agakov F, Theodoratou E, Prendergast JG, Zgaga L, Manolio T, et al. Abundant pleiotropy in human complex diseases and traits. The American Journal of Human Genetics. 2011;89(5):607–618.

3. Ferreira MA, Purcell SM. A multivariate test of association. Bioinformatics. 2008;25(1):132–133.

4. O’Reilly PF, Hoggart CJ, Pomyen Y, Calboli FC, Elliott P, Jarvelin MR, et al. MultiPhen: joint model of multiple phenotypes can increase discovery in GWAS. PloS one. 2012;7(5):e34861.

5. Aschard H, Vilhjálmsson BJ, Greliche N, Morange PE, Trégouët DA, Kraft P. Maximizing the power of principal-component analysis of correlated phenotypes in genome-wide association studies. The American Journal of Human Genetics. 2014;94(5):662–676.

6. Porter HF, O’Reilly PF. Multivariate simulation framework reveals performance of multi-trait GWAS methods. Scientific reports. 2017;7:38837.

7. Stephens M. A unified framework for association analysis with multiple related phenotypes. PloS one. 2013;8(7):e65245.

8. Zhu X, Feng T, Tayo BO, Liang J, Young JH, Franceschini N, et al. Meta-analysis of correlated traits via summary statistics from GWASs with an application in hypertension. American journal of human genetics. 2015;96(1):21–36.

9. Park H, Li X, Song YE, He KY, Zhu X. Multivariate analysis of anthropometric traits using summary statistics of genome-wide association studies from GIANT consortium. PloS one. 2016;11(10):e0163912.

10. Turley P, Walters RK, Maghzian O, Okbay A, Lee JJ, Fontana MA, et al. Multi-trait analysis of genome-wide association summary statistics using MTAG. Nature genetics. 2018; p. 1.

11. Cichonska A, Rousu J, Marttinen P, Kangas AJ, Soininen P, Lehtimäki T, et al. metaCCA: summary statistics-based multivariate meta-analysis of genome-wide association studies using canonical correlation analysis. Bioinformatics. 2016;32(13):1981–1989.

12. Shen X, Klarić L, Sharapov S, Mangino M, Ning Z, Wu D, et al. Multivariate discovery and replication of five novel loci associated with Immunoglobulin GN-glycosylation. Nature communications. 2017;8(1):447.

13. Yang J, Ferreira T, Morris AP, Medland SE, Genetic Investigation of ANthropometric Traits (GIANT) Consortium, DIAbetes Genetics Replication And Meta-analysis (DIAGRAM) Consortium, et al. Conditional and joint multiple-SNP analysis of GWAS summary statistics identifies additional variants influencing complex traits. Nature genetics. 2012;44(4):369–75–S1–3.

14. Deng Y, Pan W. Conditional analysis of multiple quantitative traits based on marginal GWAS summary statistics. Genetic epidemiology. 2017;41(5):427–436.

15. Zhu Z, Zheng Z, Zhang F, Wu Y, Trzaskowski M, Maier R, et al. Causal associations between risk factors and common diseases inferred from GWAS summary data. Nature communications. 2018;9(1):224.

16. Tsepilov Y, Sharapov S, Zaytseva O, Krumsek J, Prehn C, Adamski J, et al. A network-based conditional genetic association analysis of the human metabolome. GigaScience. 2018;7(12):giy137.

17. Randall JC, Winkler TW, Kutalik Z, Berndt SI, Jackson AU, Monda KL, et al. Sex-stratified genome-wide association studies including 270,000 individuals show sexual dimorphism in genetic loci for anthropometric traits. PLoS genetics. 2013;9(6):e1003500.

18. Wood AR, Esko T, Yang J, Vedantam S, Pers TH, Gustafsson S, et al. Defining the role of common variation in the genomic and biological architecture of adult human height. Nature genetics. 2014;46(11):1173–1186.

19. Shungin D, Winkler TW, Croteau-Chonka DC, Ferreira T, Locke AE, Mägi R, et al. New genetic loci link adipose and insulin biology to body fat distribution. Nature. 2015;518(7538):187.

20. Locke AE, Kahali B, Berndt SI, Justice AE, Pers TH, Day FR, et al. Genetic studies of body mass index yield new insights for obesity biology. Nature. 2015;518(7538):197–206.

21. Consortium GP, et al. A global reference for human genetic variation. Nature. 2015;526(7571):68.

22. Staley JR, Blackshaw J, Kamat MA, Ellis S, Surendran P, Sun BB, et al. PhenoScanner: a database of human genotype–phenotype associations. Bioinformatics. 2016;32(20):3207–3209.

23. Demissie S, Cupples LA. Bias due to two-stage residual-outcome regression analysis in genetic association studies. Genetic epidemiology. 2011;35(7):592–596.

