## Supplementary Information for "Beyond power: Multivariate discovery, replication, and interpretation of pleiotropic loci using summary association statistics"

### Supplemental Notes

#### Multi-trait association modeling and hypothesis testing

To simplify the formulae, we assume the phenotypes are standardized to have mean zero and variance one, and genotypes are centered to have mean zero. $k$ traits $Y_1, ...Y_k$ are dependent variables in MANOVA. Now we have $n$ individuals with these $k$ phenotypes $\mathbf{Y}_{n\times k}$ and a biallelic marker $\mathbf{g}_{n\times 1}$. Then the association between the group of $k$ phenotypes and the marker can be expressed as a multivariate regression

$$\mathbf{Y}_{n\times k} = \mathbf{g}_{n\times 1}\boldsymbol{\beta}'_{k\times 1} + \mathbf{e}_{n\times k}, \tag{1}$$

which can be tested via MANOVA for the null hypothesis

$$H_0 : \boldsymbol{\beta} = \mathbf{0}.$$

The estimates in the vector $\hat{\boldsymbol{\beta}}$ are known from GWA summary statistics. Below, we show how a MANOVA test statistic can be obtained without knowing the original data.

#### Calculating the multi-trait association test statistic

First of all, it is known that MANOVA test statistics, such as Pillai's trace, Wilk's lambda, etc., are all equivalent to an $F$ statistic for a single factor analysis[1], which is what we conduct in GWAS. When sample size is large, the $F$ test can be approximated by a $\chi^2$ test. Let $\mathbf{t} = [t_1, ..., t_k]'$ be the vector of single-trait t-test statistics across the $k$ phenotypes on the marker $\mathbf{g}$, and $\mathbf{R}^* \equiv \mathrm{Cor}(\mathbf{t}) = \mathrm{Var}(\mathbf{t})$. If $\mathbf{R}^*$ is available, the test statistic

$$T^2 = \mathbf{t}'\mathbf{R}^{*-1}\mathbf{t}, \tag{2}$$

which asymptotically follows a $\chi^2$ distribution with $k$ degrees of freedom under the null hypothesis.

Let $\mathbf{R}$ represent the phenotypic correlation matrix of the $k$ phenotypes. According to Zhu et al.[2], $\mathbf{R}^* = \mathbf{R}$ when the phenotypes are measured on the same set of individuals. When the samples partially overlap across different phenotypes, we derive the theory of shrinkage phenotypic correlation matrix below, which links $\mathbf{R}$ and $\mathbf{R}^*$. With the relationship between $\mathbf{R}$ and $\mathbf{R}^*$, $\mathbf{R}^*$ can be estimated using summary association statistics.

#### Shrinkage estimate of the phenotypic correlation matrix

Given a specific variant in GWAS, for trait $j$ and trait $j'$, let $\mathbf{g}$ be the genotypes of the overlapping $n_0$ individuals between the two traits. For the $n_1$ individuals with only trait $j$, we denote their genotypes as $\mathbf{x}$; and for the $n_2$ individuals with only trait $j'$, we denote their genotypes as $\mathbf{z}$. In another word, the genotypes are $[\mathbf{g}', \mathbf{x}']'$ for trait $j$, and $[\mathbf{g}', \mathbf{z}']'$ for trait $j'$. If the cohorts for trait $j$ and trait $j'$ are random samples of the same population, we can assume the variances of the variant are same and denote it as $\sigma_g^2$. Assuming $\bar{\mathbf{g}} = \bar{\mathbf{x}} = \bar{\mathbf{z}} = 0$, we have

$$\frac{\mathbf{g}'\mathbf{g}}{n_0} \approx \frac{\mathbf{x}'\mathbf{x}}{n_1} \approx \frac{\mathbf{z}'\mathbf{z}}{n_2} \approx \sigma_g^2 \tag{3}$$

Let $t_j = \hat{\beta}_j / \sqrt{\hat{\sigma}^2_{\hat{\beta}_j}}$ and $t_{j'} = \hat{\beta}_{j'} / \sqrt{\hat{\sigma}^2_{\hat{\beta}_{j'}}}$ be the test statistics of phenotypes $j$ and $j'$ against the variant, $\sigma_j^2$ and $\sigma_{j'}^2$ be the phenotypic variances, we have

$$t_j = \frac{\hat{\beta}_j}{\sqrt{\hat{\sigma}^2_{\hat{\beta}_j}}} = \frac{(\mathbf{g}'\mathbf{g} + \mathbf{x}'\mathbf{x})^{-1} [\mathbf{g}', \mathbf{x}'] \, \boldsymbol{y}_j}{\sqrt{\hat{\sigma}^2_{r_j} (\mathbf{g}'\mathbf{g} + \mathbf{x}'\mathbf{x})^{-1}}} \approx \frac{[\mathbf{g}', \mathbf{x}'] \, \boldsymbol{y}_j}{\sqrt{\sigma_j^2 \sigma_g^2 (n_0 + n_1)}},$$

where $\hat{\sigma}^2_{r_j}$ is the estimated residual variance in univariate regression. As the effect of a single variant is usually small, we can approximate $\hat{\sigma}^2_{r_j}$ by $\sigma^2_j$. Similarly,

$$t_{j'} \approx \frac{[\mathbf{g}', \mathbf{z}']\, \boldsymbol{y}_{j'}}{\sqrt{\sigma^2_{j'}\sigma^2_g(n_0+n_2)}}$$

Therefore,

$$\begin{aligned}
R^*_{j,j'} &= \mathrm{Cor}(t_j, t_{j'}) = \mathrm{Cov}(t_j, t_{j'}) && (4)\\
&\approx \mathrm{Cov}\left(\frac{[\mathbf{g}', \mathbf{x}']\, \boldsymbol{y}_j}{\sqrt{\sigma^2_j\sigma^2_g(n_0+n_1)}}, \frac{[\mathbf{g}', \mathbf{z}']\, \boldsymbol{y}_{j'}}{\sqrt{\sigma^2_{j'}\sigma^2_g(n_0+n_2)}}\right)\\
&= \frac{[\mathbf{g}', \mathbf{x}']\, \mathrm{Cov}(\boldsymbol{y}_j, \boldsymbol{y}_{j'})\, [\mathbf{g}', \mathbf{z}']'}{\sqrt{\sigma^2_j\sigma^2_{j'}\sigma^4_g(n_0+n_1)(n_0+n_2)}}\\
&= \frac{R_{j,j'}\sigma_j\sigma_{j'}\, [\mathbf{g}', \mathbf{x}'] \begin{bmatrix} \mathbf{I} & \\ & \mathbf{0} \end{bmatrix} \begin{bmatrix} \mathbf{g} \\ \mathbf{z} \end{bmatrix}}{\sqrt{\sigma^2_j\sigma^2_{j'}\sigma^4_x(n_0+n_1)(n_0+n_2)}}\\
&= \frac{R_{j,j'}\mathbf{g}'\mathbf{g}}{\sqrt{\sigma^4_g(n_0+n_1)(n_0+n_2)}}\\
&\approx \frac{n_0}{\sqrt{(n_0+n_1)(n_0+n_2)}} R_{j,j'}, && (5)
\end{aligned}$$

Therefore, the correlation of t-statistics is a shrinkage version of the phenotypic correlation, with a factor determined by the level of overlap. When no overlap individual exists across the phenotypes, the test statistic automatically reduces to Fisher's method of $\chi^2$ accumulation.

#### Conditional multivariate analysis

When cMVA is performed and $p$ SNPs $G = (G_1, ..., G_p)$ are involved, we let $\mathbf{G}_{n\times p}$ denote the genotype matrix of these $p$ SNPs with sample size of $n$. Then (1) can be extended to:

$$\boldsymbol{y}_{n\times k} = \mathbf{G}_{n\times p}\boldsymbol{\beta}'_{k\times p} + \mathbf{e}_{n\times k}. \quad (6)$$

The effects of SNP $i$ conditional on the other $p-1$ SNPs can be tested using

$$H_0 : \tilde{\boldsymbol{\beta}}_i = \mathbf{0}. \tag{7}$$

where $\tilde{\boldsymbol{\beta}}_i$ represents the conditional effects of SNP $i$ on the $k$ traits.

Similar to (2), above hypothesis can be tested using conditional t-test statistics from (6). For SNP $i$, if the conditional t-test statistics $\tilde{\mathbf{t}}_i = [\tilde{t}_{i1}, ..., \tilde{t}_{ik}]'$ in (6) and their correlation matrix $\tilde{\mathbf{R}}_i^*$ are available, then we can obtain the test statistic

$$\tilde{T}_i^2 = \tilde{\mathbf{t}}_i' \tilde{\mathbf{R}}_i^{*-1} \tilde{\mathbf{t}}_i, \tag{8}$$

which also asymptotically follows a $\chi^2$ distribution with $k$ degrees of freedom under (7).

Next, we will show how to get $\tilde{\mathbf{t}}_i$ and $\tilde{\mathbf{R}}_i^*$. As shown in literature[3,4], the joint regression results asymptotically only depends on: (i) the covariance structure between variants and traits; (ii) the LD structure between variants. Therefore we can approximate joint regression results including $\tilde{\mathbf{t}}_i$ using summary-level statistics from GWAS meta-analyses and a reference sample. In the following sections, we will use $\mathbf{A}_{i\cdot}$ and $\mathbf{A}_{\cdot j}$ to represent the $i$th row and the $j$th column of matrix $\mathbf{A}$ respectively, and $\mathbf{A}'_{i\cdot}$ represents $(\mathbf{A}_{i\cdot})'$. For trait $j$, in single trait GWAS we have

$$\begin{aligned} \hat{b}_{ij} &= (\mathbf{G}'_{\cdot i}\mathbf{G}_{\cdot i})^{-1}\mathbf{G}'_{\cdot i}\boldsymbol{y}_j \approx \frac{\mathrm{Cov}(G_i, Y_j)}{\sigma^2_{g_i}} \\ \hat{\sigma}^2_{\hat{b}_{ij}} &= \sigma^2_{r,ij}(\mathbf{G}'_{\cdot i}\mathbf{G}_{\cdot i})^{-1} \approx \frac{\sigma^2_j}{n\sigma^2_{g_i}}, \end{aligned}$$

where $\hat{b}_{ij}$ and $\hat{\sigma}^2_{\hat{b}_{ij}}$ are the estimated marginal effect of variant $i$ on trait $j$ and its variance, $\sigma^2_{r,ij}$ is the residual variance and can be approximated by phenotypic variance $\sigma^2_j$, and $\sigma^2_{g_i} = \mathrm{Var}(G_i)$.

The LD structure between SNPs can be approximated by a representative reference sample where individual-level genotype data are available[3]. Let $\mathbf{W}$ represents the $n_W \times p$ genotypes of the reference sample, then

$$\frac{\mathbf{G}'\mathbf{G}}{n} \approx \frac{\mathbf{W}'\mathbf{W}}{n_W} \approx \mathrm{Var}(G).$$

Then the conditional effect of variant $i$ on trait $j$

$$\begin{aligned}
\hat{\beta}^c_{ij} &= \left[(\mathbf{G}'\mathbf{G})^{-1}\mathbf{G}'\boldsymbol{y}_j\right]_i \\
&\approx \left[\mathrm{Var}^{-1}(G)\mathrm{Cov}(G, Y_j)\right]_i \\
&= \left[\mathrm{Var}^{-1}(G)\right]_{i\cdot} \begin{bmatrix} \sigma^2_{g_1}\hat{b}_{1j} \\ \vdots \\ \sigma^2_{g_p}\hat{b}_{pj} \end{bmatrix} \\
&= \frac{1}{\sigma_{g_i}} \left[\mathrm{Cor}^{-1}(G)\right]_{i\cdot} \begin{bmatrix} \sigma^{-1}_{g_1} & & \\ & \ddots & \\ & & \sigma^{-1}_{g_p} \end{bmatrix} \begin{bmatrix} \sigma^2_{g_1}\hat{b}_{1j} \\ \vdots \\ \sigma^2_{g_p}\hat{b}_{pj} \end{bmatrix} \\
&= \frac{1}{\sigma_{g_i}} \left[\mathrm{Cor}^{-1}(G)\right]_{i\cdot} \begin{bmatrix} \sigma_{g_1}\hat{b}_{1j} \\ \vdots \\ \sigma_{g_p}\hat{b}_{pj} \end{bmatrix},
\end{aligned}$$

and its variance

$$\hat{\sigma}^2_{\hat{\beta}^c_{ij}} = \left[\sigma^2_{r,j}(\mathbf{G}'\mathbf{G})^{-1}\right]_{ii} \approx \frac{\sigma^2_j}{n}\left[\mathrm{Var}^{-1}(G)\right]_{ii} = \frac{\sigma^2_j}{n\sigma^2_{g_i}}\left[\mathrm{Cor}^{-1}(G)\right]_{ii},$$

so that the conditional t-statistic

$$\begin{aligned}
\tilde{t}_{ij} &= \frac{\hat{\beta}^c_{ij}}{\sqrt{\hat{\sigma}^2_{\hat{\beta}^c_{ij}}}} \\
&= \frac{\left[\mathrm{Cor}^{-1}(G)\right]_{i\cdot}}{\left[\mathrm{Cor}^{-1}(G)\right]_{ii}} \begin{bmatrix} \frac{\sqrt{n}\sigma_{g_1}\hat{b}_{1j}}{\sigma_j} \\ \vdots \\ \frac{\sqrt{n}\sigma_{g_p}\hat{b}_{pj}}{\sigma_j} \end{bmatrix} \\
&= \frac{\left[\mathrm{Cor}^{-1}(G)\right]_{i\cdot}}{\left[\mathrm{Cor}^{-1}(G)\right]_{ii}} \begin{bmatrix} \frac{\hat{b}_{1j}}{\hat{\sigma}_{\hat{b}_{1j}}} \\ \vdots \\ \frac{\hat{b}_{pj}}{\hat{\sigma}_{\hat{b}_{pj}}} \end{bmatrix} \\
&= \frac{\left[\mathrm{Cor}^{-1}(G)\right]_{i\cdot}}{\left[\mathrm{Cor}^{-1}(G)\right]_{ii}} \begin{bmatrix} t_{1j} \\ \vdots \\ t_{pj} \end{bmatrix}.
\end{aligned}$$

In another word, the conditional t-statistics of SNP $i$ are linear combinations of the marginal t-statistics of all the SNPs.

Now we focus on $\tilde{\mathbf{R}}^*_i$, which is the correlation matrix of the conditional t-statistics for SNP $i$. The following derivation is an extension of (4) to (5). Here we use $\mathbf{G}_{n_0 \times p}$, $\mathbf{X}_{n_1 \times p}$ and $\mathbf{Z}_{n_2 \times p}$ to represent the genotypes of shared individuals, trait $j$ only individuals and trait $j'$ only individuals. Then the genotypes for trait $j$ are $[\mathbf{G}', \mathbf{X}']'$, and those for trait $j'$ are $[\mathbf{G}', \mathbf{Z}']'$. Similar to (3), we have

$$\frac{\mathbf{G}'\mathbf{G}}{n_0} \approx \frac{\mathbf{X}'\mathbf{X}}{n_1} \approx \frac{\mathbf{Z}'\mathbf{Z}}{n_2} \approx \mathrm{Var}(G) \tag{9}$$

To simplify symbols, we define $\mathbf{A}_{p\times p} = \mathrm{Var}^{-1}(G)$, $\mathbf{B}_{p\times p} = (\mathbf{G}'\mathbf{G} + \mathbf{X}'\mathbf{X})^{-1}$ and $\mathbf{C}_{p\times p} = (\mathbf{G}'\mathbf{G} + \mathbf{Z}'\mathbf{Z})^{-1}$. Then according to (9),

$$\mathbf{B} \approx \frac{1}{n_0 + n_1}\mathbf{A}, \quad \text{and} \quad \mathbf{C} \approx \frac{1}{n_0 + n_2}\mathbf{A}.$$

Therefore for a pair of traits $j$ and $j'$,

$$\begin{aligned}
\tilde{R}^*_{i,jj'} &= \text{Cor}(\tilde{t}_{ij}, \tilde{t}_{ij'}) = \text{Cov}(\tilde{t}_{ij}, \tilde{t}_{ij'}) \qquad (10)\\
&\approx \text{Cov}\left( \frac{\mathbf{B}_{i\cdot}\,[\mathbf{G}', \mathbf{X}']\,\boldsymbol{y}_j}{\sqrt{\sigma_j^2 \mathbf{B}_{ii}}}, \frac{\mathbf{C}_{i\cdot}\,[\mathbf{G}', \mathbf{Z}']\,\boldsymbol{y}_{j'}}{\sqrt{\sigma_{j'}^2 \mathbf{C}_{ii}}} \right)\\
&= \frac{\mathbf{B}_{i\cdot}\,[\mathbf{G}', \mathbf{X}']\,\text{Cov}(\boldsymbol{y}_j, \boldsymbol{y}_{j'})\,[\mathbf{G}', \mathbf{Z}']'\,\mathbf{C}'_{i\cdot}}{\sqrt{\sigma_j^2 \sigma_{j'}^2 \mathbf{B}_{ii}\mathbf{C}_{ii}}}\\
&= \frac{R_{j,j'}\sigma_j\sigma_{j'}\mathbf{B}_{i\cdot}\,[\mathbf{G}', \mathbf{X}'] \begin{bmatrix} \mathbf{I} & \\ & \mathbf{0} \end{bmatrix} \begin{bmatrix} \mathbf{G} \\ \mathbf{Z} \end{bmatrix} \mathbf{C}'_{i\cdot}}{\sqrt{\sigma_j^2 \sigma_{j'}^2 \mathbf{B}_{ii}\mathbf{C}_{ii}}}\\
&= \frac{R_{j,j'}\mathbf{B}_{i\cdot}(\mathbf{G}'\mathbf{G})\mathbf{C}'_{i\cdot}}{\sqrt{\mathbf{B}_{ii}\mathbf{C}_{ii}}}\\
&\approx \frac{n_0 R_{j,j'}}{\sqrt{(n_0+n_1)(n_0+n_2)}} \cdot \frac{\mathbf{A}_{i\cdot}\mathbf{A}^{-1}\mathbf{A}'_{i\cdot}}{\mathbf{A}_{ii}}\\
&= \frac{n_0}{\sqrt{(n_0+n_1)(n_0+n_2)}} R_{j,j'}. \qquad (11)
\end{aligned}$$

Above derivation shows that, for all SNPs, the conditional t-statistics correlation equals to the shrinkage phenotypic correlation. This means our previous $\hat{\mathbf{R}}^*$ estimated from correlating GWAS t-statistics can be used as $\tilde{\mathbf{R}}^*_i$ for any SNP in cMVA.

#### Correlation replication approach

For SNPs discovered in MVA or cMVA, to replicate their potential pleiotropic pattern in the replication sample, we develop a MC-based correlation replication approach. Following the $k$ traits, $p$ SNPs model in (6), we repeatedly drew $\boldsymbol{\beta}^{MC}$ from $p \times k$ variate normal distribution $\mathcal{N}(\hat{\boldsymbol{\beta}}^c, \hat{\boldsymbol{\Sigma}})$, where $\hat{\boldsymbol{\beta}}^c = (\hat{\beta}^c_{11}, ...\hat{\beta}^c_{p1}, \hat{\beta}^c_{12}, ..., \hat{\beta}^c_{p2}, ..., \hat{\beta}^c_{1k}, ..., \hat{\beta}^c_{pk})$ is the vector of estimated conditional effects, and $\hat{\boldsymbol{\Sigma}}$ is an estimate of $\boldsymbol{\Sigma} = \text{Cov}(\hat{\boldsymbol{\beta}}^c)$. Because $\hat{\boldsymbol{\beta}}^c$ and the variances of $\{\hat{\beta}^c_{ij}\}$ can be obtained from cMVA, we only need to estimate the elements of $\mathrm{Cor}(\hat{\boldsymbol{\beta}}^c)$. Similar to (10)-(11),

$$\begin{aligned}
\mathrm{Cor}(\hat{\beta}^c_{ij}, \hat{\beta}^c_{i'j'}) &= \mathrm{Cor}(\tilde{t}_{ij}, \tilde{t}_{i'j'}) = \mathrm{Cov}(\tilde{t}_{ij}, \tilde{t}_{i'j'}) \\
&\approx \mathrm{Cov}\left( \frac{\mathbf{B}_{i\cdot}\,[\mathbf{G}', \mathbf{X}']\,\boldsymbol{y}_j}{\sqrt{\sigma_j^2 \mathbf{B}_{ii}}}, \frac{\mathbf{C}_{i'\cdot}\,[\mathbf{G}', \mathbf{Z}']\,\boldsymbol{y}_{j'}}{\sqrt{\sigma_{j'}^2 \mathbf{C}_{i'i'}}} \right) \\
&= \frac{\mathbf{B}_{i\cdot}\,[\mathbf{G}', \mathbf{X}']\,\mathrm{Cov}(\boldsymbol{y}_j, \boldsymbol{y}_{j'})\,[\mathbf{G}', \mathbf{Z}']'\,\mathbf{C}'_{i'\cdot}}{\sqrt{\sigma_j^2 \sigma_{j'}^2 \mathbf{B}_{ii} \mathbf{C}_{i'i'}}} \\
&= \frac{R_{j,j'}\sigma_j\sigma_{j'}\mathbf{B}_{i\cdot}\,[\mathbf{G}', \mathbf{X}'] \begin{bmatrix} \mathbf{I} & \\ & \mathbf{0} \end{bmatrix} \begin{bmatrix} \mathbf{G} \\ \mathbf{Z} \end{bmatrix} \mathbf{C}'_{i'\cdot}}{\sqrt{\sigma_j^2 \sigma_{j'}^2 \mathbf{B}_{ii} \mathbf{C}_{i'i'}}} \\
&= \frac{R_{j,j'}\mathbf{B}_{i\cdot}(\mathbf{G}'\mathbf{G})\mathbf{C}'_{i'\cdot}}{\sqrt{\mathbf{B}_{ii}\mathbf{C}_{i'i'}}} \\
&= \frac{n_0 R_{j,j'}}{\sqrt{(n_0+n_1)(n_0+n_2)}} \cdot \frac{\mathbf{A}_{i\cdot}\mathbf{A}^{-1}\mathbf{A}'_{i'\cdot}}{\sqrt{\mathbf{A}_{ii}\mathbf{A}_{i'i'}}} \\
&\approx \frac{n_0 R_{j,j'}}{\sqrt{(n_0+n_1)(n_0+n_2)}} \cdot \frac{\mathbf{A}_{ii'}}{\sqrt{\mathbf{A}_{ii}\mathbf{A}_{i'i'}}},
\end{aligned}$$

which means $\mathrm{Cor}(\hat{\boldsymbol{\beta}}^c) \approx \hat{\mathbf{R}}^* \otimes \mathrm{Cor}^{-1}(G)$, where $\otimes$ represents Kronecker product. Specifically, for MVA where $p = 1$, $\mathrm{Cor}(\hat{\boldsymbol{\beta}}^c) \approx \hat{\mathbf{R}}^*$.

With $\hat{\boldsymbol{\beta}}^c$ and $\hat{\boldsymbol{\Sigma}}$ for the discovery sample and the replication sample, we can draw $\boldsymbol{\beta}^{MC}_{disc}$ and $\boldsymbol{\beta}^{MC}_{rep}$, then compute their Kendall's rank correlation coefficient

$$\hat{\tau}_\beta = \frac{2}{k(k-1)} \sum_{j<j'} \mathrm{sgn}(\beta^{MC}_{j,disc} - \beta^{MC}_{j',disc}) \cdot \mathrm{sgn}(\beta^{MC}_{j,rep} - \beta^{MC}_{j',rep}).$$

After repeating the parametric sampling many times (10,000 times in this study), we can get an estimated distribution of $\tau_\beta$. The parametric bootstrap confidence intervals (CI) based on this distribution can be used for inference. In this study, we reject the null hypothesis $H_0 : \tau_\beta = 0$ if the lower bound of the 95% CI of parametric bootstrap distribution is larger than 0.

#### Constructing the new phenotype score

We construct a new phenotype score as a linear combination of the original six phenotypes, via the following multiple regression model,

$$\mathbf{g} = \mathbf{Y}\mathbf{b} + \boldsymbol{\epsilon}, \tag{12}$$

which is equivalent to CCA.

Now we show how the coefficients estimates in (12) can also be obtained without knowing the original data. First of all, we derive the coefficients estimates of each *swapped* GWA simple regression model,

$$\mathbf{g} = \mathbf{y}_j b_j^* + \boldsymbol{\epsilon}_j^*$$

From the summary statistics, we know the estimates of the following GWA simple regression model,

$$\mathbf{y}_j = \mathbf{g}\beta_j + \mathbf{e}_j$$

i.e.

$$\hat{\beta}_j = \frac{\mathbf{g}'\mathbf{y}_j}{\mathbf{g}'\mathbf{g}} = \frac{\mathbf{g}'\mathbf{y}_j}{2nf(1-f)}$$

assuming Hardy-Weinberg equilibrium (HWE), where $f$ is the coding allele frequency of the marker. We also have

$$\hat{b}_j^* = \frac{\mathbf{y}_j'\mathbf{g}}{\mathbf{y}_j'\mathbf{y}_j} = \frac{\mathbf{y}_j'\mathbf{g}}{n}$$

As $\mathbf{g}'\mathbf{y}_j = \mathbf{y}_j'\mathbf{g}$, we have

$$\hat{b}_j^* = 2f(1-f)\hat{\beta}_j$$

Thereafter, when $\text{Var}(y_j) = 1$ for all $j$, the estimates $\hat{\mathbf{b}}$ in (12) can be calculated as

$$\begin{aligned}\hat{\mathbf{b}} &= (\mathbf{Y}'\mathbf{Y})^{-1}\mathbf{D}\hat{\mathbf{b}}^* \\ &= \mathbf{R}^{-1}\hat{\mathbf{b}}^*\end{aligned}$$

where $\mathbf{D}$ is a diagonal matrix with the $j$-th element $\mathbf{y}_j'\mathbf{y}_j$. So that a new phenotype score can be defined as

$$\mathbf{S} = \mathbf{Y}\hat{\mathbf{b}}$$

The variance-covariance matrix of $\hat{\mathbf{b}}$ is

$$V(\hat{\mathbf{b}}) = \sigma_b^2(\mathbf{Y}'\mathbf{Y})^{-1}$$

where

$$\sigma_b^2 = \frac{\mathbf{g}'\mathbf{g} - \hat{\mathbf{b}}'\mathbf{D}\hat{\mathbf{b}}^*}{n-k} = \frac{2nf(1-f) - \hat{\mathbf{b}}'\mathbf{D}\hat{\mathbf{b}}^*}{n-k}$$

The standard errors of $\hat{\mathbf{b}}$ can be obtained as the square roots of the diagonal elements of $V(\hat{\mathbf{b}})$, which can be used for testing individual-phenotype associations with the SNP, corrected for all the other phenotypes as covariates.

#### Estimating the genetic effect on the new phenotype score

In a replication cohort, the genetic effect of each SNP on the new phenotype score $\mathbf{S}$ can be tested via simple regression of $\mathbf{S}$ on the allelic dosages $\mathbf{g}$,

$$\mathbf{S} = \mathbf{g}\beta_s + \mathbf{e}_s. \tag{13}$$

Interestingly, without knowing the original data, we can obtain the estimate of $\beta_s$ in the discovery sample using summary statistics. The following proof shows that $\hat{\beta}_s$ always equals to Pillai's trace $V$ in (1). According to (12), we have

$$\hat{\mathbf{b}} = (\mathbf{Y}'\mathbf{Y})^{-1}\mathbf{Y}'\mathbf{g},$$

so that in (13),

$$\hat{\beta}_s = \frac{\mathbf{S}'\mathbf{g}}{\mathbf{g}'\mathbf{g}} = \frac{\hat{\mathbf{b}}'\mathbf{Y}'\mathbf{g}}{\mathbf{g}'\mathbf{g}} = \frac{\mathbf{g}'\mathbf{Y}(\mathbf{Y}'\mathbf{Y})^{-1}\mathbf{Y}'\mathbf{g}}{\mathbf{g}'\mathbf{g}}. \tag{14}$$

In (1), by definition, Pillai's trace $V = \mathrm{tr}\{(\mathbf{T}-\mathbf{E})\,\mathbf{T}^{-1}\}$, where

$$\begin{aligned} \mathbf{E} &= \mathbf{Y}'\left\{I - \mathbf{g}\left(\mathbf{g}'\mathbf{g}\right)^{-1}\mathbf{g}'\right\}\mathbf{Y}, \\ \mathbf{T} &= \mathbf{Y}'\mathbf{Y}. \end{aligned} \tag{15}$$

Hence

$$\mathbf{T} - \mathbf{E} = \mathbf{Y}'\mathbf{g}\left(\mathbf{g}'\mathbf{g}\right)^{-1}\mathbf{g}'\mathbf{Y}. \tag{16}$$

Combining with (15) and (16), we get

$$\begin{aligned} V &= \mathrm{tr}\{(\mathbf{T}-\mathbf{E})\,\mathbf{T}^{-1}\} \\ &= \mathrm{tr}\left\{\frac{(\mathbf{Y}'\mathbf{g})(\mathbf{g}'\mathbf{Y})(\mathbf{Y}'\mathbf{Y})^{-1}}{\mathbf{g}'\mathbf{g}}\right\} \\ &= \mathrm{tr}\left\{\frac{(\mathbf{g}'\mathbf{Y})(\mathbf{Y}'\mathbf{Y})^{-1}(\mathbf{Y}'\mathbf{g})}{\mathbf{g}'\mathbf{g}}\right\} \\ &= \hat{\beta}_s. \end{aligned}$$

To get the standard error of $\hat{\beta}_s$, we should notice that given the numerical equivalence of $V$ and $\hat{\beta}_s$, $\hat{\beta}_s$ has the same distribution as $V$. Denote $F$ as the F-test statistic in (1), $V$ can be transformed as

$$F = \frac{V^2/k}{(1-V^2)/(n-k-1)}.$$

After rearranging,

$$\hat{\beta}_s = V = \frac{kF}{(n-k-1)+kF}.$$

As $F \sim F(k, n-k-1)$, we have

$$\hat{\beta}_s = V \sim Beta\left(\frac{k}{2}, \frac{n-k-1}{2}\right),$$

which is the exact distribution of $\hat{\beta}_s$. In practice, the standard error of $\hat{\beta}_s$ can be obtained by Gaussian approximation of the Beta distribution. If we translate the MANOVA p-value back to a 1 d.f. $\chi^2$ statistic $C$, we have

$$\frac{\hat{\beta}_s^2}{\mathrm{Var}(\hat{\beta}_s)} = \frac{V^2}{\mathrm{Var}(\hat{\beta}_s)} \approx C.$$

Then we can compute the standard error of $\hat{\beta}_s$ in the meta-GWAS population as $VC^{-1/2}$.

Also, if we denote $R^2$ as the coefficient of determination of both regressions (12) and (13) in the discovery sample, we have

$$
\begin{aligned}
R^2 &= \frac{\hat{\beta}_s' \mathbf{g}' \mathbf{g} \hat{\beta}_s}{\mathbf{s}'\mathbf{s}} = \frac{\hat{\beta}_s^2 \mathbf{g}'\mathbf{g}}{\hat{\mathbf{b}}'\mathbf{Y}'\mathbf{Y}\hat{\mathbf{b}}} \\
&= \frac{\hat{\beta}_s^2 \mathbf{g}'\mathbf{g}}{\mathbf{g}'\mathbf{Y}(\mathbf{Y}'\mathbf{Y})^{-1}\mathbf{Y}'\mathbf{Y}(\mathbf{Y}'\mathbf{Y})^{-1}\mathbf{Y}'\mathbf{g}} \\
&= \frac{\hat{\beta}_s^2 \mathbf{g}'\mathbf{g}}{\mathbf{g}'\mathbf{Y}(\mathbf{Y}'\mathbf{Y})^{-1}\mathbf{Y}'\mathbf{g}} \\
&= \frac{\hat{\beta}_s^2}{\left\{\mathbf{g}'\mathbf{Y}(\mathbf{Y}'\mathbf{Y})^{-1}\mathbf{Y}'\mathbf{g}\right\}/(\mathbf{g}'\mathbf{g})} \\
&= \frac{\hat{\beta}_s^2}{\hat{\beta}_s} \\
&= \hat{\beta}_s
\end{aligned}
$$

Therefore, Pillai's trace $V$ directly represents the proportion of the variance of $\mathbf{S}$ explained by the SNP.

#### Pleiotropic effects of MVA loci

Intuitively, comparing to loci detected by single-trait GWAS, MVA loci are potentially associated with more traits beyond the analyzed set of traits. We performed a comparison between MVA and UVA loci in GIANT2015. For the six anthropometric traits, at a p-value threshold of $5 \times 10^{-8}$, there are 456 six-traits MVA loci and 519 UVA loci. After excluding 425 loci which are significant in both MVA and UVA, we compared the 31 loci only detected by six-traits MVA and the 94 loci only detected by UVA. We examined the pleiotropic effects of these loci by extracting GWAS database records of the top SNPs from PhenoScanner[5]. In PhenoScanner, we set the p-value threshold ranged from $5 \times 10^{-8}$ to 0.05 so that the associations with various complex traits were taken into account. According to our results, even when all of the associations with anthropometric traits were excluded from the database records, MVA loci are still associated with significantly more other traits than UVA loci, regardless of the p-value thresholds (**Figure S7.A**). Additionally, we computed the parametric bootstrap distribution of $\tau_\beta$ for these MVA-only loci and UVA-only loci. We found for both MVA-only loci and UVA-only loci, the number of associated traits increases with the lower bound of $\tau_\beta$ CI, especially for MVA loci (**Figure S7.B**). It is also interesting to notice that all the six loci with the highest lower bound (0.46) are MVA-only loci.

We then extracted the PhenoScanner records of the 49 novel loci discovered in GIANT2015. Filtered at a false discovery rate (FDR) threshold of 5%, there were 427 significant associations with different classes of phenotypes, including gene expression, DNA methylation, metabolomics, blood lipids, and complex disease etc. (**Fig. 1**, **Table S4**).

#### Functional annotation via SMR-HEIDI

We have looked up each SNP of the newly mapped anthropometric loci using the PhenoScanner database, to check whether it was previously reported to be associated with other complex traits with p-value lower than $5 \times 10^{-8}$ and proxy $r^2 = 0.7$.

To test for potential pleiotropic effects of identified variants we have used the following procedure. Starting with a significant index SNP, we have screened for the traits which may be affected by variation in the same region, and then performed pleiotropy vs. linkage disequilibrium test (HEIDI test[6]). In the screening stage, we have used Phenoscanner to look up complex traits that have demonstrated $p < 5 \times 10^{-8}$ for association with the index SNP (or with a SNP having $r^2 > 0.7$ with the index SNP). Note that to perform HEIDI testing one needs regional summary level GWAS results, including signed regression coefficient estimates and standard error of these estimates. Therefore for complex traits identified, we attempted to collect GWAS summary statistics to perform HEIDI test (own implementation). When summary-level GWAS data (which should minimally include effective allele, p-value for association, and regression coefficient estimate) were available, we have verified whether $p < 5 \times 10^{-8}$ was indeed observed for the index SNP (or one of the strongly associated 'proxy'

SNPs in case the index SNP was missing in the secondary GWAS) before performing HEIDI test. We have also looked up for overlap between our results and blood[7] eQTLs, using similar procedure with modifications. For blood eQTLs[7], we did not use *phenoscanner* in the screening phase, but rather tested directly if association results for specific probe are reported in the region of interest and, if positive, tested whether index/proxy SNP had $p < 5 \times 10^{-8}$ in eQTL analysis; if positive, the HEIDI test was performed.

Our implementation of the HEIDI test follows the method as described by Zhu et al.[6]. In short, for specific region, we compute statistics $T_H = \sum_i^m z_{d(i)}^2$, where $m$ is the number of markers selected for the test and $z_{d(i)}$ is the scaled measure of deviation of the ratio of the regression coefficients from the primary and the secondary GWAS from that at the top associated SNP (see original work for details). In analysis, we have used the same matrices of LD as these used of the PAINTOR analysis (see above). For testing, we have selected up to 20 SNPs in the following manner:

1. Define the set of eligible SNPs as these that fall into +/- 250 kb from the top primary GWAS association, having $\chi^2 > 10$ in primary GWAS, and having results reported in secondary GWAS
2. Make empty "target" ($S_t$) and "rejected" ($S_r$) SNP sets
3. Select current SNP with the lowest $p$ from the primary GWAS that is neither in the target nor in the rejected set
4. If current SNPs has $r^2 > 0.9$ (computed with PLINK 1.9) with any SNP in the target SNP set, add current SNP to the rejected set
5. Otherwise add current SNP to the target set
6. Repeat from the third step until either the eligible SNP set is exhausted or the target set has 20 SNPs
7. Use target set for HEIDI test

In case the number of SNPs was less than 3, no test was performed.

The method was implemented using Python 3.5 as the main programming language, Pandas 1.19.2 (https://conference.scipy.org/proceedings/scipy2010/mckinney.html) for data processing, Dask 0.14.1 (http://dask.pydata.org/) for distributed data pre-processing, Bokeh 1.12.4 (http://bokeh.pydata.org/) for graphics.

(**Table S5**)

#### Fine-mapping and candidate genes investigation

Among the 49 loci, 22 loci were firstly replicated by MVA-significance test and further replicated by correlation or score replications (**Figure S1**). With the evidence of pleiotropy from these two follow-up replications, the top SNPs of these 22 loci were extracted for subsequent investigation (**Table S4**). The list of complex traits showing associations in the same region included Parkinson disease, type 2 diabetes, birth length, age at menarche, years of educatipon attainment, HDL and triglyceride levels, fasting blood glucose and proinsulin (**Table S4**). One of explanation for this overlap may be pleiotropic action of detected variants. Other plausible explanation would be that the region includes different variants affecting different traits, as these variants are in linkage disequilibrium. To distinguish between these two hypotheses, we have performed the HEIDI test[6]. For the regions of interest we have performed meta-analysis of UKB and GIANT data for six anthropometric traits. The regional association pattern for the trait with strongest association was compared to the regional association pattern for traits for which summary association statistics were freely available, including type 2 diabetes, birth length, age at menarche, years of educational attainment, HDL and triglyceride levels, fasting blood glucose.

For the top seven highly pleiotropic loci (number of pleiotropic records > 20 from PhenoScanner) in our new discoveries (**Fig. 1**, **Table S2**), we also extracted all the annotated genes within a ±500kb window of each top SNP according to the hg19 reference panel. Subsequently, we checked whether mouse knock-out (KO) data are available for any of these genes on growth/size/body region phenotypes, in the database of the International Mouse Phenotyping Consortium (IMPC; http://www.mousephenotype.org/). The required phenotypic data have been established for ten genes, i.e. *ELK4*, *SLC45A3* (top SNP rs823114) on chromosome 1, *KLF14* (top SNP rs972283) on chromosome 7, and *JMJD1C* (top SNP rs10761785) on chromosome 10, *STARD10*, *ARAP1*, *ATG16L2* (top SNP rs1552224) on chromosome 11, and *YDJC*, *CCDC116*, *YPEL1* (top SNP rs181362) on chromosome 22.

At a significance threshold of $1 \times 10^{-4}$, three of these ten genes, *ELK4*, *ARAP1*, and *YDJC* have significantly different phenotypic values in their corresponding KO mice (**Figure S8**). For instance, *ARAP1* mutant mice have significantly larger body weight and reduced number of caudal vertebrae, and *YDJC* mutants have significantly reduced lean body mass (**Figure S9**). In the whole IMPC database, to date, 647 genes across the whole genome have significant associations with growth/size/body region phenotypes at the same 1e-4 threshold. This is less than 2.5% of all the 26292 annotated genes in the hg19 panel. Except for *ATG16L2*, *YPEL1*, and *SLC45A3*, the other seven genes have significant KO effects on the phenotypes with a liberal 0.05 threshold.

### Supplemental Figures

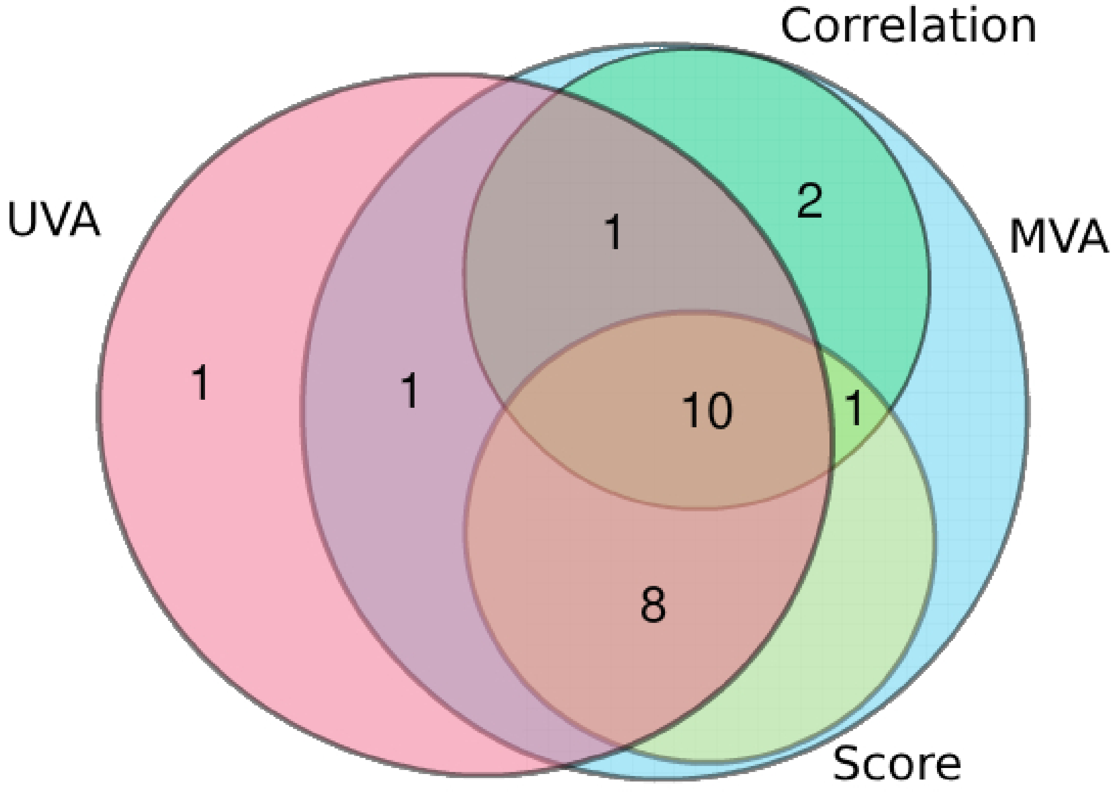


Figure S1: The number of loci replicated by different replication strategies. Each circle in the Venn diagram represents one replication strategy. The four replication strategies are: single-trait GWAS replication (UVA), MVA-significance replication (MVA), MVA-score replication (Score) and correlation replication (Correlation).

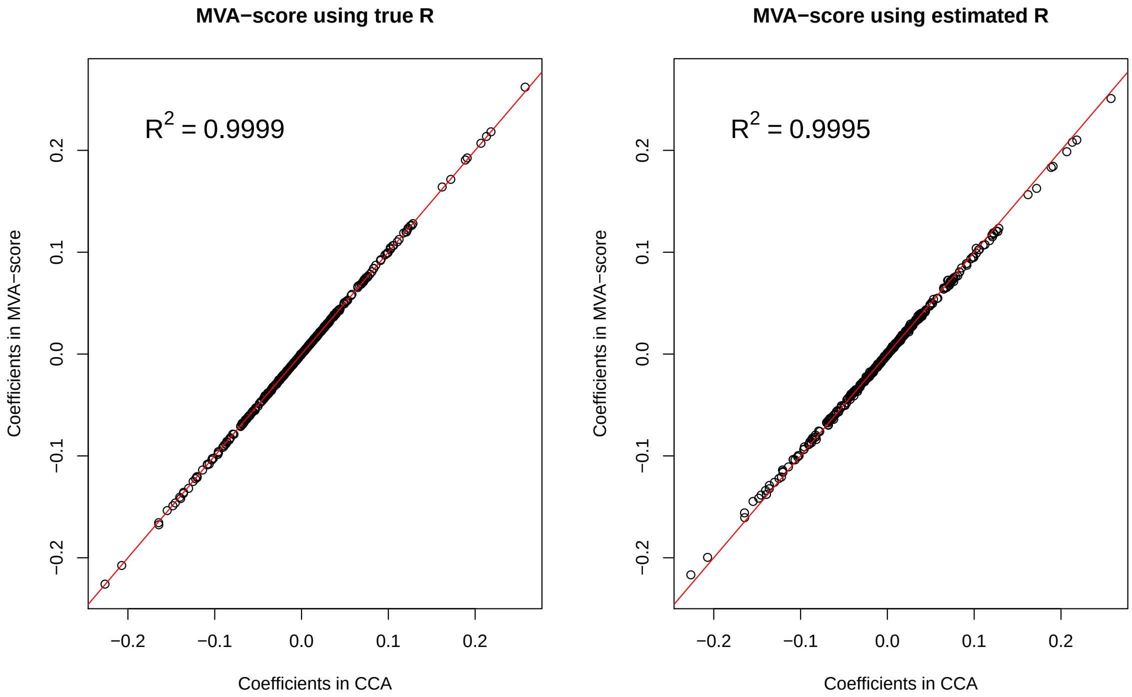


Figure S2: Plot of coefficients from MVA-score against those from CCA. We randomly sampled 50,000 individuals and 100 SNPs from UKB chromosome 22. Then for each SNP, we generated six marginal effects and traits. The estimated shrinkage phenotypic correlation matrix from GIANT was used as the phenotypic correlation matrix of simulated traits. Each SNP explains 0.01% variance of each trait. The x-axis represents the coefficients in reverse regression estimated from CCA based on individual-level data. The y-axis represents the coefficients estimated by MVA-score using summary statistics. In the left panel, true phenotypic correlation matrix was used in MVA-score. In the right panel, the phenotypic correlation matrix was estimated using the t-statistics from 10,000 simulated SNPs without effect. In both cases, MVA-score are almost equivalent to CCA.

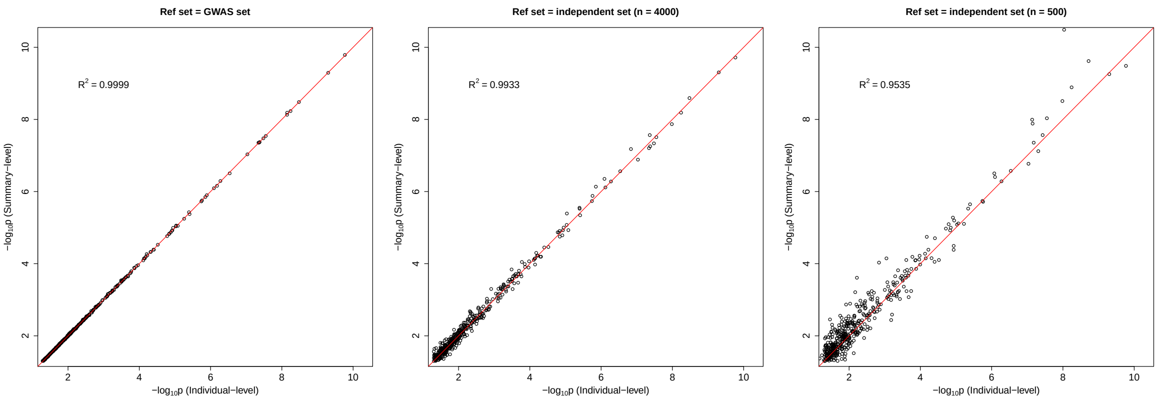


Figure S3: Plot of -$\log_{10}$ p-values from cMVA against those from multiple regression based on individual-level data. We randomly sampled 50,000 individuals from UKB and took their genotypes of 103 snps around SNP rs132622 as an example. Among the 103 snps, we randomly picked two as causal variants. Then the phenotypes were simulated as: $Y_1 = 0.3X_1 + 0.1X_2 + \epsilon_1$, $Y_2 = 0.3X_1 - 0.1X_2 + \epsilon_2$, where $\mathrm{Var}(\epsilon_1) = \mathrm{Var}(\epsilon_2) = 2\mathrm{Cov}(\epsilon_1, \epsilon_2) = 100$. 40,000 individuals among 50,000 were used to perform GWAS and to generate summary statistics. A subset of the rest 10,000 individuals is used as reference sample to approximate LD matrix. The phenotypic correlation matrix in cMVA was estimated using the t-statistics from 10,000 simulated SNPs without effect. The reference sample used in cMVA is the original GWAS sample (left panel), an independent sample with 4,000 individuals (middle panel) or an independent sample with 500 individuals (right panel).

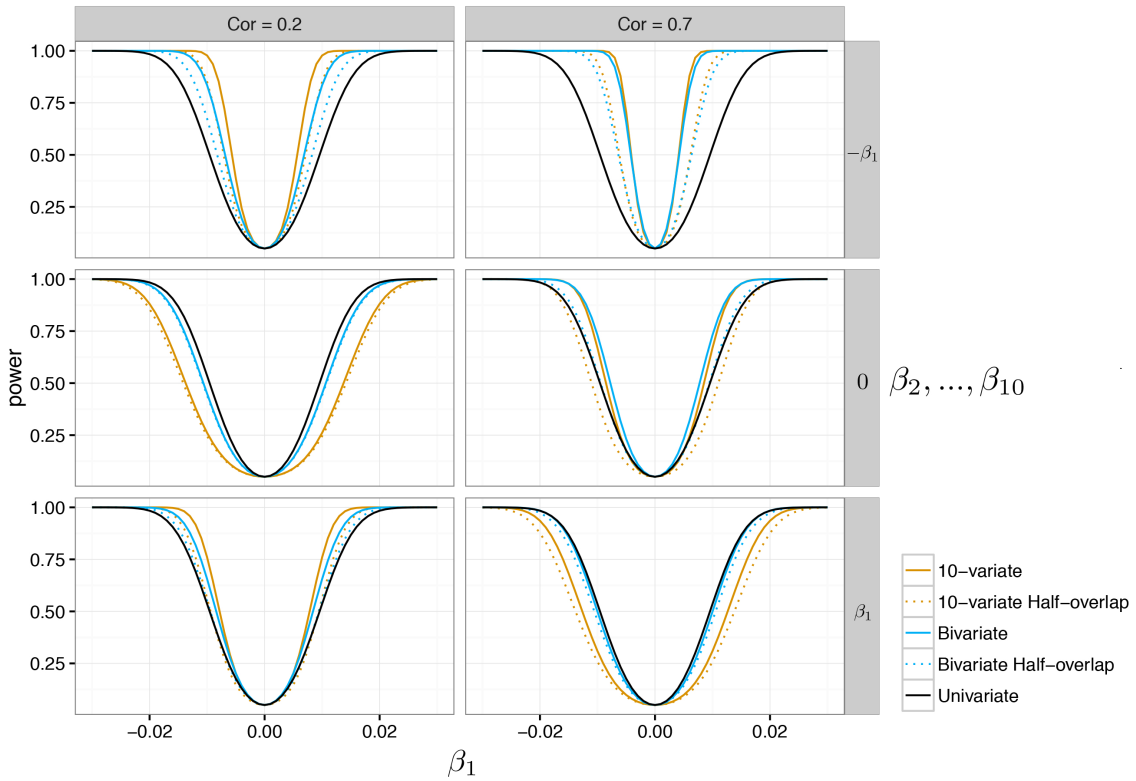


Figure S4: Power comparison of multi-trait and single-trait analyses. Cor: correlation coefficient between each pair of phenotypes; $\beta_i$: the genetic effect on the $i$-th phenotype; Half-overlap: each pair of the phenotypes share only 50% of the genotyped samples.

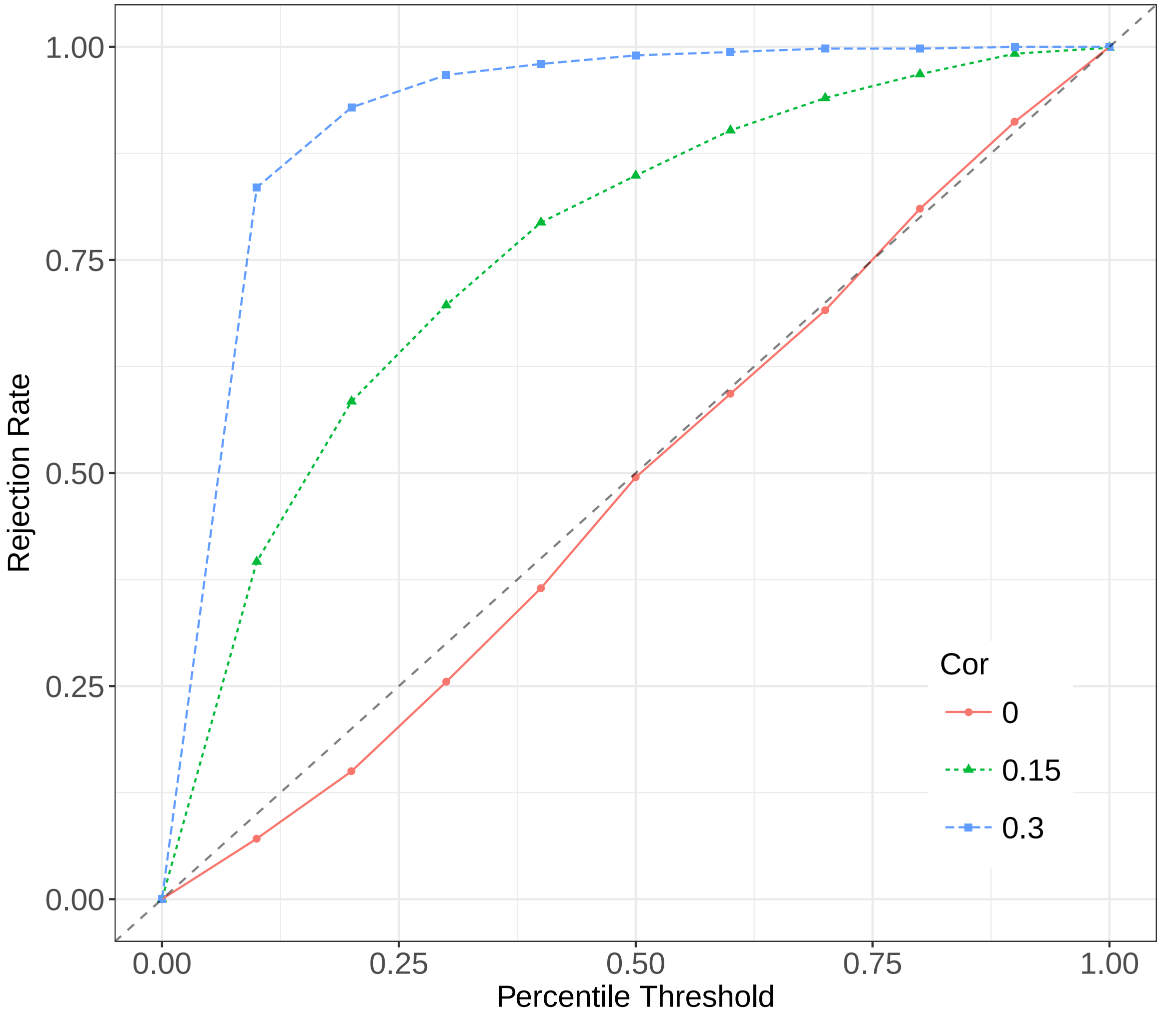


Figure S5: The rejection rate of correlation test under different scenarios. In correlation test, the null hypothesis is rejected if the lower bound of MC-based CI is larger than 0. The lines represent the change of rejection rate when the percentile used for the lower bound is set to be different values. For example, percentile threshold $= 0.05$ means the test is based on whether 0 is above or below the 5th percentile of MC-based distribution for $\tau_\beta$. The first step is to simulate 12 traits in two samples. We firstly simulated 1,000 groups of marginal effects. In each group, 12 pairs of coefficients were drawn from $\mathcal{N}_2(\mathbf{0}, \mathbf{I})$, which are the marginal effects of a SNP on 12 traits in discovery and replication sample. Those groups with $\tau_\beta = 0$ or 0.15 or 0.3 were saved for next step. We then simulated a SNP for 10,000 individuals. The SNP explains 0.1% variance of each trait. After this, we sampled one group of coefficients from the saved groups and simulated phenotypes. The phenotypic correlation matrix of the 12 simulated traits is set as a block diagonal matrix, where the first $6 \times 6$ is the estimated shrinkage phenotypic correlation matrix from GIANT and the second $6 \times 6$ is the phenotypic correlation matrix from UKB. Then we performed the replication test and got the parametric bootstrap distribution of $\tau_\beta$. Each dot is based on 1,000 simulations. This figure shows that the test has no inflation or deflation when the true $\tau_\beta = 0$. The power of this test increases as the true $\tau_\beta$ becomes larger.

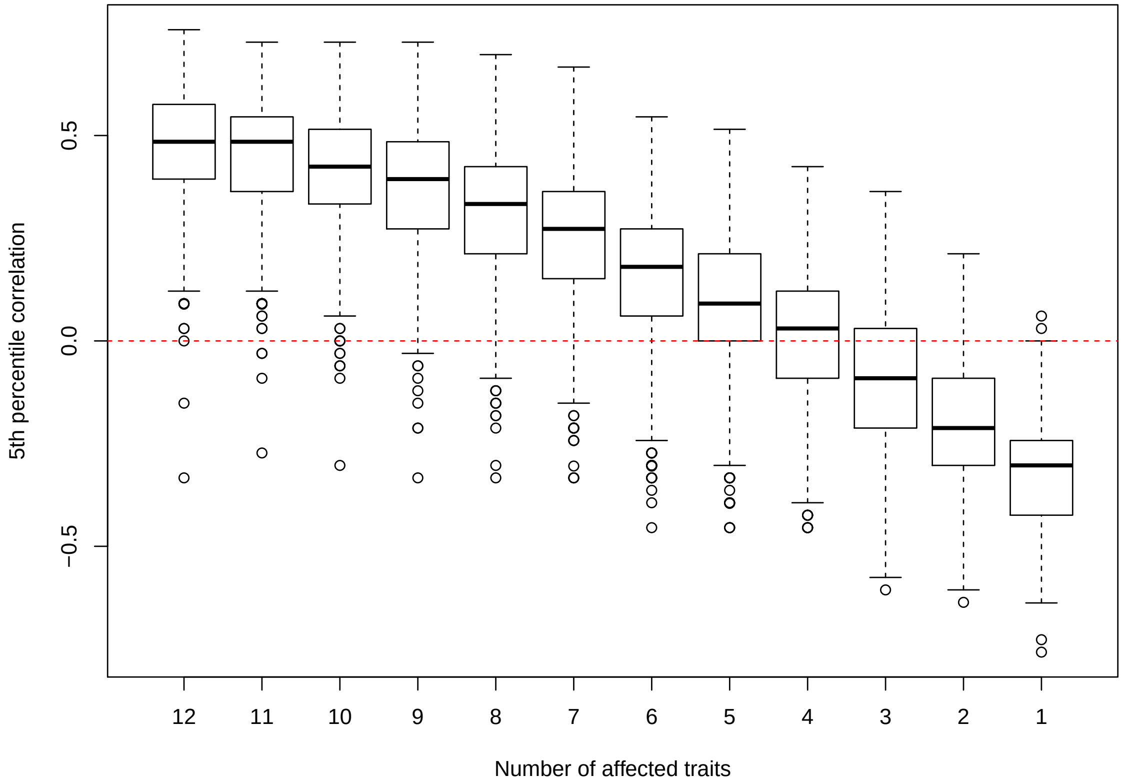


Figure S6: The performance of correlation replication when zero effect sizes exist. In this simulation, we set the marginal effects of a SNP on 12 traits in discovery and replication sample to be same. We firstly simulated 1,000 groups of marginal effects. In each group, 12 coefficients were drawn from $\mathcal{N}(0,1)$, which are the marginal effects of a SNP on 12 traits in discovery and replication sample. Because the effect sizes for each trait are same across two samples, the true $\tau_\beta = 1$. To simulate the impact of zero effect sizes on the MC-based distribution of $\tau_\beta$ in correlation test, we set the first several effect sizes as zero. In this case, the true $\tau_\beta = 1$ still, but the MC-based distribution of $\tau_\beta$ would change. We then simulated a SNP for 10,000 individuals. The SNP explains 0.1% variance of each trait. The phenotypic correlation matrix of the 12 simulated traits is set as a block diagonal matrix, where the first $6 \times 6$ is the estimated shrinkage phenotypic correlation matrix from GIANT and the second $6 \times 6$ is the phenotypic correlation matrix from UKB. After this, we sampled one group of coefficients from the 1,000 groups and simulated phenotypes. Then we performed the replication test and got the parametric bootstrap distribution of $\tau_\beta$. The x-axis represents the number of traits on which the SNP has non-zero effect. The y-axis is the 5th percentile of the MC-based distribution of $\tau_\beta$.

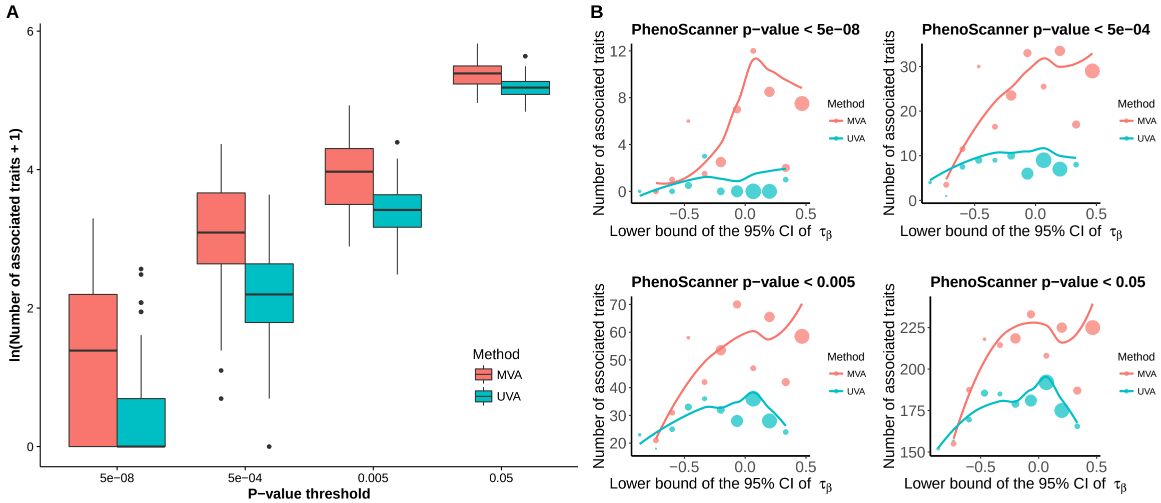


Figure S7: The pleiotropic effects of six-traits MVA-only and UVA-only loci in GIANT2015 across different PhenoScanner p-value threshold. (A) The x-axis represents p-value threshold in PhenoScanner; the y-axis is the natural logarithm of the number of associated traits plus one. (B) The parametric bootstrap distribution is computed for each locus based on the summary statistics of the six anthropometric traits in GIANT2015 and UKB. In each panel, the x-axis represents the lower bound of 95% CI of Kendall's tau. The y-axis is the number of associated traits. To facilitate visualization, loci with the same lower bound values are clustered. At each lower bound value, only the median of the number of associated traits is plotted for each method. The proportion of each cluster is represented by diameter of dots, which is computed separately for MVA and UVA. The curves are based on LOESS fit using data without clustering.

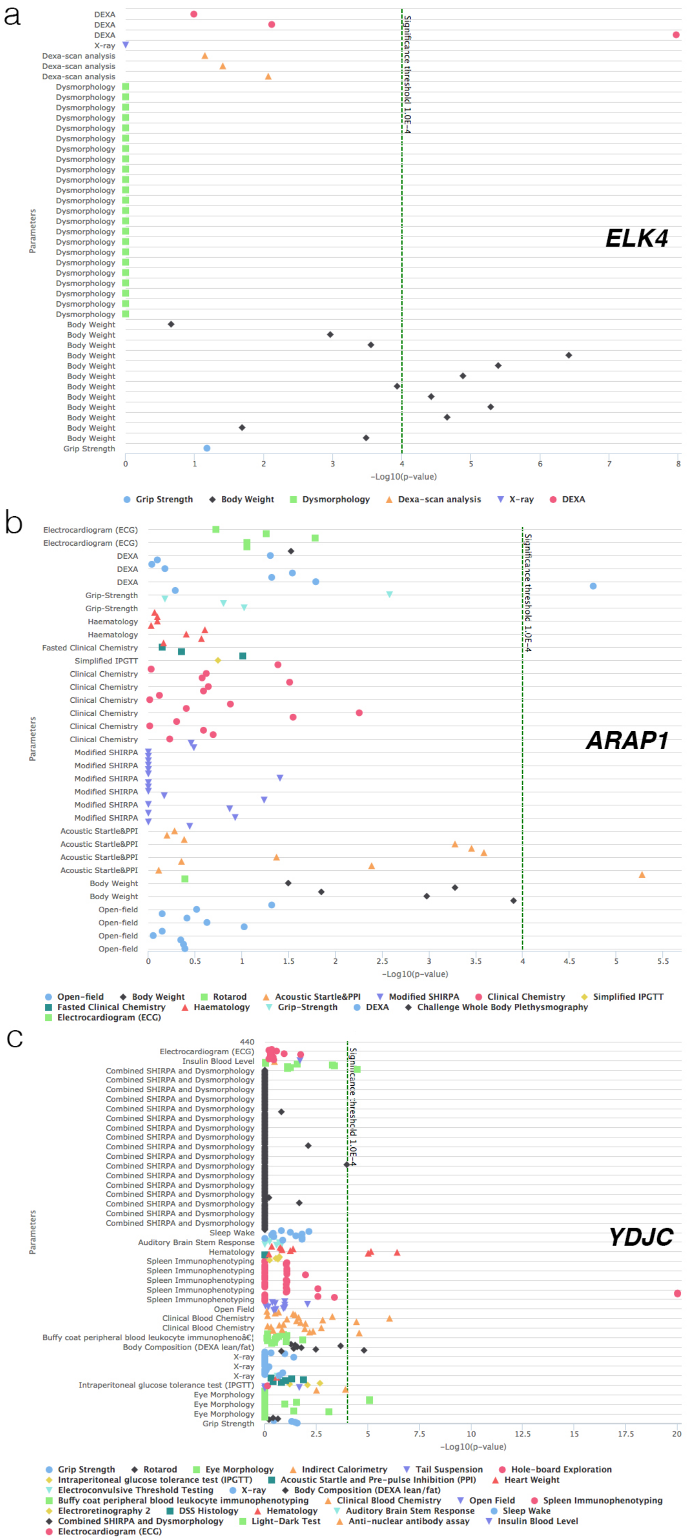


Figure S8: Effects of ELK4, ARAP1, and YDJC knock-out in mice across different phenotypes. The figure data were extracted from the International Mouse Phenotyping Consortium.

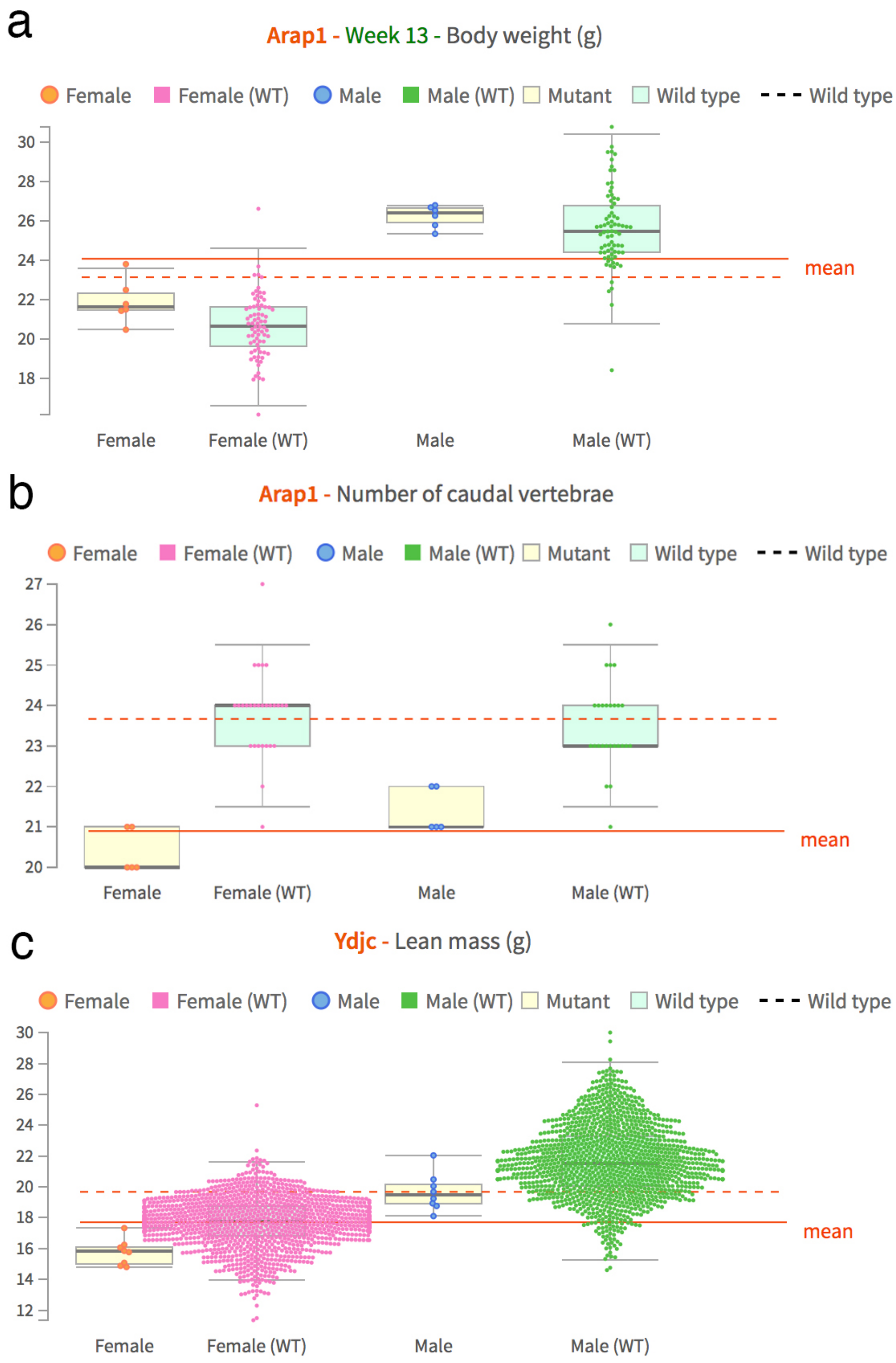


Figure S9: ARAP1 and YDJC knock-out mice show difference in body mass and caudal vertebrae. The figure data were extracted from the International Mouse Phenotyping Consortium.

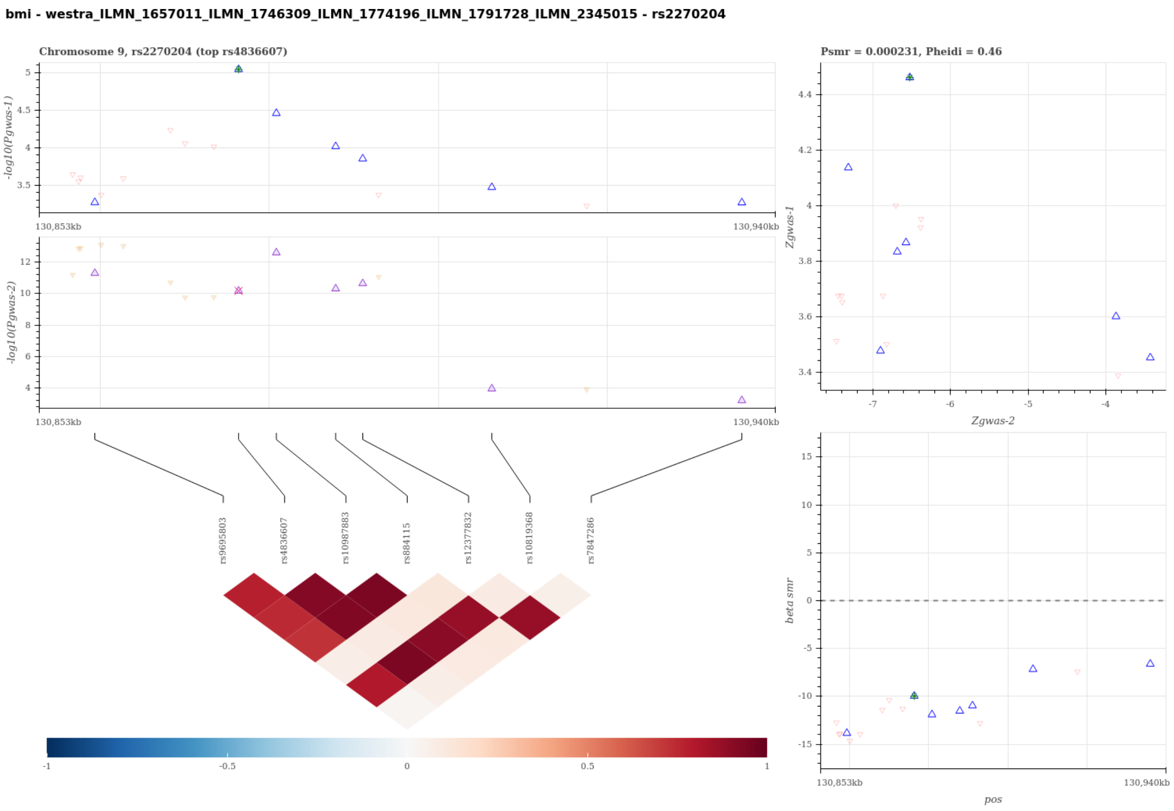


Figure S10: Visualization of the SMR-HEIDI test results to predict candidate gene for BMI using probe ILMN 1657011 ILMN 1746309 ILMN 1774196 ILMN 1791728 ILMN 2345015. A statistical significant SMR test and non-significant HEIDI test suggest that the gene expression of the probe-tagged gene has a cis-eQTL signal (gwas-2) sharing the same single causal variant with BMI (gwas-1). The top left panel compares the regional association results in -log scale p-values between BMI and the eQTL scan. The top right panel compares the regional Z-statistics of the two association scans for the SMR test, and the bottom right panel visualizes the SMR causal effect estimates, i.e. the ratio of the genetic effect on BMI to that on the gene expression tagged by probe ILMN 1657011 ILMN 1746309 ILMN 1774196 ILMN 1791728 ILMN 2345015, for the heterogeneity HEIDI test. The Bottom left panel visualizes the linkage disequilibrium structure of the region. In the plots, each triangle represents a SNP, where the pruned SNP for the SMR-HEIDI test are shown in bigger blue triangles and the top gwas-1 SNP is marked with extra colors.

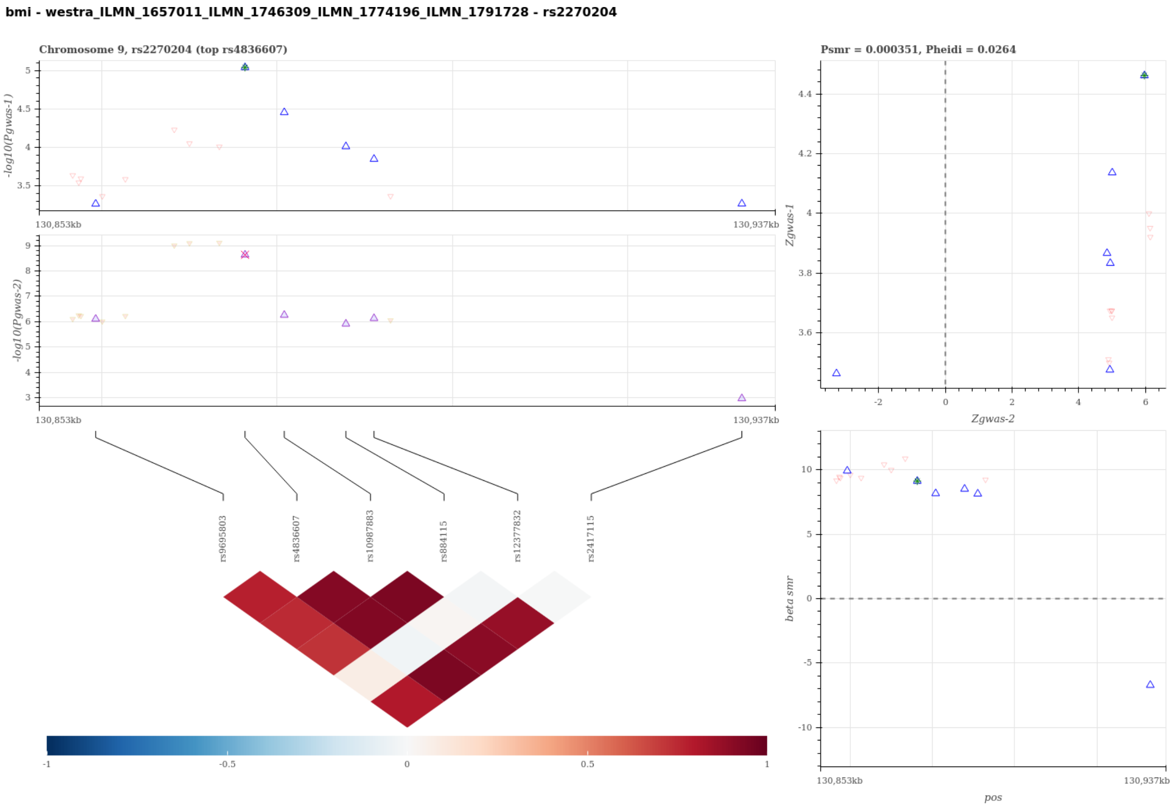


Figure S11: Visualization of the SMR-HEIDI test results to predict candidate gene for BMI using probe ILMN 1657011 ILMN 1746309 ILMN 1774196 ILMN 1791728. A statistical significant SMR test and non-significant HEIDI test suggest that the gene expression of the probe-tagged gene has a cis-eQTL signal (gwas-2) sharing the same single causal variant with BMI (gwas-1). The top left panel compares the regional association results in -log scale p-values between BMI and the eQTL scan. The top right panel compares the regional Z-statistics of the two association scans for the SMR test, and the bottom right panel visualizes the SMR causal effect estimates, i.e. the ratio of the genetic effect on BMI to that on the gene expression tagged by probe ILMN 1657011 ILMN 1746309 ILMN 1774196 ILMN 1791728, for the heterogeneity HEIDI test. The Bottom left panel visualizes the linkage disequilibrium structure of the region. In the plots, each triangle represents a SNP, where the pruned SNP for the SMR-HEIDI test are shown in bigger blue triangles and the top gwas-1 SNP is marked with extra colors.

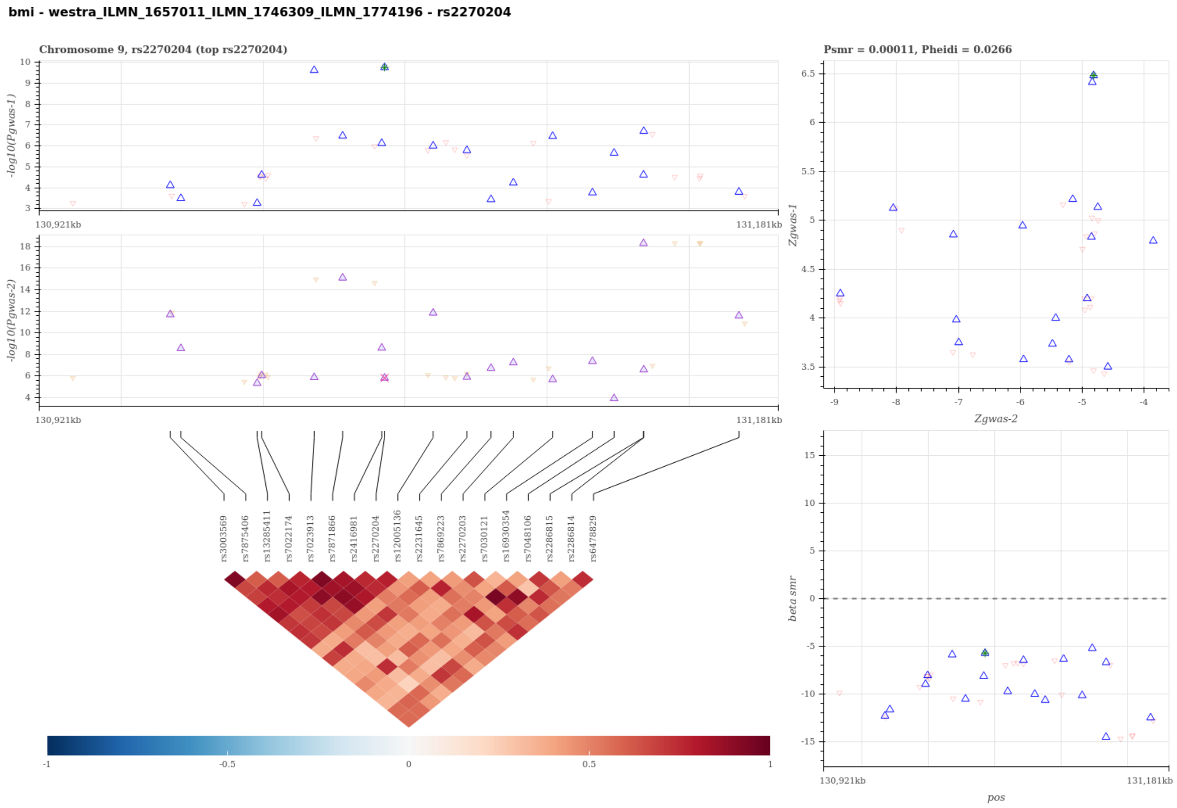


Figure S12: Visualization of the SMR-HEIDI test results to predict candidate gene for BMI using probe ILMN 1657011 ILMN 1746309 ILMN 1774196. A statistical significant SMR test and non-significant HEIDI test suggest that the gene expression of the probe-tagged gene has a cis-eQTL signal (gwas-2) sharing the same single causal variant with BMI (gwas-1). The top left panel compares the regional association results in -log scale p-values between BMI and the eQTL scan. The top right panel compares the regional Z-statistics of the two association scans for the SMR test, and the bottom right panel visualizes the SMR causal effect estimates, i.e. the ratio of the genetic effect on BMI to that on the gene expression tagged by probe ILMN 1657011 ILMN 1746309 ILMN 1774196, for the heterogeneity HEIDI test. The Bottom left panel visualizes the linkage disequilibrium structure of the region. In the plots, each triangle represents a SNP, where the pruned SNP for the SMR-HEIDI test are shown in bigger blue triangles and the top gwas-1 SNP is marked with extra colors.

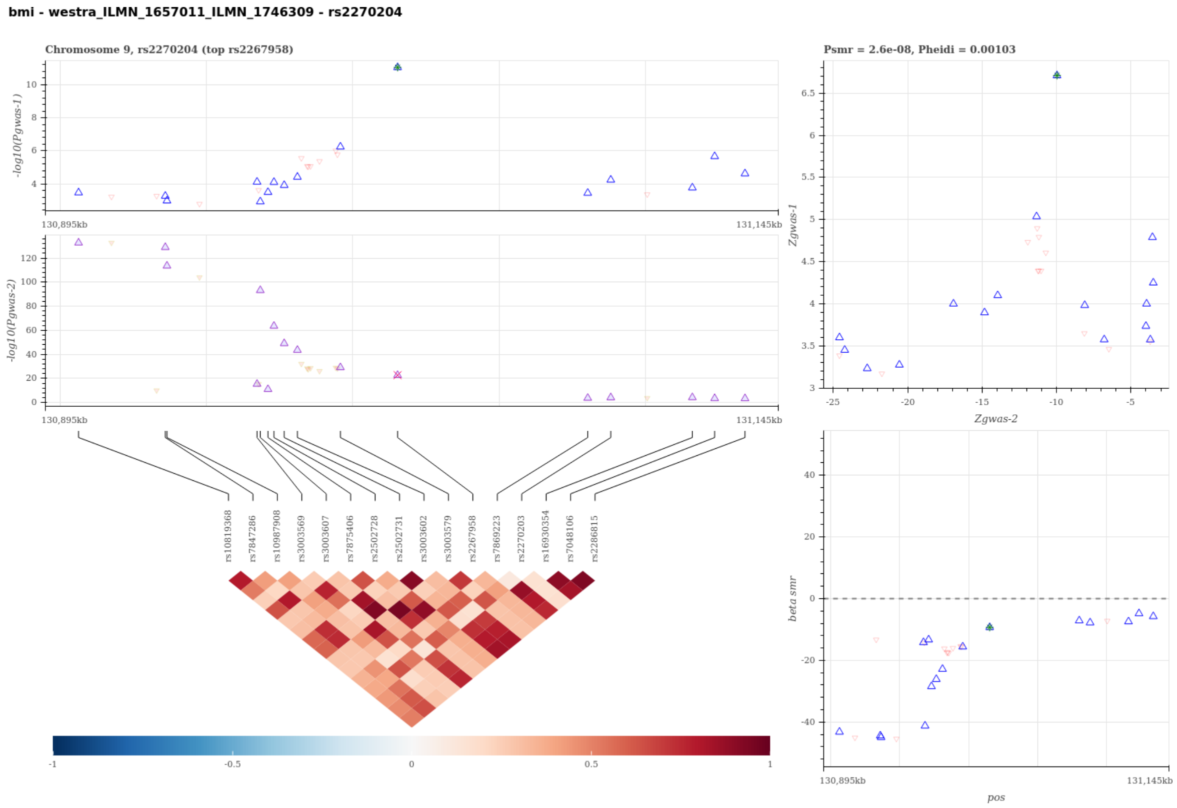


Figure S13: Visualization of the SMR-HEIDI test results to predict candidate gene for BMI using probe ILMN 1657011 ILMN 1746309. A statistical significant SMR test and non-significant HEIDI test suggest that the gene expression of the probe-tagged gene has a cis-eQTL signal (gwas-2) sharing the same single causal variant with BMI (gwas-1). The top left panel compares the regional association results in -log scale p-values between BMI and the eQTL scan. The top right panel compares the regional Z-statistics of the two association scans for the SMR test, and the bottom right panel visualizes the SMR causal effect estimates, i.e. the ratio of the genetic effect on BMI to that on the gene expression tagged by probe ILMN 1657011 ILMN 1746309, for the heterogeneity HEIDI test. The Bottom left panel visualizes the linkage disequilibrium structure of the region. In the plots, each triangle represents a SNP, where the pruned SNP for the SMR-HEIDI test are shown in bigger blue triangles and the top gwas-1 SNP is marked with extra colors.

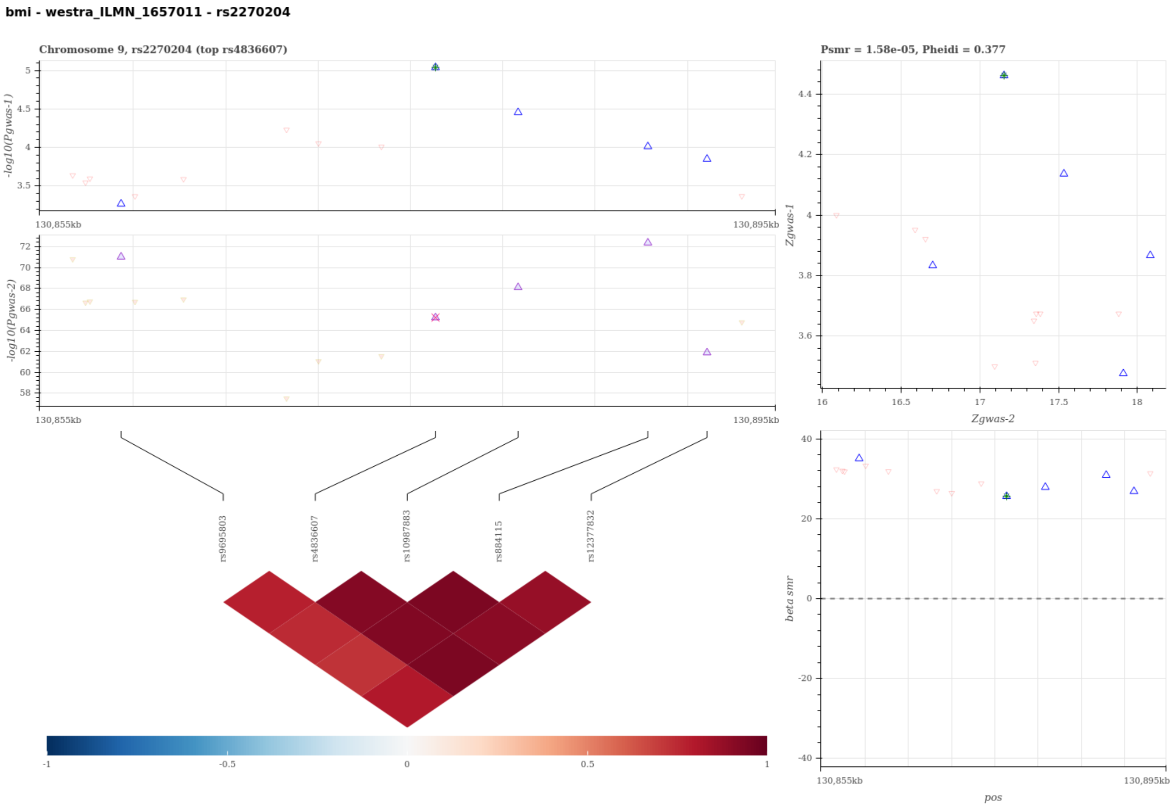


Figure S14: Visualization of the SMR-HEIDI test results to predict candidate gene for BMI using probe ILMN 1657011. A statistical significant SMR test and non-significant HEIDI test suggest that the gene expression of the probe-tagged gene has a cis-eQTL signal (gwas-2) sharing the same single causal variant with BMI (gwas-1). The top left panel compares the regional association results in -log scale p-values between BMI and the eQTL scan. The top right panel compares the regional Z-statistics of the two association scans for the SMR test, and the bottom right panel visualizes the SMR causal effect estimates, i.e. the ratio of the genetic effect on BMI to that on the gene expression tagged by probe ILMN 1657011, for the heterogeneity HEIDI test. The Bottom left panel visualizes the linkage disequilibrium structure of the region. In the plots, each triangle represents a SNP, where the pruned SNP for the SMR-HEIDI test are shown in bigger blue triangles and the top gwas-1 SNP is marked with extra colors.

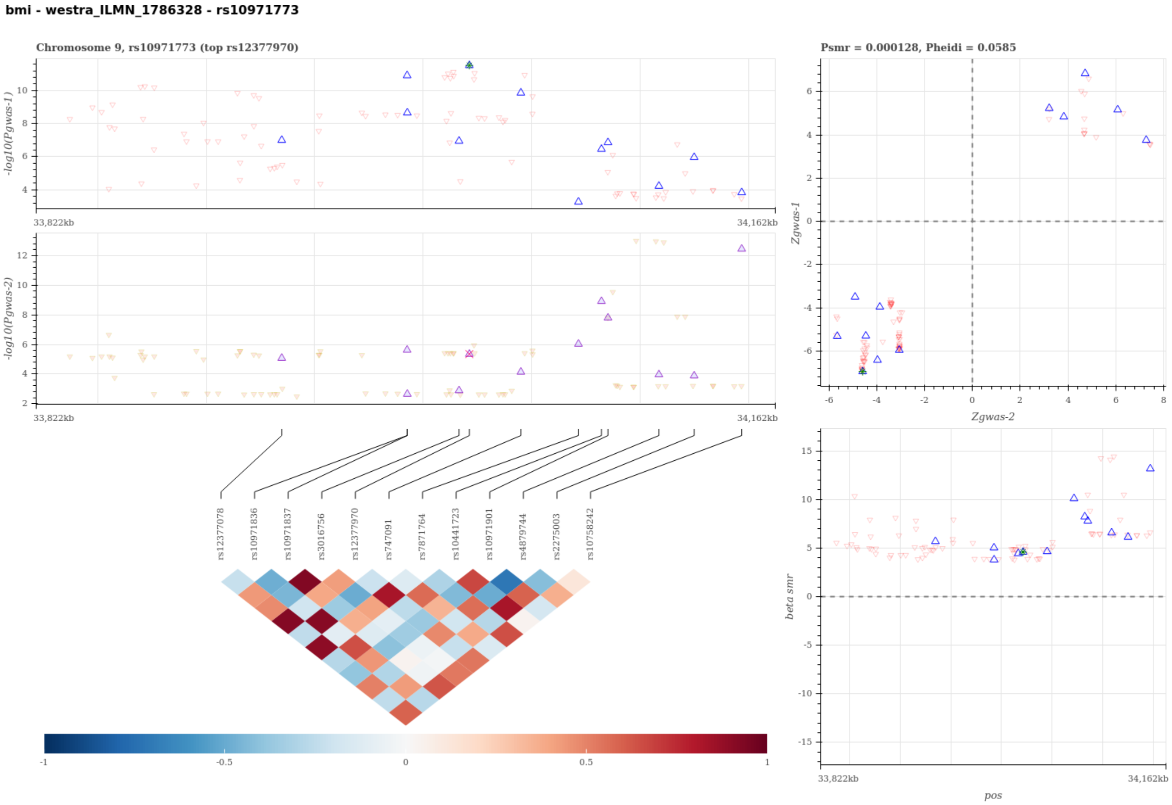


Figure S15: Visualization of the SMR-HEIDI test results to predict candidate gene for BMI using probe ILMN 1786328. A statistical significant SMR test and non-significant HEIDI test suggest that the gene expression of the probe-tagged gene has a cis-eQTL signal (gwas-2) sharing the same single causal variant with BMI (gwas-1). The top left panel compares the regional association results in -log scale p-values between BMI and the eQTL scan. The top right panel compares the regional Z-statistics of the two association scans for the SMR test, and the bottom right panel visualizes the SMR causal effect estimates, i.e. the ratio of the genetic effect on BMI to that on the gene expression tagged by probe ILMN 1786328, for the heterogeneity HEIDI test. The Bottom left panel visualizes the linkage disequilibrium structure of the region. In the plots, each triangle represents a SNP, where the pruned SNP for the SMR-HEIDI test are shown in bigger blue triangles and the top gwas-1 SNP is marked with extra colors.

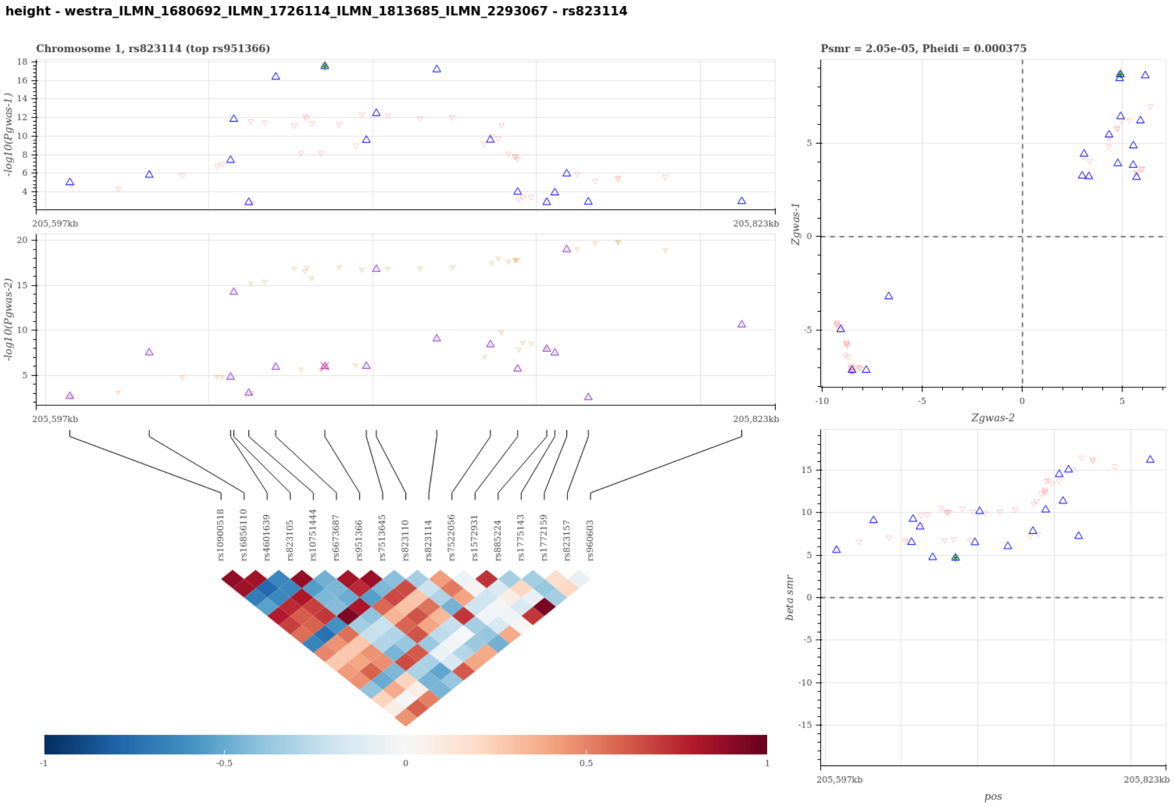


Figure S16: Visualization of the SMR-HEIDI test results to predict candidate gene for HEIGHT using probe ILMN 1680692 ILMN 1726114 ILMN 1813685 ILMN 2293067. A statistical significant SMR test and non-significant HEIDI test suggest that the gene expression of the probe-tagged gene has a cis-eQTL signal (gwas-2) sharing the same single causal variant with HEIGHT (gwas-1). The top left panel compares the regional association results in -log scale p-values between HEIGHT and the eQTL scan. The top right panel compares the regional Z-statistics of the two association scans for the SMR test, and the bottom right panel visualizes the SMR causal effect estimates, i.e. the ratio of the genetic effect on HEIGHT to that on the gene expression tagged by probe ILMN 1680692 ILMN 1726114 ILMN 1813685 ILMN 2293067, for the heterogeneity HEIDI test. The Bottom left panel visualizes the linkage disequilibrium structure of the region. In the plots, each triangle represents a SNP, where the pruned SNP for the SMR-HEIDI test are shown in bigger blue triangles and the top gwas-1 SNP is marked with extra colors.

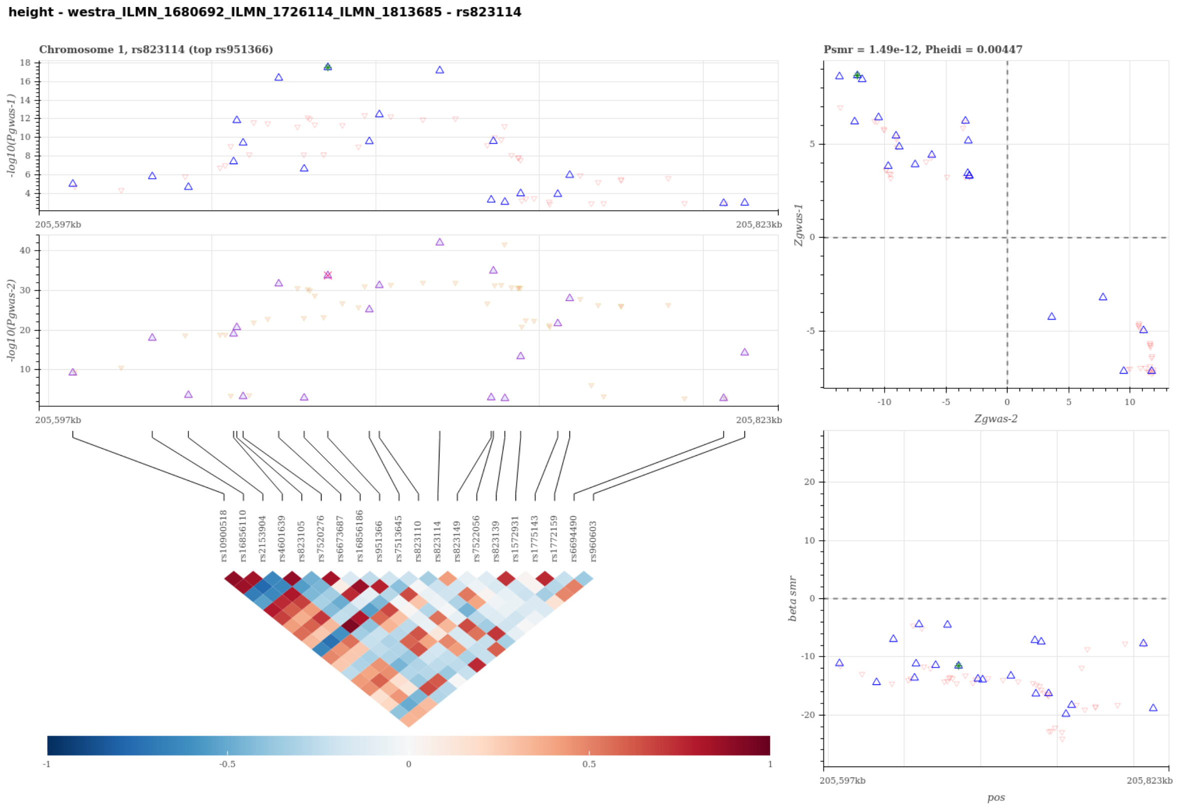


Figure S17: Visualization of the SMR-HEIDI test results to predict candidate gene for HEIGHT using probe ILMN 1680692 ILMN 1726114 ILMN 1813685. A statistical significant SMR test and non-significant HEIDI test suggest that the gene expression of the probe-tagged gene has a cis-eQTL signal (gwas-2) sharing the same single causal variant with HEIGHT (gwas-1). The top left panel compares the regional association results in -log scale p-values between HEIGHT and the eQTL scan. The top right panel compares the regional Z-statistics of the two association scans for the SMR test, and the bottom right panel visualizes the SMR causal effect estimates, i.e. the ratio of the genetic effect on HEIGHT to that on the gene expression tagged by probe ILMN 1680692 ILMN 1726114 ILMN 1813685, for the heterogeneity HEIDI test. The Bottom left panel visualizes the linkage disequilibrium structure of the region. In the plots, each triangle represents a SNP, where the pruned SNP for the SMR-HEIDI test are shown in bigger blue triangles and the top gwas-1 SNP is marked with extra colors.

height - westra_ILMN_1680692_ILMN_1726114 - rs823114

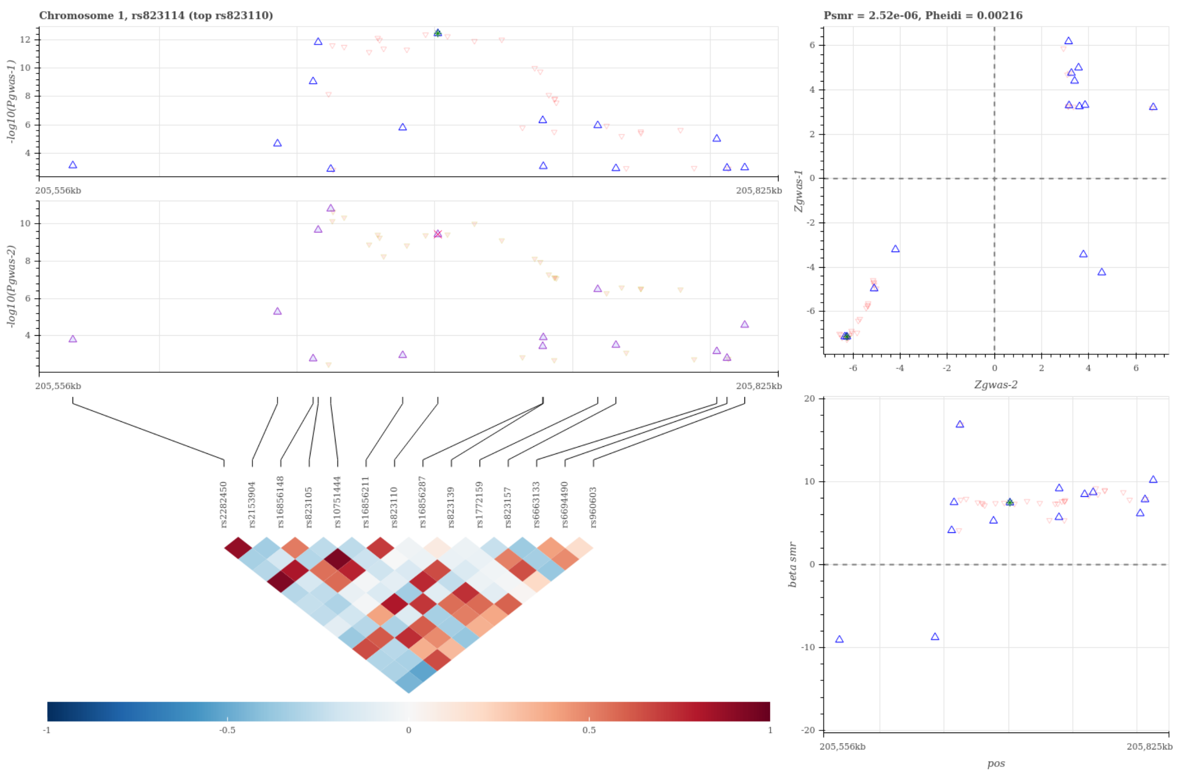


Figure S18: Visualization of the SMR-HEIDI test results to predict candidate gene for HEIGHT using probe ILMN 1680692 ILMN 1726114. A statistical significant SMR test and non-significant HEIDI test suggest that the gene expression of the probe-tagged gene has a cis-eQTL signal (gwas-2) sharing the same single causal variant with HEIGHT (gwas-1). The top left panel compares the regional association results in -log scale p-values between HEIGHT and the eQTL scan. The top right panel compares the regional Z-statistics of the two association scans for the SMR test, and the bottom right panel visualizes the SMR causal effect estimates, i.e. the ratio of the genetic effect on HEIGHT to that on the gene expression tagged by probe ILMN 1680692 ILMN 1726114, for the heterogeneity HEIDI test. The Bottom left panel visualizes the linkage disequilibrium structure of the region. In the plots, each triangle represents a SNP, where the pruned SNP for the SMR-HEIDI test are shown in bigger blue triangles and the top gwas-1 SNP is marked with extra colors.

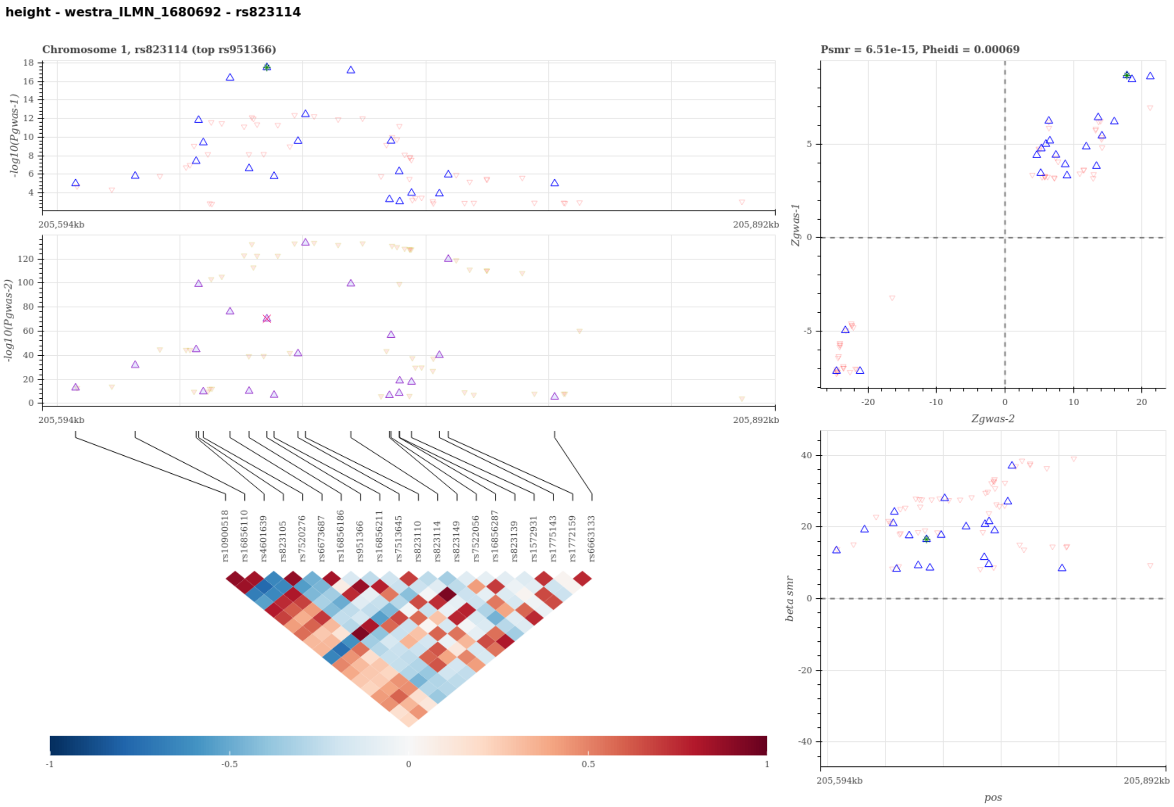


Figure S19: Visualization of the SMR-HEIDI test results to predict candidate gene for HEIGHT using probe ILMN 1680692. A statistical significant SMR test and non-significant HEIDI test suggest that the gene expression of the probe-tagged gene has a cis-eQTL signal (gwas-2) sharing the same single causal variant with HEIGHT (gwas-1). The top left panel compares the regional association results in -log scale p-values between HEIGHT and the eQTL scan. The top right panel compares the regional Z-statistics of the two association scans for the SMR test, and the bottom right panel visualizes the SMR causal effect estimates, i.e. the ratio of the genetic effect on HEIGHT to that on the gene expression tagged by probe ILMN 1680692, for the heterogeneity HEIDI test. The Bottom left panel visualizes the linkage disequilibrium structure of the region. In the plots, each triangle represents a SNP, where the pruned SNP for the SMR-HEIDI test are shown in bigger blue triangles and the top gwas-1 SNP is marked with extra colors.

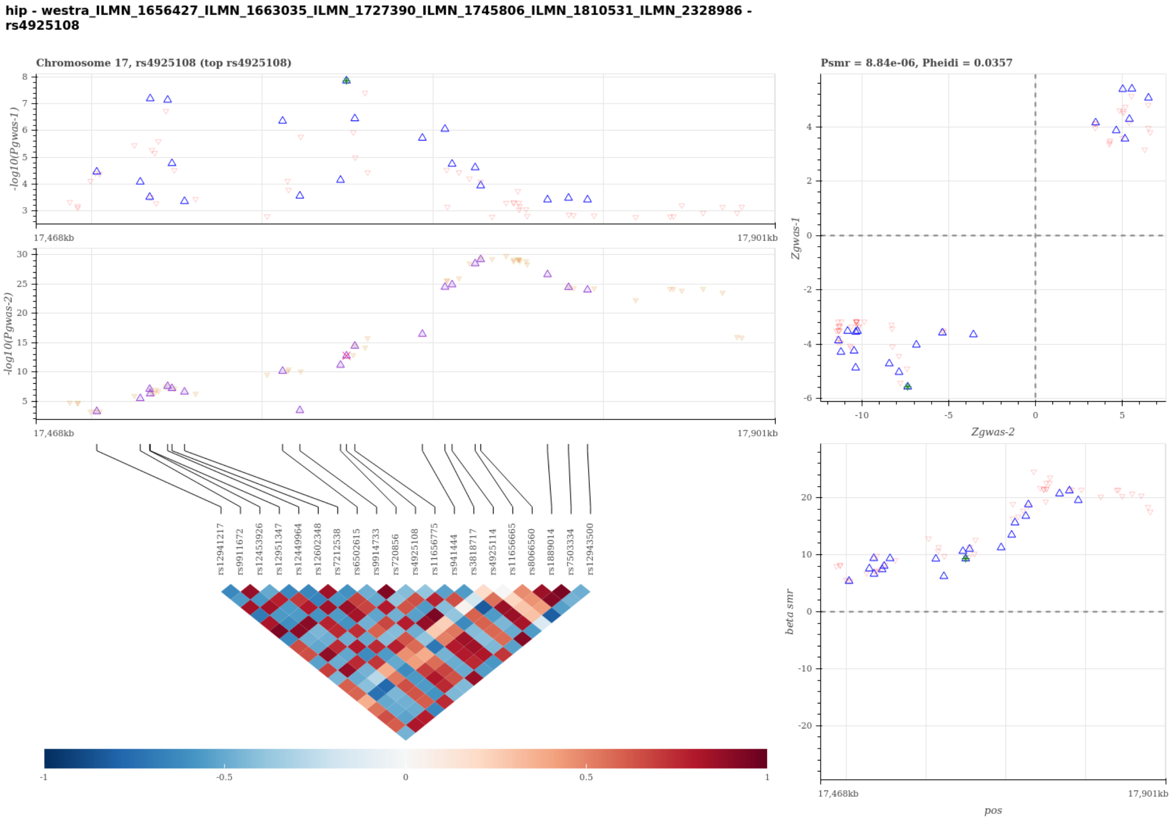


Figure S20: Visualization of the SMR-HEIDI test results to predict candidate gene for HIP using probe ILMN 1656427 ILMN 1663035 ILMN 1727390 ILMN 1745806 ILMN 1810531 ILMN 2328986. A statistical significant SMR test and non-significant HEIDI test suggest that the gene expression of the probe-tagged gene has a cis-eQTL signal (gwas-2) sharing the same single causal variant with HIP (gwas-1). The top left panel compares the regional association results in -log scale p-values between HIP and the eQTL scan. The top right panel compares the regional Z-statistics of the two association scans for the SMR test, and the bottom right panel visualizes the SMR causal effect estimates, i.e. the ratio of the genetic effect on HIP to that on the gene expression tagged by probe ILMN 1656427 ILMN 1663035 ILMN 1727390 ILMN 1745806 ILMN 1810531 ILMN 2328986, for the heterogeneity HEIDI test. The Bottom left panel visualizes the linkage disequilibrium structure of the region. In the plots, each triangle represents a SNP, where the pruned SNP for the SMR-HEIDI test are shown in bigger blue triangles and the top gwas-1 SNP is marked with extra colors.

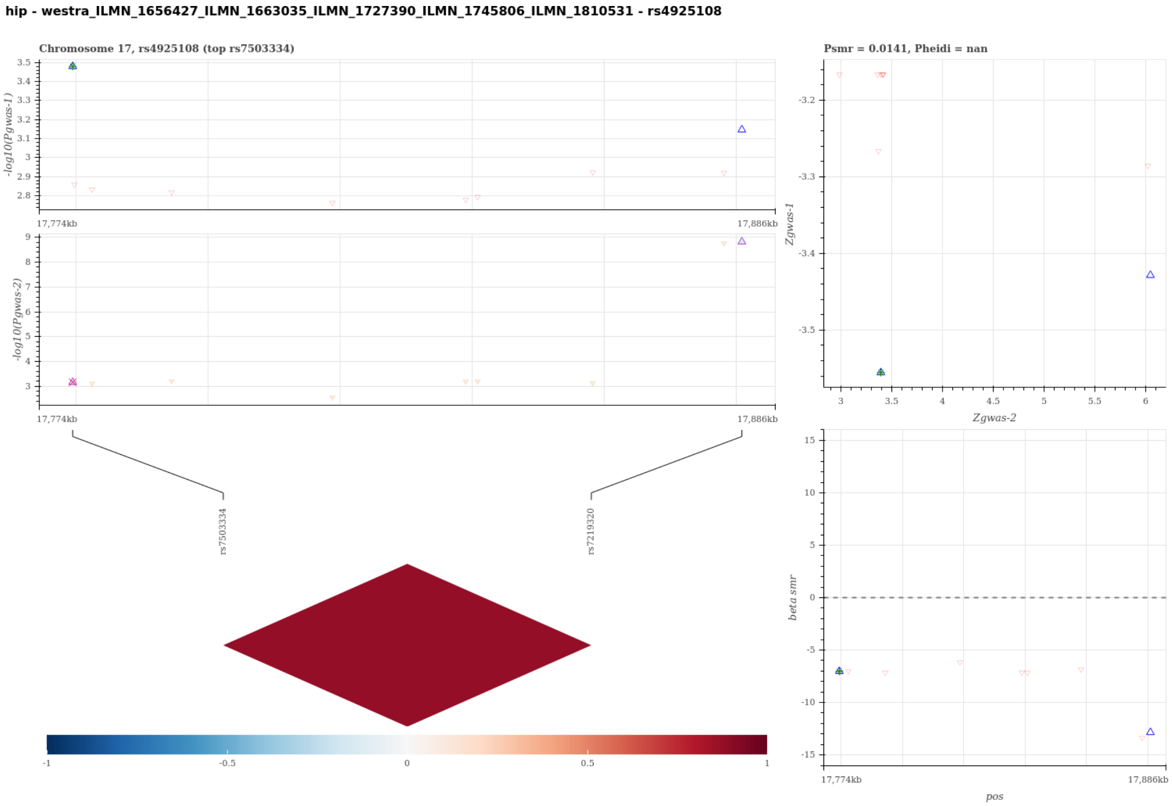


Figure S21: Visualization of the SMR-HEIDI test results to predict candidate gene for HIP using probe ILMN 1656427 ILMN 1663035 ILMN 1727390 ILMN 1745806 ILMN 1810531. A statistical significant SMR test and non-significant HEIDI test suggest that the gene expression of the probe-tagged gene has a cis-eQTL signal (gwas-2) sharing the same single causal variant with HIP (gwas-1). The top left panel compares the regional association results in -log scale p-values between HIP and the eQTL scan. The top right panel compares the regional Z-statistics of the two association scans for the SMR test, and the bottom right panel visualizes the SMR causal effect estimates, i.e. the ratio of the genetic effect on HIP to that on the gene expression tagged by probe ILMN 1656427 ILMN 1663035 ILMN 1727390 ILMN 1745806 ILMN 1810531, for the heterogeneity HEIDI test. The Bottom left panel visualizes the linkage disequilibrium structure of the region. In the plots, each triangle represents a SNP, where the pruned SNP for the SMR-HEIDI test are shown in bigger blue triangles and the top gwas-1 SNP is marked with extra colors.

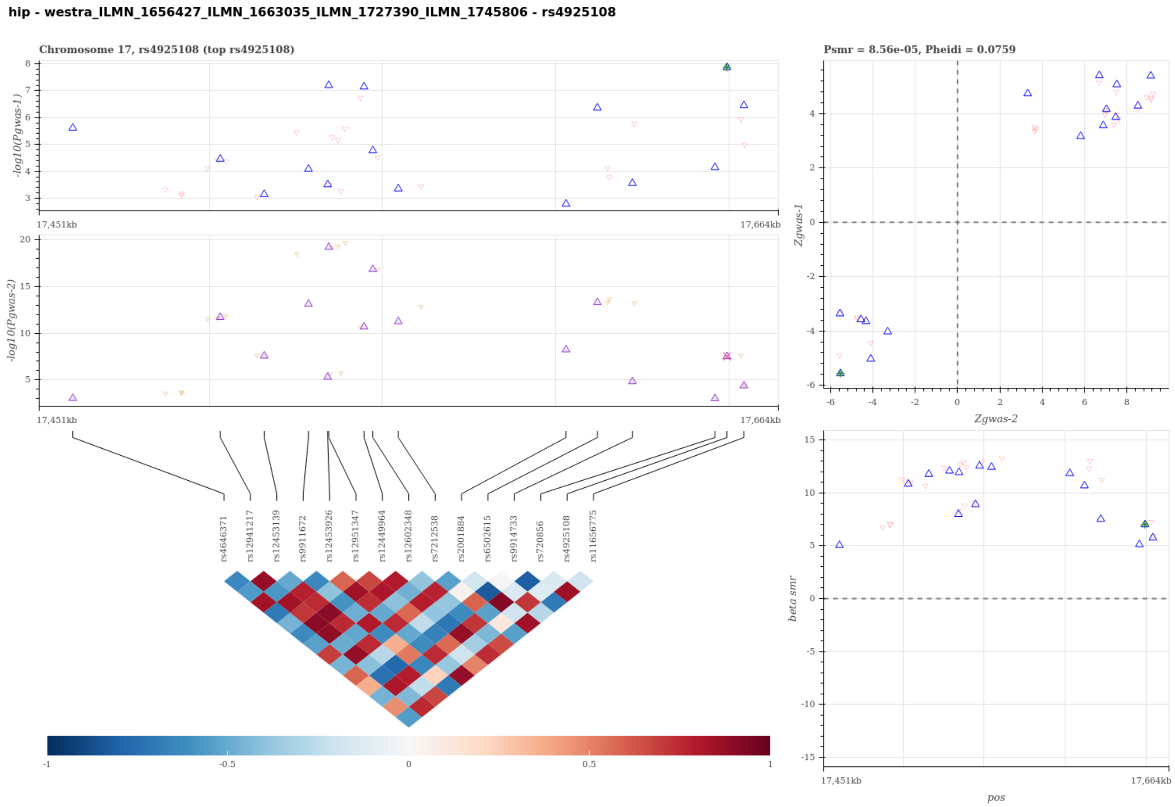


Figure S22: Visualization of the SMR-HEIDI test results to predict candidate gene for HIP using probe ILMN 1656427 ILMN 1663035 ILMN 1727390 ILMN 1745806. A statistical significant SMR test and non-significant HEIDI test suggest that the gene expression of the probe-tagged gene has a cis-eQTL signal (gwas-2) sharing the same single causal variant with HIP (gwas-1). The top left panel compares the regional association results in -log scale p-values between HIP and the eQTL scan. The top right panel compares the regional Z-statistics of the two association scans for the SMR test, and the bottom right panel visualizes the SMR causal effect estimates, i.e. the ratio of the genetic effect on HIP to that on the gene expression tagged by probe ILMN 1656427 ILMN 1663035 ILMN 1727390 ILMN 1745806, for the heterogeneity HEIDI test. The Bottom left panel visualizes the linkage disequilibrium structure of the region. In the plots, each triangle represents a SNP, where the pruned SNP for the SMR-HEIDI test are shown in bigger blue triangles and the top gwas-1 SNP is marked with extra colors.

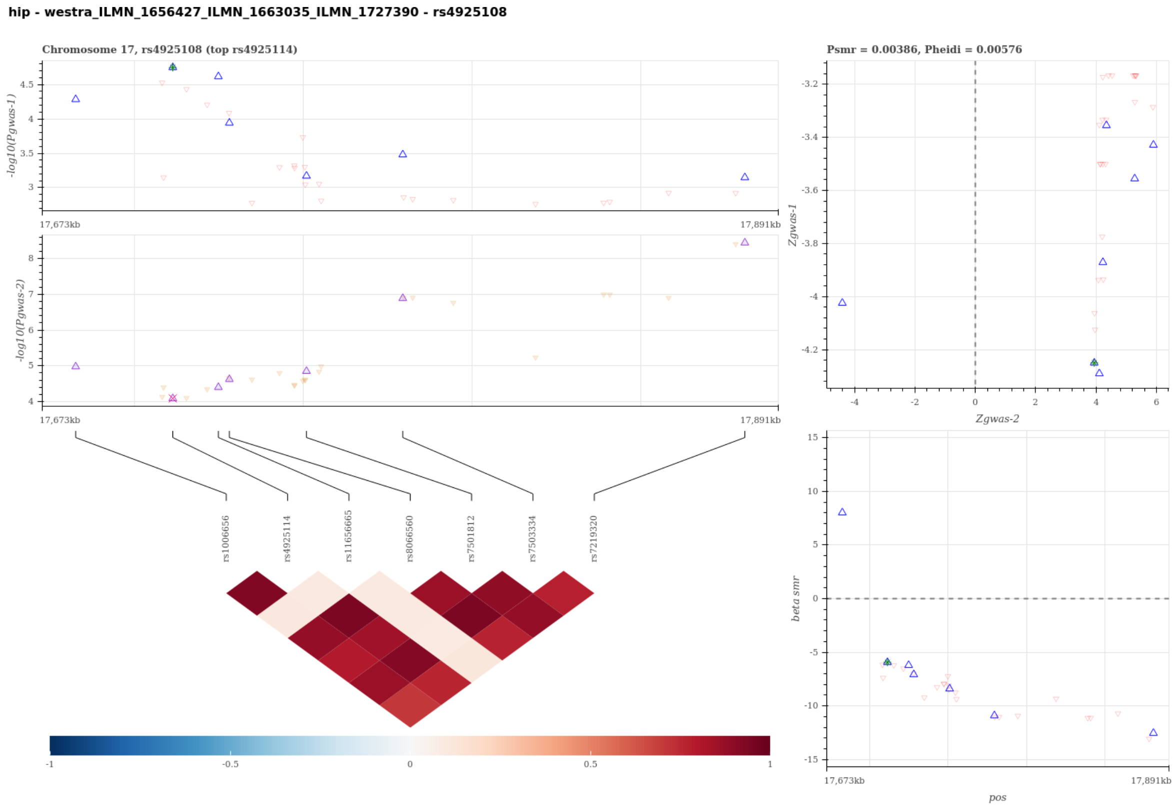


Figure S23: Visualization of the SMR-HEIDI test results to predict candidate gene for HIP using probe ILMN 1656427 ILMN 1663035 ILMN 1727390. A statistical significant SMR test and non-significant HEIDI test suggest that the gene expression of the probe-tagged gene has a cis-eQTL signal (gwas-2) sharing the same single causal variant with HIP (gwas-1). The top left panel compares the regional association results in -log scale p-values between HIP and the eQTL scan. The top right panel compares the regional Z-statistics of the two association scans for the SMR test, and the bottom right panel visualizes the SMR causal effect estimates, i.e. the ratio of the genetic effect on HIP to that on the gene expression tagged by probe ILMN 1656427 ILMN 1663035 ILMN 1727390, for the heterogeneity HEIDI test. The Bottom left panel visualizes the linkage disequilibrium structure of the region. In the plots, each triangle represents a SNP, where the pruned SNP for the SMR-HEIDI test are shown in bigger blue triangles and the top gwas-1 SNP is marked with extra colors.

**hip - westra_ILMN_1656427_ILMN_1663035 - rs4925108**

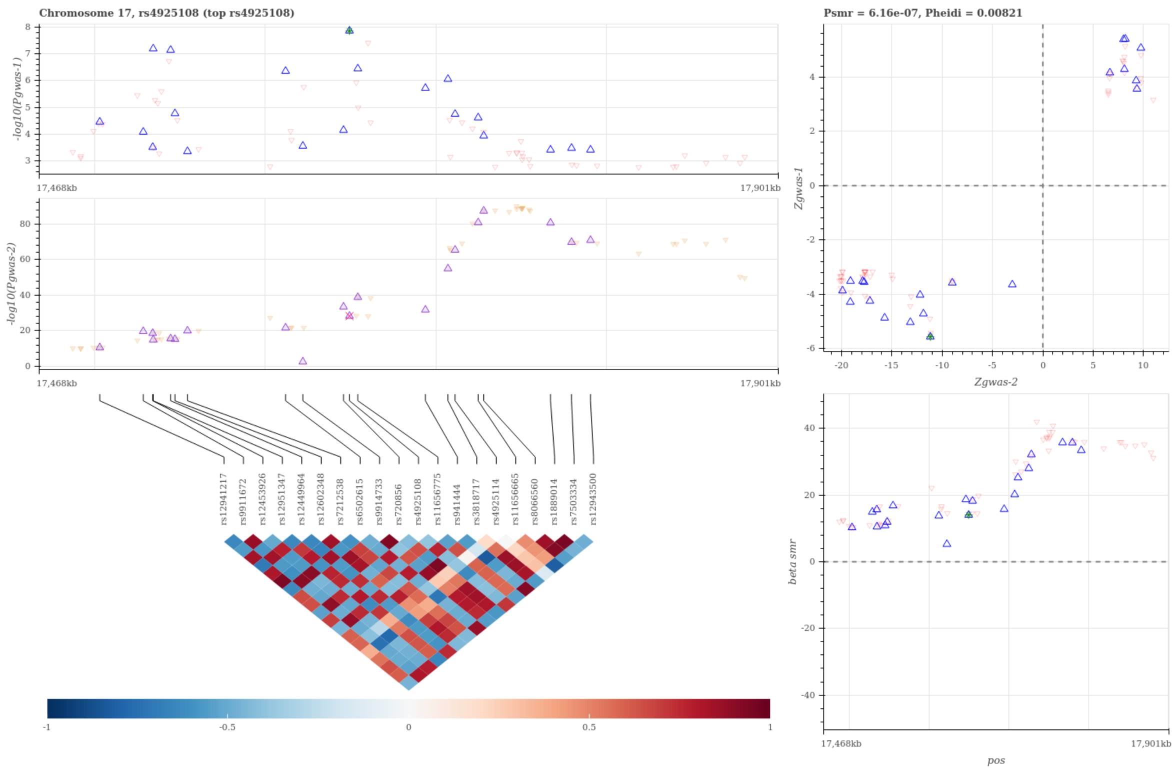


Figure S24: Visualization of the SMR-HEIDI test results to predict candidate gene for HIP using probe ILMN 1656427 ILMN 1663035. A statistical significant SMR test and non-significant HEIDI test suggest that the gene expression of the probe-tagged gene has a cis-eQTL signal (gwas-2) sharing the same single causal variant with HIP (gwas-1). The top left panel compares the regional association results in -log scale p-values between HIP and the eQTL scan. The top right panel compares the regional Z-statistics of the two association scans for the SMR test, and the bottom right panel visualizes the SMR causal effect estimates, i.e. the ratio of the genetic effect on HIP to that on the gene expression tagged by probe ILMN 1656427 ILMN 1663035, for the heterogeneity HEIDI test. The Bottom left panel visualizes the linkage disequilibrium structure of the region. In the plots, each triangle represents a SNP, where the pruned SNP for the SMR-HEIDI test are shown in bigger blue triangles and the top gwas-1 SNP is marked with extra colors.

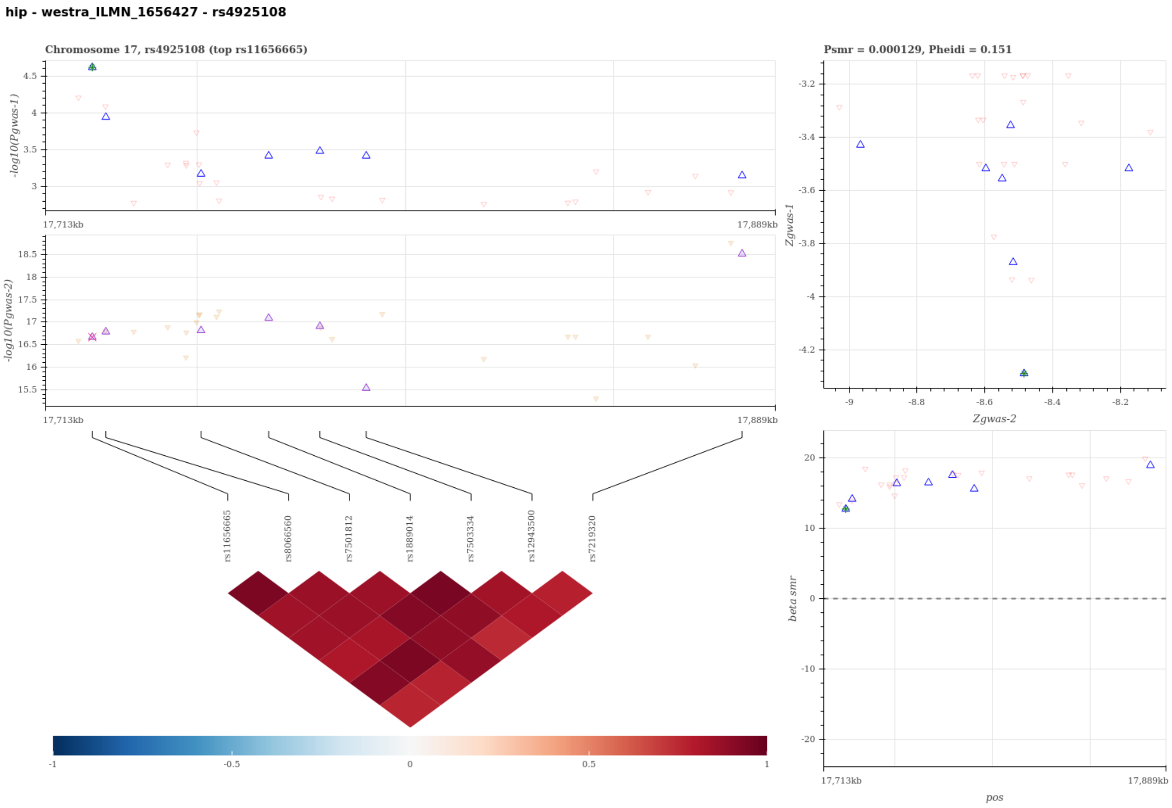


Figure S25: Visualization of the SMR-HEIDI test results to predict candidate gene for HIP using probe ILMN 1656427. A statistical significant SMR test and non-significant HEIDI test suggest that the gene expression of the probe-tagged gene has a cis-eQTL signal (gwas-2) sharing the same single causal variant with HIP (gwas-1). The top left panel compares the regional association results in -log scale p-values between HIP and the eQTL scan. The top right panel compares the regional Z-statistics of the two association scans for the SMR test, and the bottom right panel visualizes the SMR causal effect estimates, i.e. the ratio of the genetic effect on HIP to that on the gene expression tagged by probe ILMN 1656427, for the heterogeneity HEIDI test. The Bottom left panel visualizes the linkage disequilibrium structure of the region. In the plots, each triangle represents a SNP, where the pruned SNP for the SMR-HEIDI test are shown in bigger blue triangles and the top gwas-1 SNP is marked with extra colors.

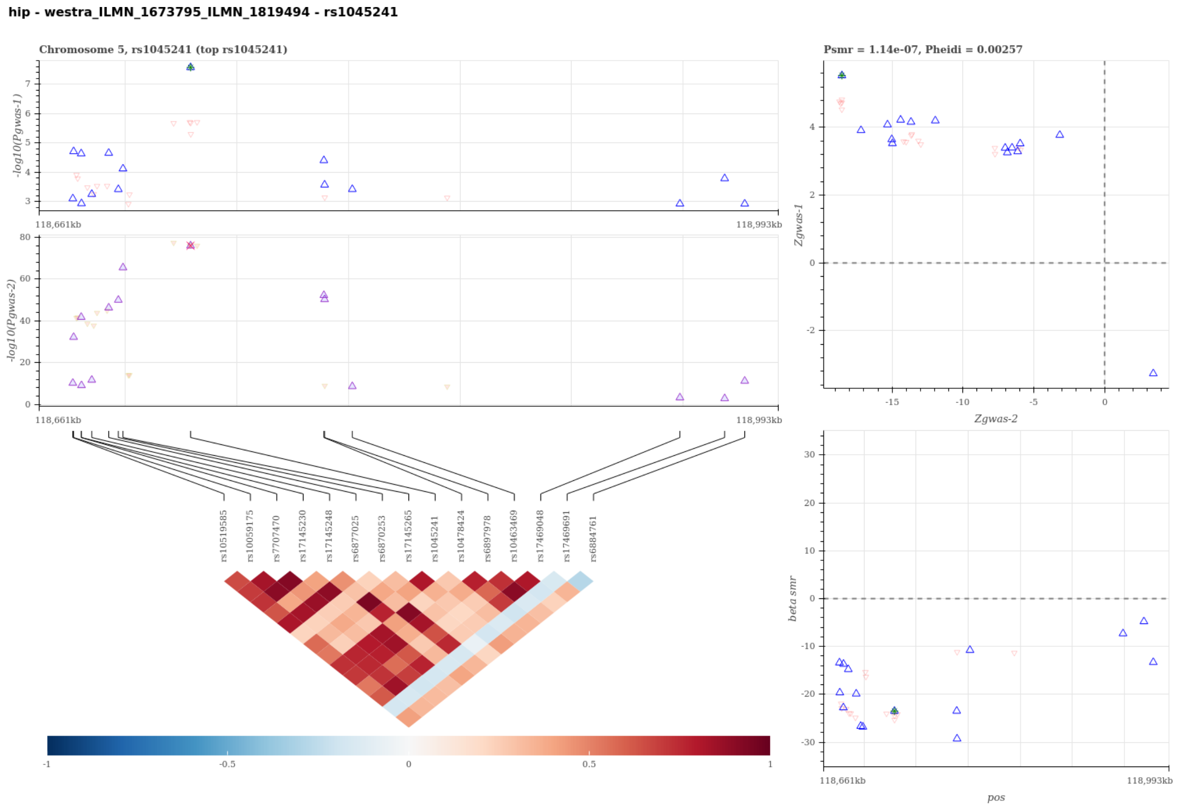


Figure S26: Visualization of the SMR-HEIDI test results to predict candidate gene for HIP using probe ILMN 1673795 ILMN 1819494. A statistical significant SMR test and non-significant HEIDI test suggest that the gene expression of the probe-tagged gene has a cis-eQTL signal (gwas-2) sharing the same single causal variant with HIP (gwas-1). The top left panel compares the regional association results in -log scale p-values between HIP and the eQTL scan. The top right panel compares the regional Z-statistics of the two association scans for the SMR test, and the bottom right panel visualizes the SMR causal effect estimates, i.e. the ratio of the genetic effect on HIP to that on the gene expression tagged by probe ILMN 1673795 ILMN 1819494, for the heterogeneity HEIDI test. The Bottom left panel visualizes the linkage disequilibrium structure of the region. In the plots, each triangle represents a SNP, where the pruned SNP for the SMR-HEIDI test are shown in bigger blue triangles and the top gwas-1 SNP is marked with extra colors.

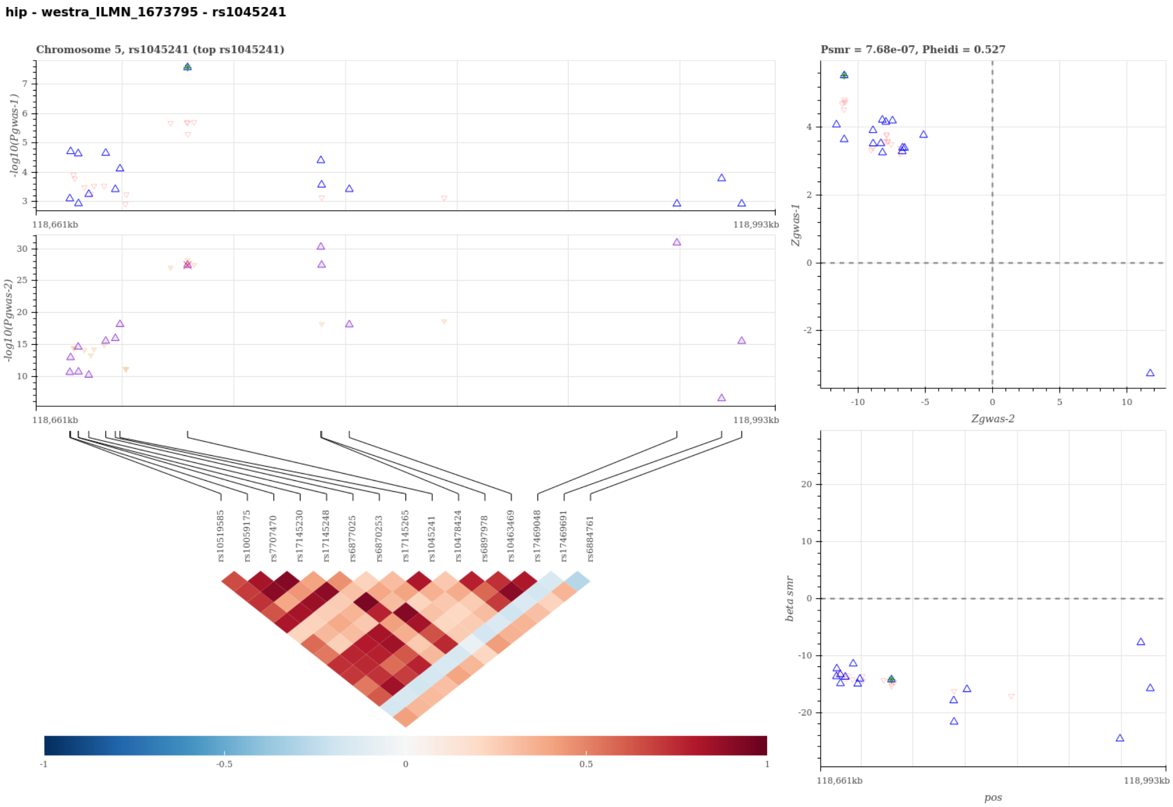


Figure S27: Visualization of the SMR-HEIDI test results to predict candidate gene for HIP using probe ILMN 1673795. A statistical significant SMR test and non-significant HEIDI test suggest that the gene expression of the probe-tagged gene has a cis-eQTL signal (gwas-2) sharing the same single causal variant with HIP (gwas-1). The top left panel compares the regional association results in -log scale p-values between HIP and the eQTL scan. The top right panel compares the regional Z-statistics of the two association scans for the SMR test, and the bottom right panel visualizes the SMR causal effect estimates, i.e. the ratio of the genetic effect on HIP to that on the gene expression tagged by probe ILMN 1673795, for the heterogeneity HEIDI test. The Bottom left panel visualizes the linkage disequilibrium structure of the region. In the plots, each triangle represents a SNP, where the pruned SNP for the SMR-HEIDI test are shown in bigger blue triangles and the top gwas-1 SNP is marked with extra colors.

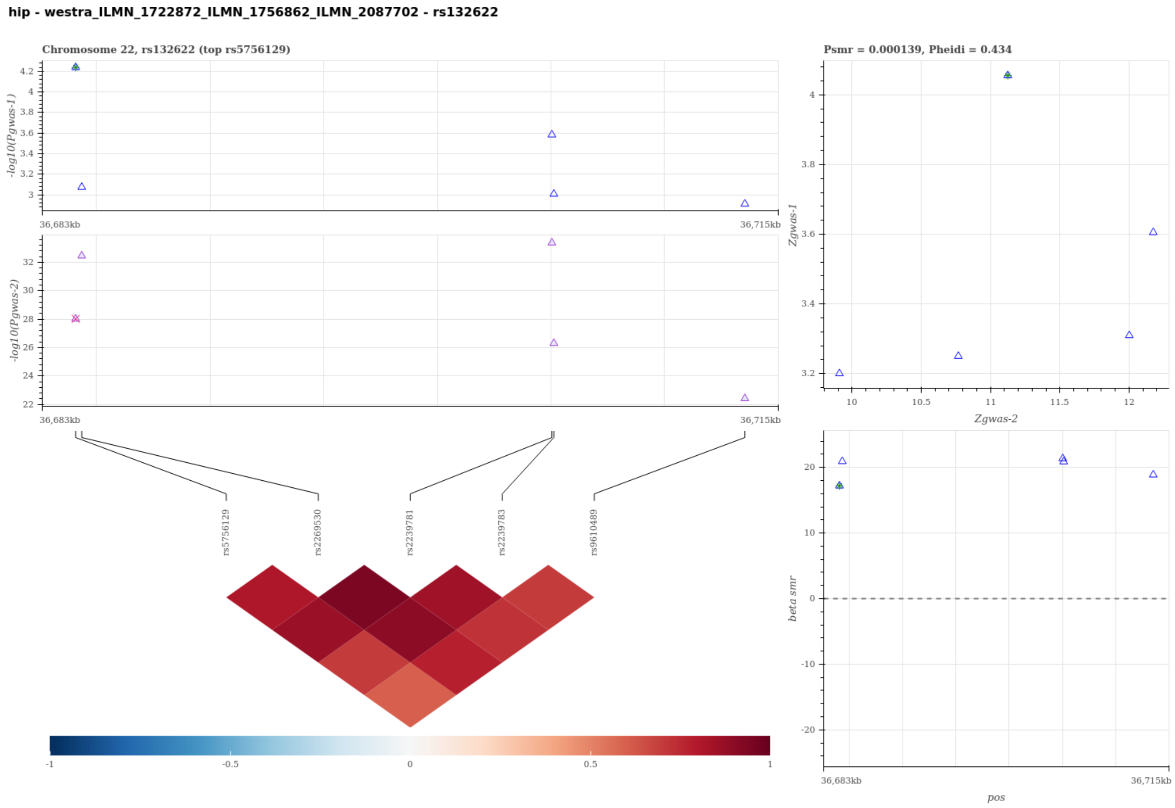


Figure S28: Visualization of the SMR-HEIDI test results to predict candidate gene for HIP using probe ILMN 1722872 ILMN 1756862 ILMN 2087702. A statistical significant SMR test and non-significant HEIDI test suggest that the gene expression of the probe-tagged gene has a cis-eQTL signal (gwas-2) sharing the same single causal variant with HIP (gwas-1). The top left panel compares the regional association results in -log scale p-values between HIP and the eQTL scan. The top right panel compares the regional Z-statistics of the two association scans for the SMR test, and the bottom right panel visualizes the SMR causal effect estimates, i.e. the ratio of the genetic effect on HIP to that on the gene expression tagged by probe ILMN 1722872 ILMN 1756862 ILMN 2087702, for the heterogeneity HEIDI test. The Bottom left panel visualizes the linkage disequilibrium structure of the region. In the plots, each triangle represents a SNP, where the pruned SNP for the SMR-HEIDI test are shown in bigger blue triangles and the top gwas-1 SNP is marked with extra colors.

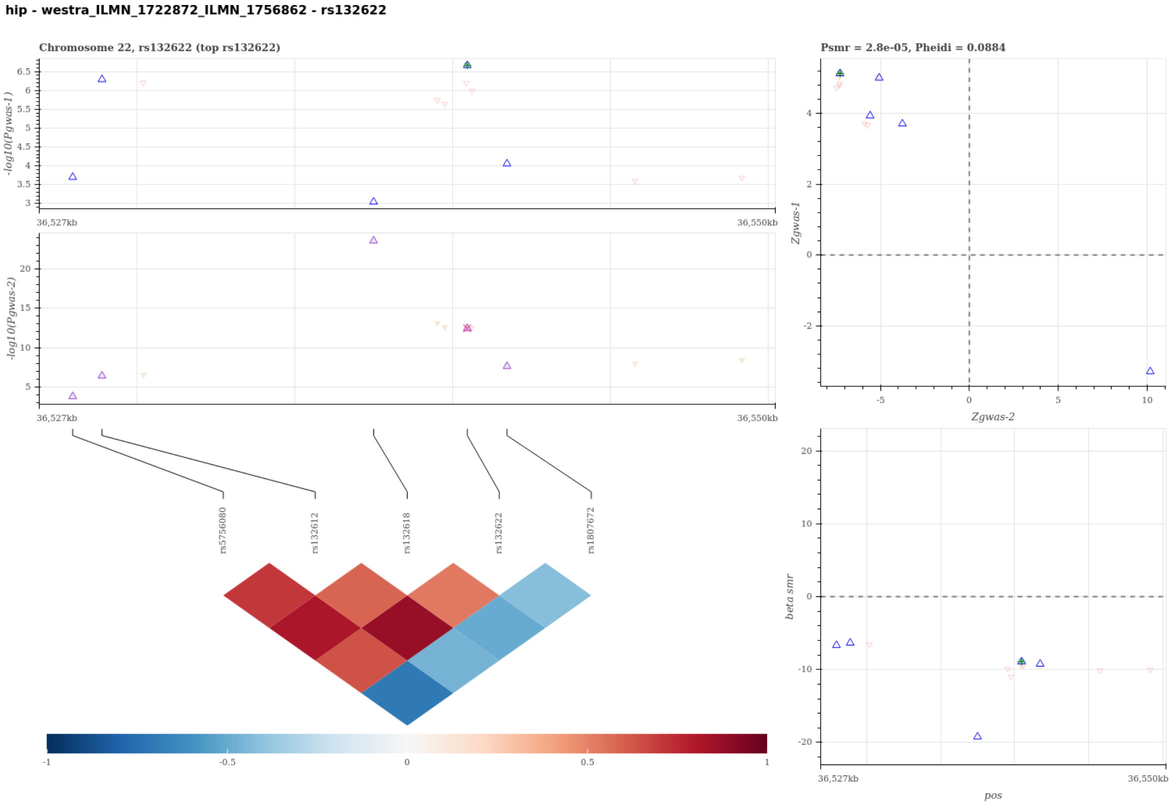


Figure S29: Visualization of the SMR-HEIDI test results to predict candidate gene for HIP using probe ILMN 1722872 ILMN 1756862. A statistical significant SMR test and non-significant HEIDI test suggest that the gene expression of the probe-tagged gene has a cis-eQTL signal (gwas-2) sharing the same single causal variant with HIP (gwas-1). The top left panel compares the regional association results in -log scale p-values between HIP and the eQTL scan. The top right panel compares the regional Z-statistics of the two association scans for the SMR test, and the bottom right panel visualizes the SMR causal effect estimates, i.e. the ratio of the genetic effect on HIP to that on the gene expression tagged by probe ILMN 1722872 ILMN 1756862, for the heterogeneity HEIDI test. The Bottom left panel visualizes the linkage disequilibrium structure of the region. In the plots, each triangle represents a SNP, where the pruned SNP for the SMR-HEIDI test are shown in bigger blue triangles and the top gwas-1 SNP is marked with extra colors.

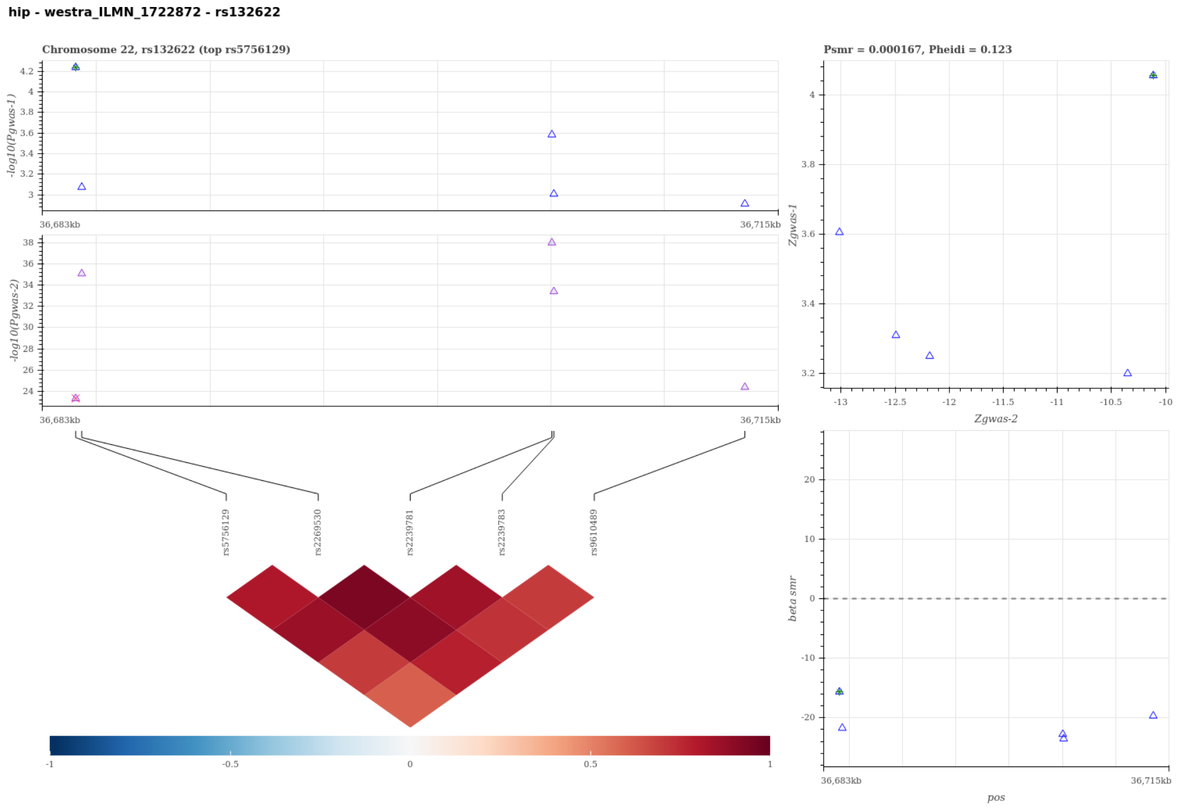


Figure S30: Visualization of the SMR-HEIDI test results to predict candidate gene for HIP using probe ILMN 1722872. A statistical significant SMR test and non-significant HEIDI test suggest that the gene expression of the probe-tagged gene has a cis-eQTL signal (gwas-2) sharing the same single causal variant with HIP (gwas-1). The top left panel compares the regional association results in -log scale p-values between HIP and the eQTL scan. The top right panel compares the regional Z-statistics of the two association scans for the SMR test, and the bottom right panel visualizes the SMR causal effect estimates, i.e. the ratio of the genetic effect on HIP to that on the gene expression tagged by probe ILMN 1722872, for the heterogeneity HEIDI test. The Bottom left panel visualizes the linkage disequilibrium structure of the region. In the plots, each triangle represents a SNP, where the pruned SNP for the SMR-HEIDI test are shown in bigger blue triangles and the top gwas-1 SNP is marked with extra colors.

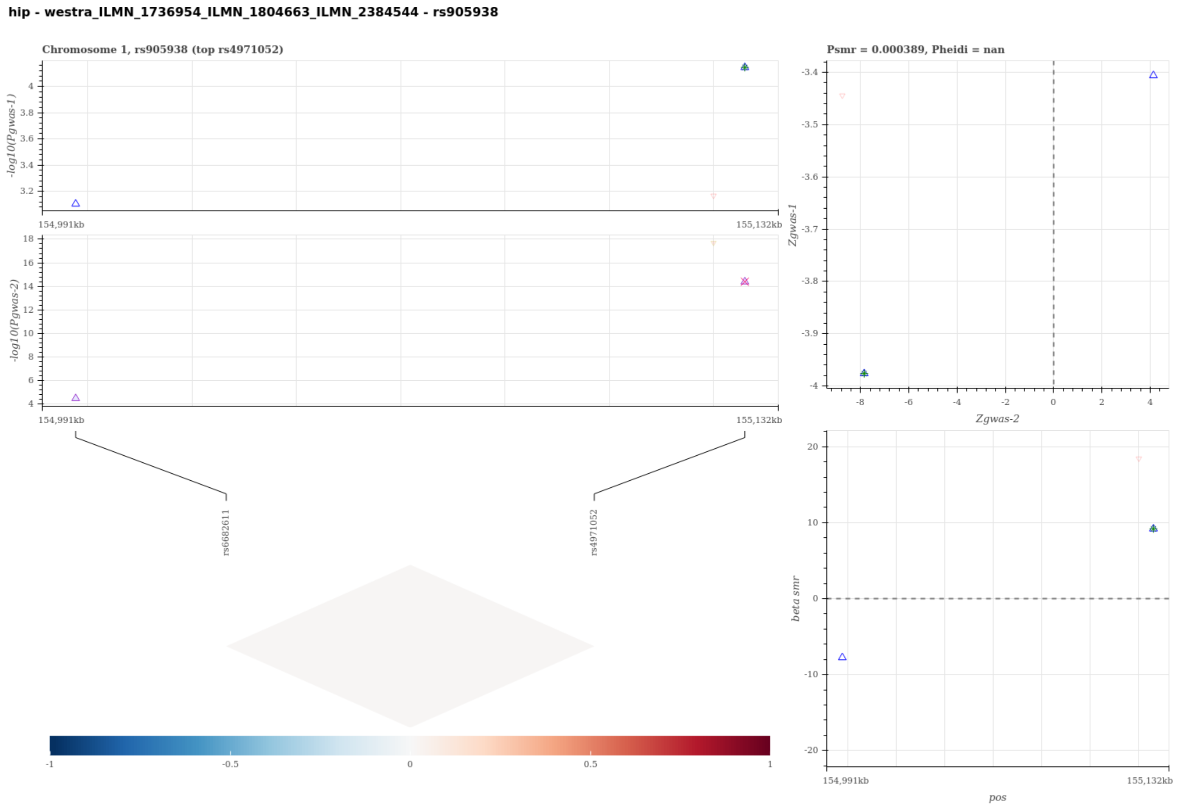


Figure S31: Visualization of the SMR-HEIDI test results to predict candidate gene for HIP using probe ILMN 1736954 ILMN 1804663 ILMN 2384544. A statistical significant SMR test and non-significant HEIDI test suggest that the gene expression of the probe-tagged gene has a cis-eQTL signal (gwas-2) sharing the same single causal variant with HIP (gwas-1). The top left panel compares the regional association results in -log scale p-values between HIP and the eQTL scan. The top right panel compares the regional Z-statistics of the two association scans for the SMR test, and the bottom right panel visualizes the SMR causal effect estimates, i.e. the ratio of the genetic effect on HIP to that on the gene expression tagged by probe ILMN 1736954 ILMN 1804663 ILMN 2384544, for the heterogeneity HEIDI test. The Bottom left panel visualizes the linkage disequilibrium structure of the region. In the plots, each triangle represents a SNP, where the pruned SNP for the SMR-HEIDI test are shown in bigger blue triangles and the top gwas-1 SNP is marked with extra colors.

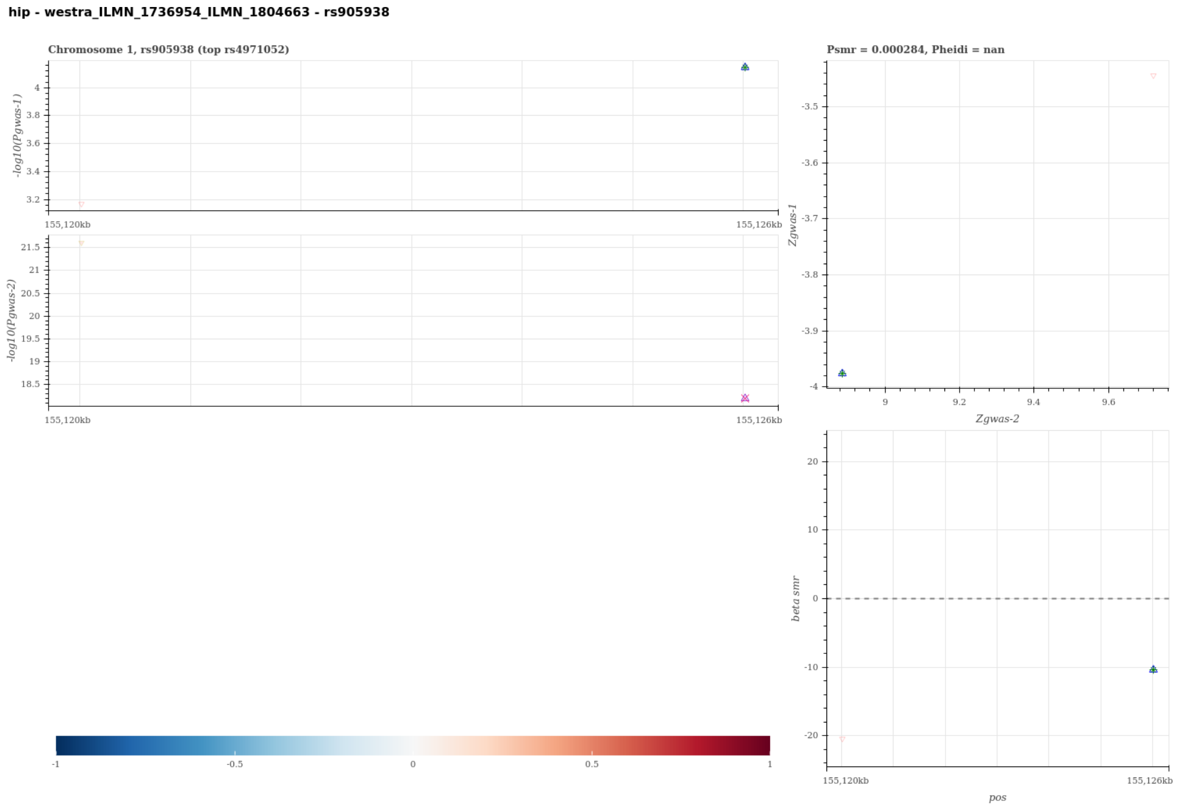


Figure S32: Visualization of the SMR-HEIDI test results to predict candidate gene for HIP using probe ILMN 1736954 ILMN 1804663. A statistical significant SMR test and non-significant HEIDI test suggest that the gene expression of the probe-tagged gene has a cis-eQTL signal (gwas-2) sharing the same single causal variant with HIP (gwas-1). The top left panel compares the regional association results in -log scale p-values between HIP and the eQTL scan. The top right panel compares the regional Z-statistics of the two association scans for the SMR test, and the bottom right panel visualizes the SMR causal effect estimates, i.e. the ratio of the genetic effect on HIP to that on the gene expression tagged by probe ILMN 1736954 ILMN 1804663, for the heterogeneity HEIDI test. The Bottom left panel visualizes the linkage disequilibrium structure of the region. In the plots, each triangle represents a SNP, where the pruned SNP for the SMR-HEIDI test are shown in bigger blue triangles and the top gwas-1 SNP is marked with extra colors.

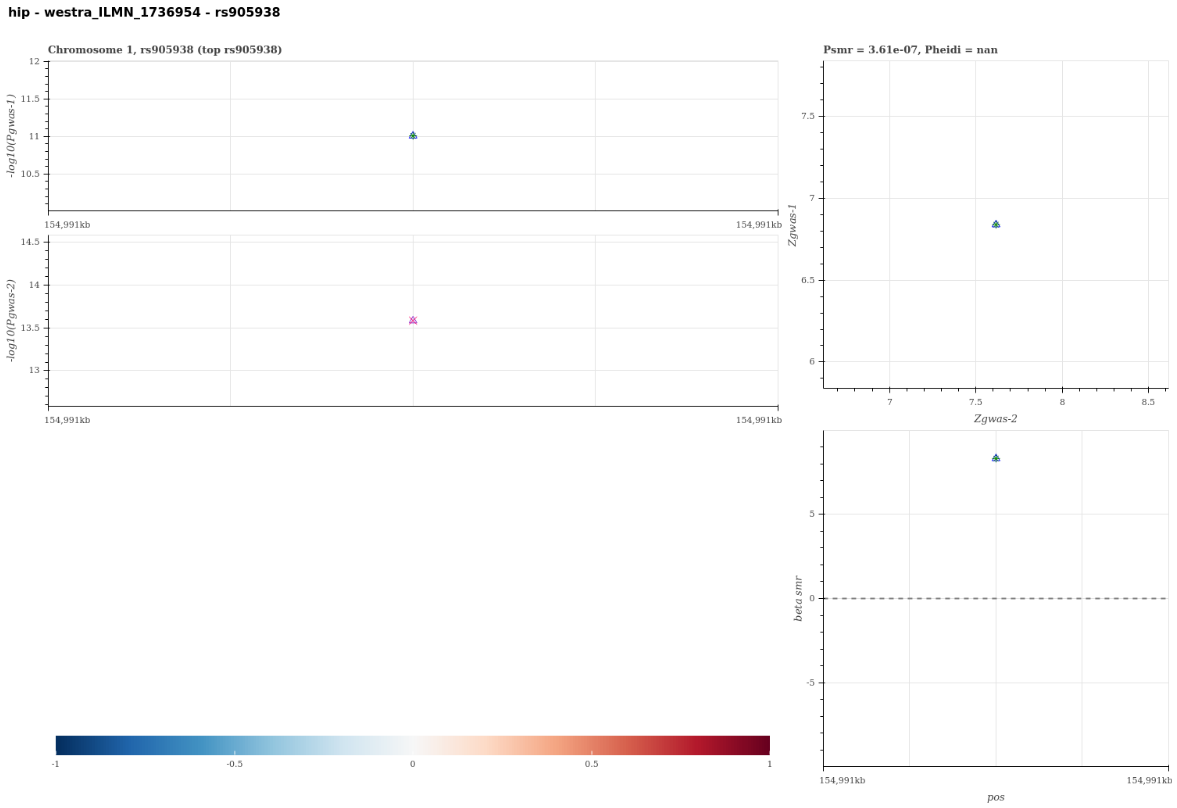


Figure S33: Visualization of the SMR-HEIDI test results to predict candidate gene for HIP using probe ILMN 1736954. A statistical significant SMR test and non-significant HEIDI test suggest that the gene expression of the probe-tagged gene has a cis-eQTL signal (gwas-2) sharing the same single causal variant with HIP (gwas-1). The top left panel compares the regional association results in -log scale p-values between HIP and the eQTL scan. The top right panel compares the regional Z-statistics of the two association scans for the SMR test, and the bottom right panel visualizes the SMR causal effect estimates, i.e. the ratio of the genetic effect on HIP to that on the gene expression tagged by probe ILMN 1736954, for the heterogeneity HEIDI test. The Bottom left panel visualizes the linkage disequilibrium structure of the region. In the plots, each triangle represents a SNP, where the pruned SNP for the SMR-HEIDI test are shown in bigger blue triangles and the top gwas-1 SNP is marked with extra colors.

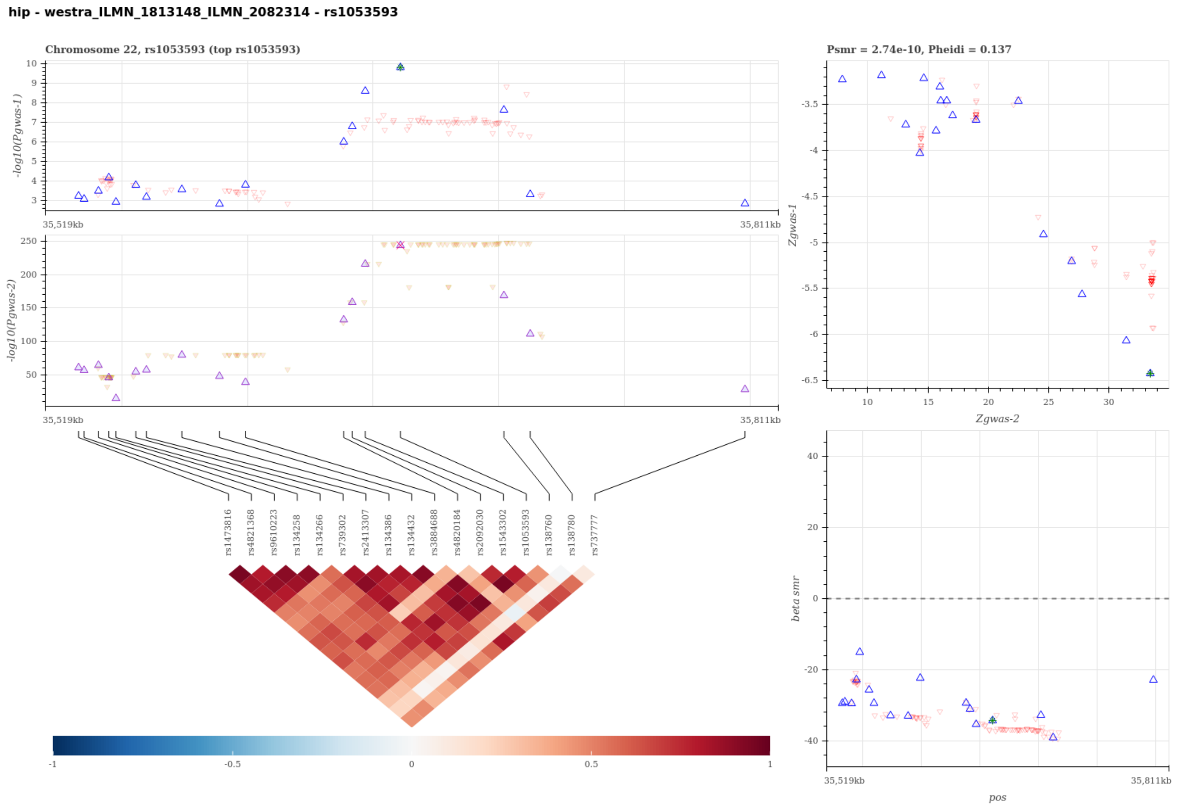


Figure S34: Visualization of the SMR-HEIDI test results to predict candidate gene for HIP using probe ILMN 1813148 ILMN 2082314. A statistical significant SMR test and non-significant HEIDI test suggest that the gene expression of the probe-tagged gene has a cis-eQTL signal (gwas-2) sharing the same single causal variant with HIP (gwas-1). The top left panel compares the regional association results in -log scale p-values between HIP and the eQTL scan. The top right panel compares the regional Z-statistics of the two association scans for the SMR test, and the bottom right panel visualizes the SMR causal effect estimates, i.e. the ratio of the genetic effect on HIP to that on the gene expression tagged by probe ILMN 1813148 ILMN 2082314, for the heterogeneity HEIDI test. The Bottom left panel visualizes the linkage disequilibrium structure of the region. In the plots, each triangle represents a SNP, where the pruned SNP for the SMR-HEIDI test are shown in bigger blue triangles and the top gwas-1 SNP is marked with extra colors.

**hip - westra_ILMN_1813148 - rs1053593**

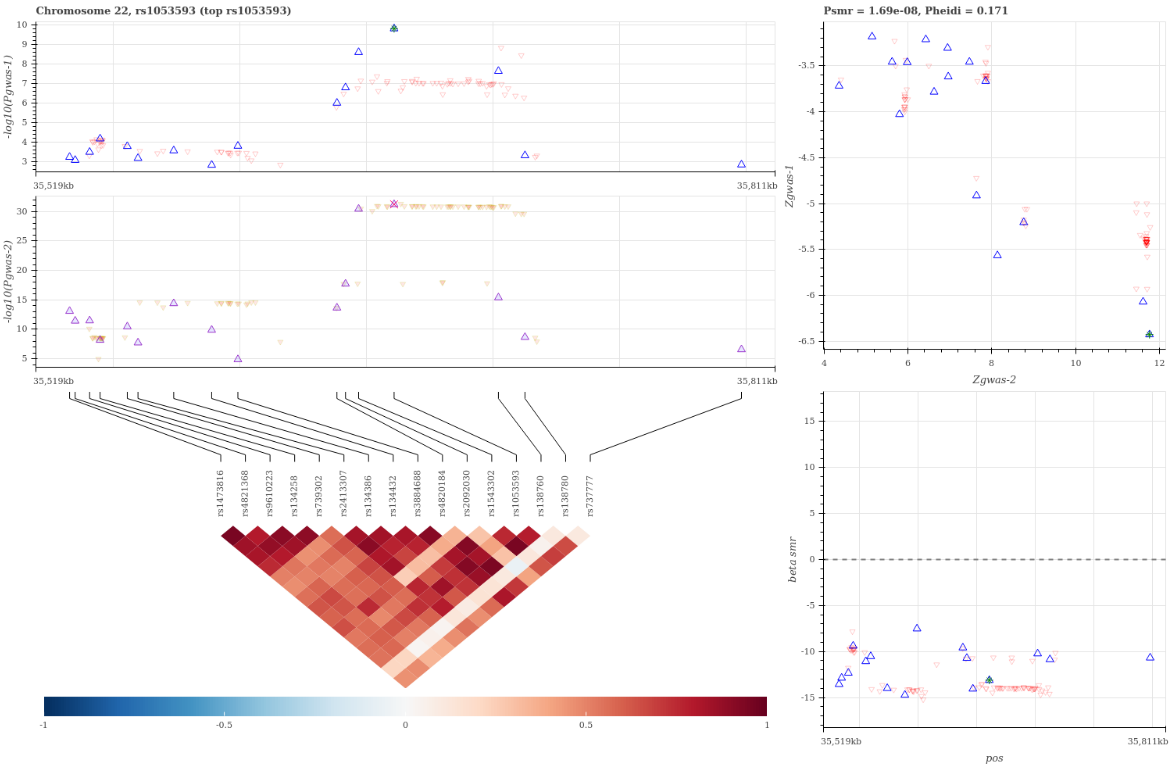


Figure S35: Visualization of the SMR-HEIDI test results to predict candidate gene for HIP using probe ILMN 1813148. A statistical significant SMR test and non-significant HEIDI test suggest that the gene expression of the probe-tagged gene has a cis-eQTL signal (gwas-2) sharing the same single causal variant with HIP (gwas-1). The top left panel compares the regional association results in -log scale p-values between HIP and the eQTL scan. The top right panel compares the regional Z-statistics of the two association scans for the SMR test, and the bottom right panel visualizes the SMR causal effect estimates, i.e. the ratio of the genetic effect on HIP to that on the gene expression tagged by probe ILMN 1813148, for the heterogeneity HEIDI test. The Bottom left panel visualizes the linkage disequilibrium structure of the region. In the plots, each triangle represents a SNP, where the pruned SNP for the SMR-HEIDI test are shown in bigger blue triangles and the top gwas-1 SNP is marked with extra colors.

**wc - westra_ILMN_1669905_ILMN_1758941 - rs459552**

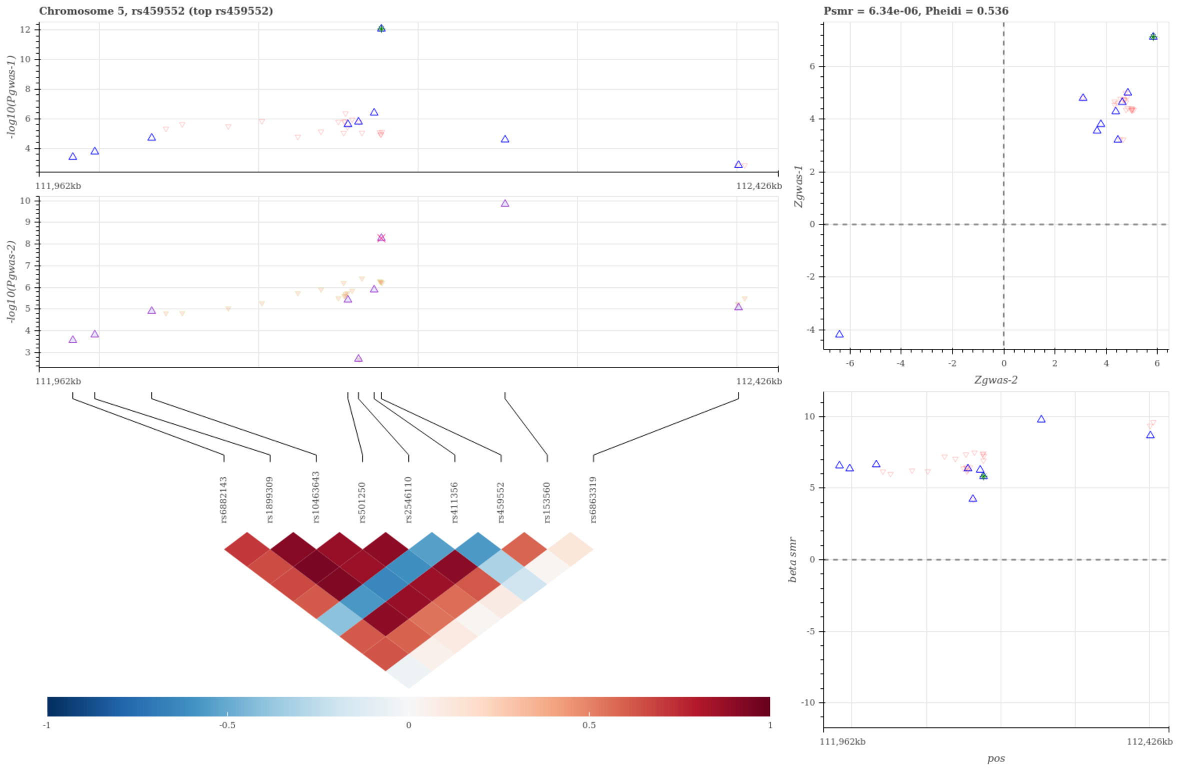


Figure S36: Visualization of the SMR-HEIDI test results to predict candidate gene for WC using probe ILMN 1669905 ILMN 1758941. A statistical significant SMR test and non-significant HEIDI test suggest that the gene expression of the probe-tagged gene has a cis-eQTL signal (gwas-2) sharing the same single causal variant with WC (gwas-1). The top left panel compares the regional association results in -log scale p-values between WC and the eQTL scan. The top right panel compares the regional Z-statistics of the two association scans for the SMR test, and the bottom right panel visualizes the SMR causal effect estimates, i.e. the ratio of the genetic effect on WC to that on the gene expression tagged by probe ILMN 1669905 ILMN 1758941, for the heterogeneity HEIDI test. The Bottom left panel visualizes the linkage disequilibrium structure of the region. In the plots, each triangle represents a SNP, where the pruned SNP for the SMR-HEIDI test are shown in bigger blue triangles and the top gwas-1 SNP is marked with extra colors.

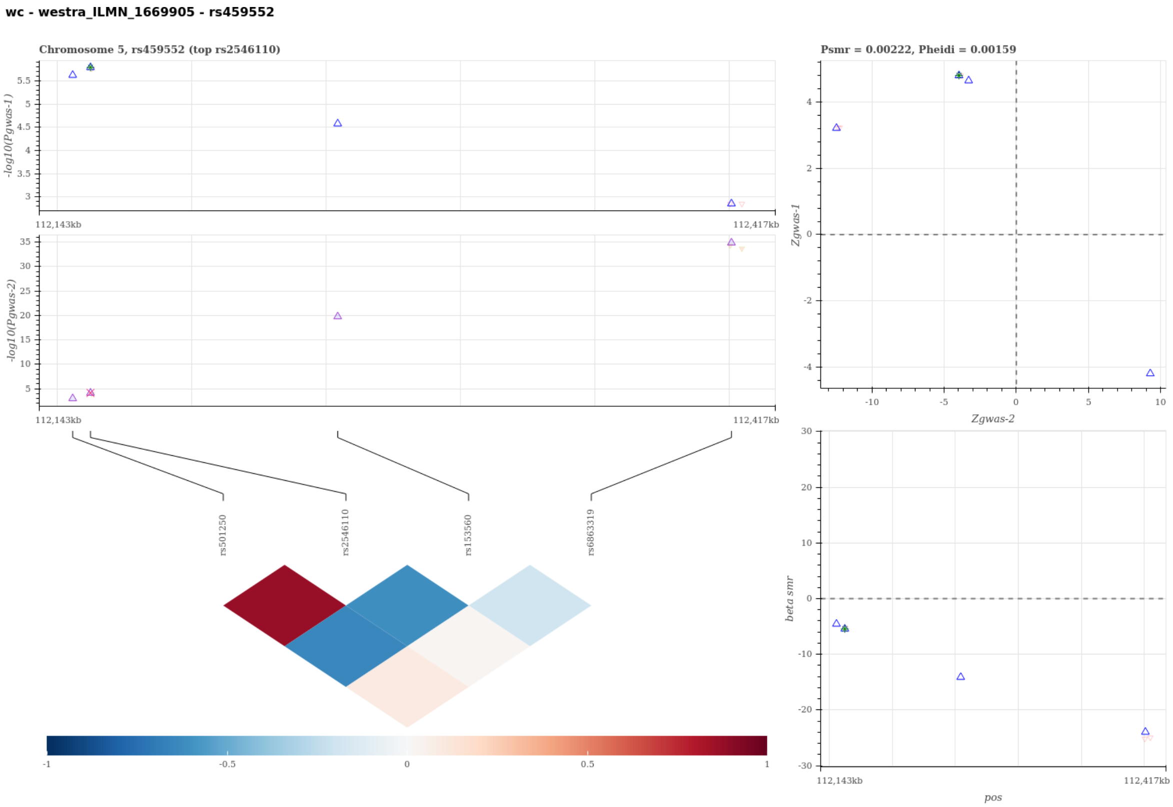


Figure S37: Visualization of the SMR-HEIDI test results to predict candidate gene for WC using probe ILMN 1669905. A statistical significant SMR test and non-significant HEIDI test suggest that the gene expression of the probe-tagged gene has a cis-eQTL signal (gwas-2) sharing the same single causal variant with WC (gwas-1). The top left panel compares the regional association results in -log scale p-values between WC and the eQTL scan. The top right panel compares the regional Z-statistics of the two association scans for the SMR test, and the bottom right panel visualizes the SMR causal effect estimates, i.e. the ratio of the genetic effect on WC to that on the gene expression tagged by probe ILMN 1669905, for the heterogeneity HEIDI test. The Bottom left panel visualizes the linkage disequilibrium structure of the region. In the plots, each triangle represents a SNP, where the pruned SNP for the SMR-HEIDI test are shown in bigger blue triangles and the top gwas-1 SNP is marked with extra colors.

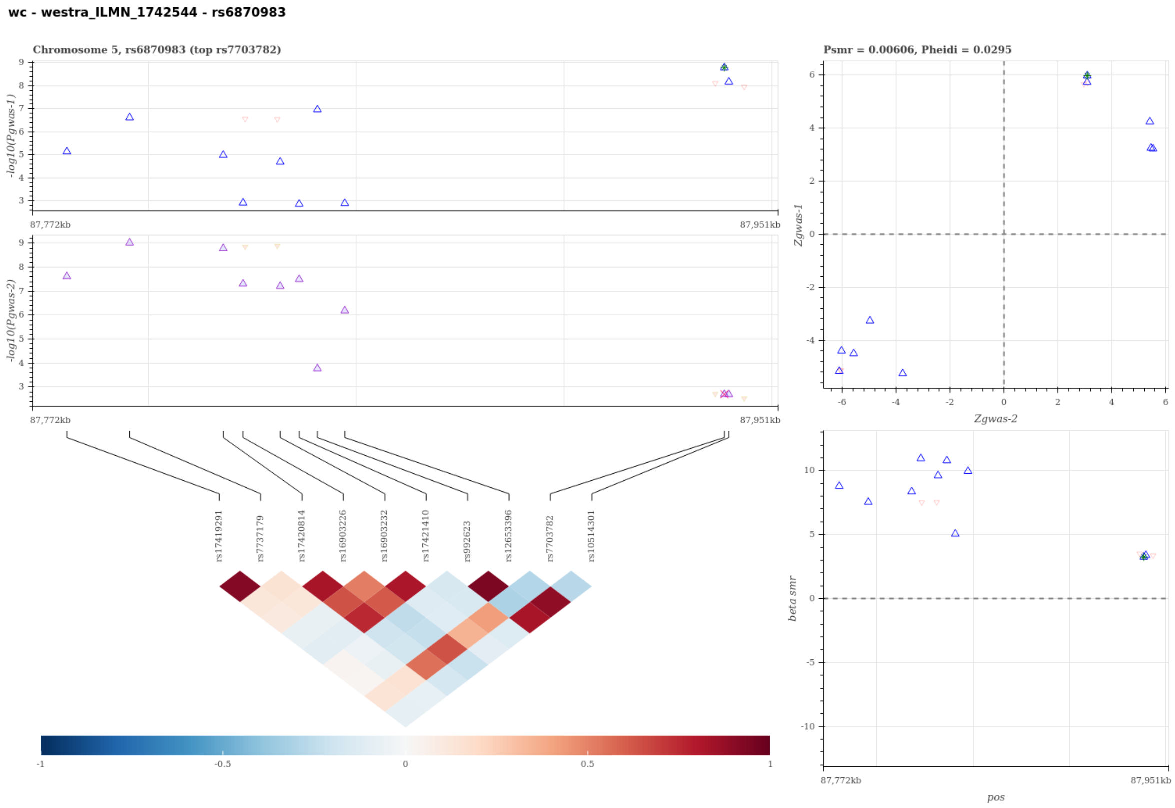


Figure S38: Visualization of the SMR-HEIDI test results to predict candidate gene for WC using probe ILMN 1742544. A statistical significant SMR test and non-significant HEIDI test suggest that the gene expression of the probe-tagged gene has a cis-eQTL signal (gwas-2) sharing the same single causal variant with WC (gwas-1). The top left panel compares the regional association results in -log scale p-values between WC and the eQTL scan. The top right panel compares the regional Z-statistics of the two association scans for the SMR test, and the bottom right panel visualizes the SMR causal effect estimates, i.e. the ratio of the genetic effect on WC to that on the gene expression tagged by probe ILMN 1742544, for the heterogeneity HEIDI test. The Bottom left panel visualizes the linkage disequilibrium structure of the region. In the plots, each triangle represents a SNP, where the pruned SNP for the SMR-HEIDI test are shown in bigger blue triangles and the top gwas-1 SNP is marked with extra colors.

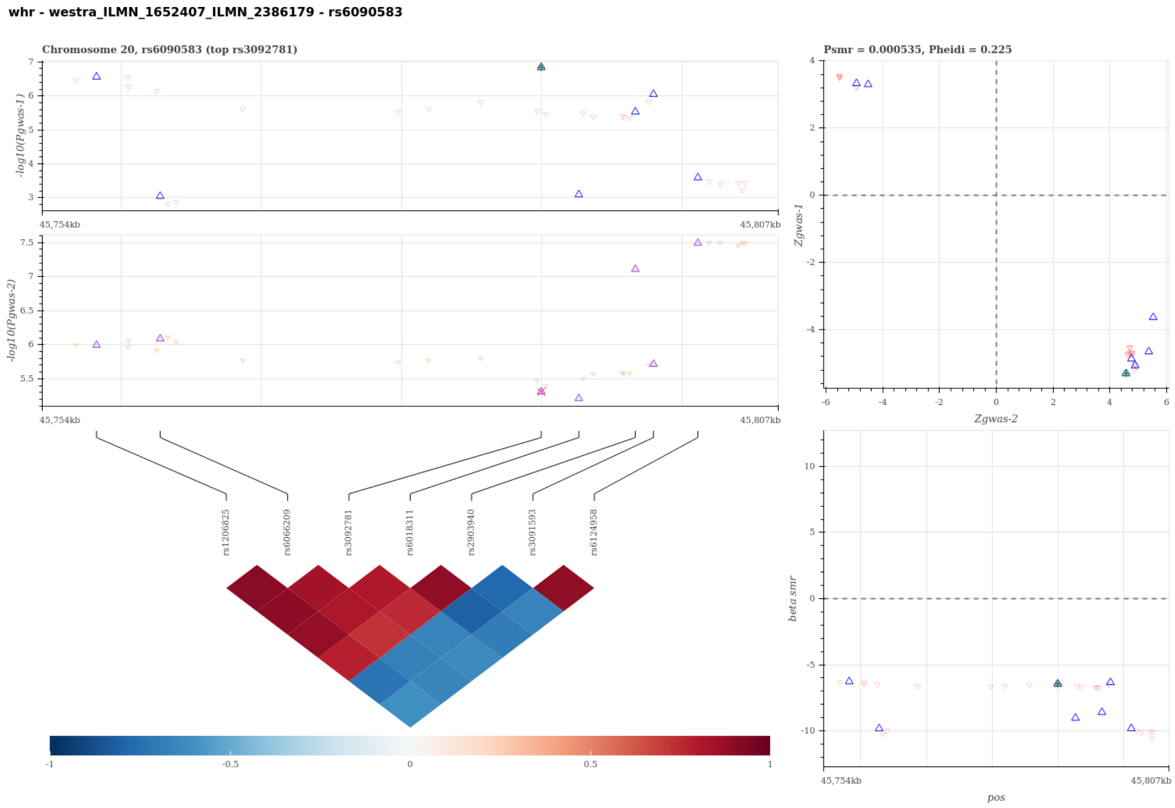


Figure S39: Visualization of the SMR-HEIDI test results to predict candidate gene for WHR using probe ILMN 1652407 ILMN 2386179. A statistical significant SMR test and non-significant HEIDI test suggest that the gene expression of the probe-tagged gene has a cis-eQTL signal (gwas-2) sharing the same single causal variant with WHR (gwas-1). The top left panel compares the regional association results in -log scale p-values between WHR and the eQTL scan. The top right panel compares the regional Z-statistics of the two association scans for the SMR test, and the bottom right panel visualizes the SMR causal effect estimates, i.e. the ratio of the genetic effect on WHR to that on the gene expression tagged by probe ILMN 1652407 ILMN 2386179, for the heterogeneity HEIDI test. The Bottom left panel visualizes the linkage disequilibrium structure of the region. In the plots, each triangle represents a SNP, where the pruned SNP for the SMR-HEIDI test are shown in bigger blue triangles and the top gwas-1 SNP is marked with extra colors.

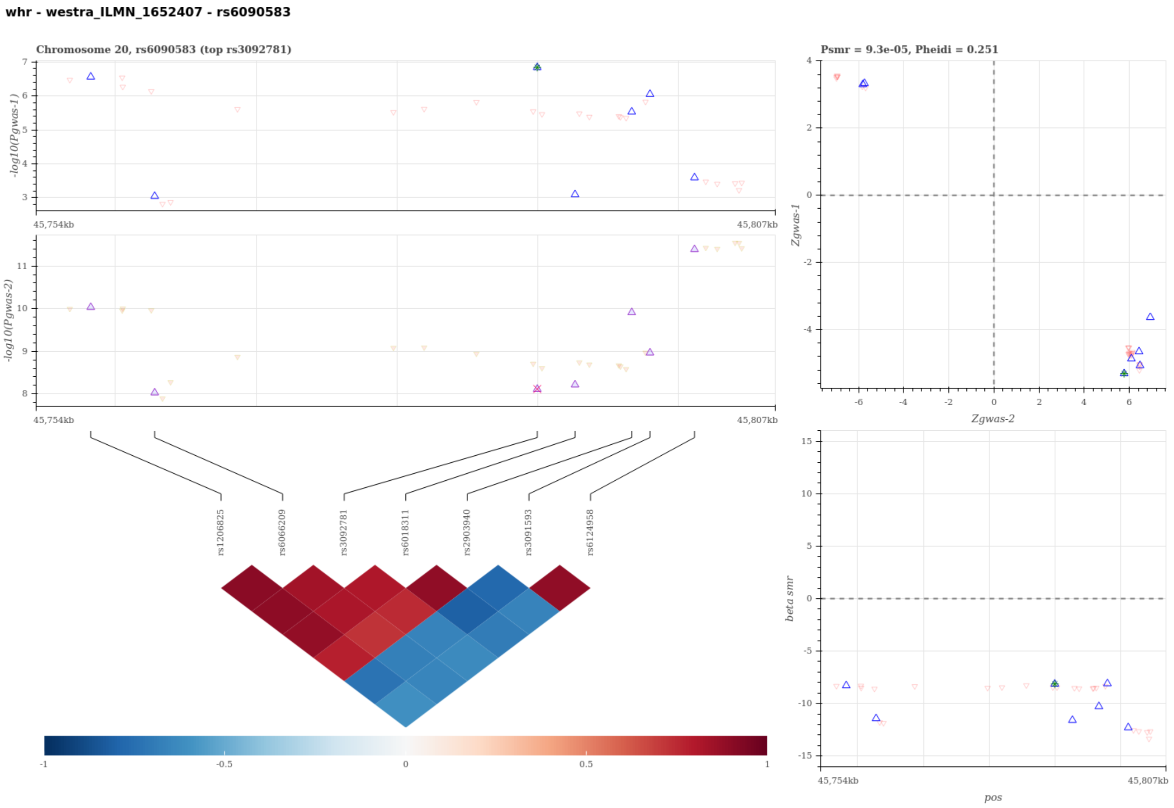


Figure S40: Visualization of the SMR-HEIDI test results to predict candidate gene for WHR using probe ILMN 1652407. A statistical significant SMR test and non-significant HEIDI test suggest that the gene expression of the probe-tagged gene has a cis-eQTL signal (gwas-2) sharing the same single causal variant with WHR (gwas-1). The top left panel compares the regional association results in -log scale p-values between WHR and the eQTL scan. The top right panel compares the regional Z-statistics of the two association scans for the SMR test, and the bottom right panel visualizes the SMR causal effect estimates, i.e. the ratio of the genetic effect on WHR to that on the gene expression tagged by probe ILMN 1652407, for the heterogeneity HEIDI test. The Bottom left panel visualizes the linkage disequilibrium structure of the region. In the plots, each triangle represents a SNP, where the pruned SNP for the SMR-HEIDI test are shown in bigger blue triangles and the top gwas-1 SNP is marked with extra colors.

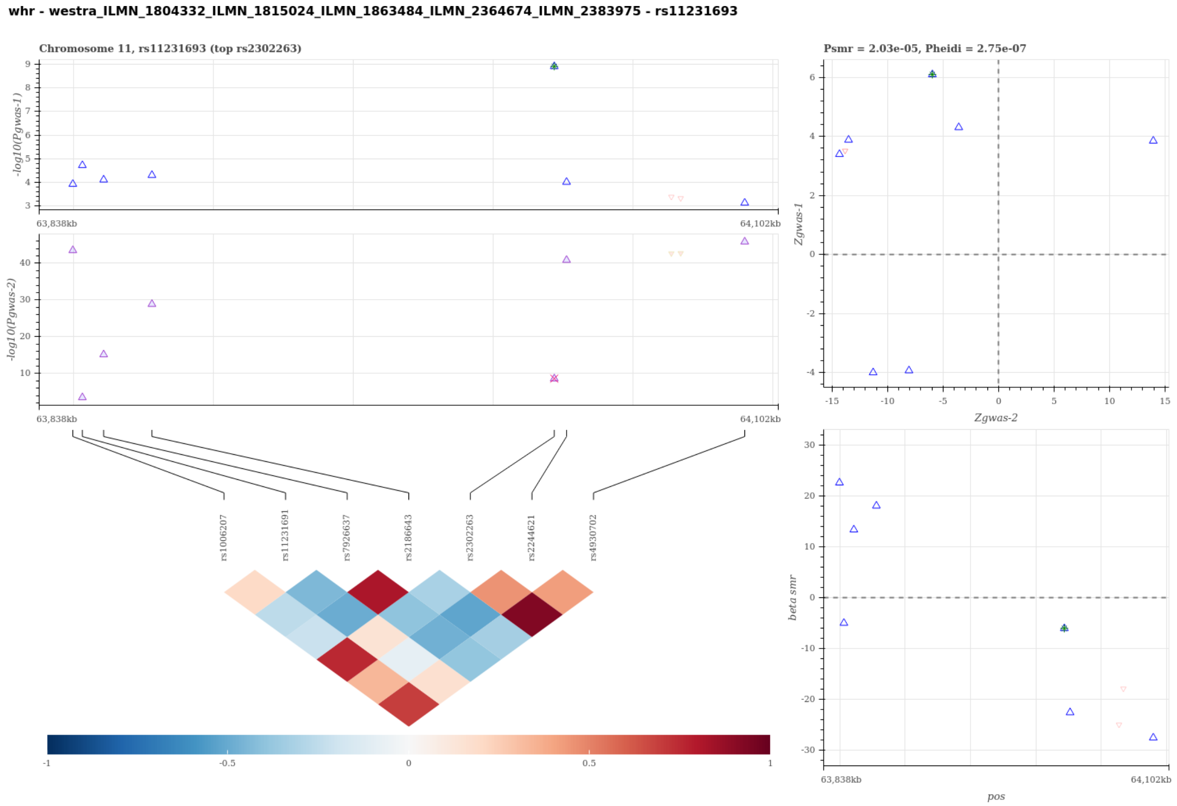


Figure S41: Visualization of the SMR-HEIDI test results to predict candidate gene for WHR using probe ILMN 1804332 ILMN 1815024 ILMN 1863484 ILMN 2364674 ILMN 2383975. A statistical significant SMR test and non-significant HEIDI test suggest that the gene expression of the probe-tagged gene has a cis-eQTL signal (gwas-2) sharing the same single causal variant with WHR (gwas-1). The top left panel compares the regional association results in -log scale p-values between WHR and the eQTL scan. The top right panel compares the regional Z-statistics of the two association scans for the SMR test, and the bottom right panel visualizes the SMR causal effect estimates, i.e. the ratio of the genetic effect on WHR to that on the gene expression tagged by probe ILMN 1804332 ILMN 1815024 ILMN 1863484 ILMN 2364674 ILMN 2383975, for the heterogeneity HEIDI test. The Bottom left panel visualizes the linkage disequilibrium structure of the region. In the plots, each triangle represents a SNP, where the pruned SNP for the SMR-HEIDI test are shown in bigger blue triangles and the top gwas-1 SNP is marked with extra colors.

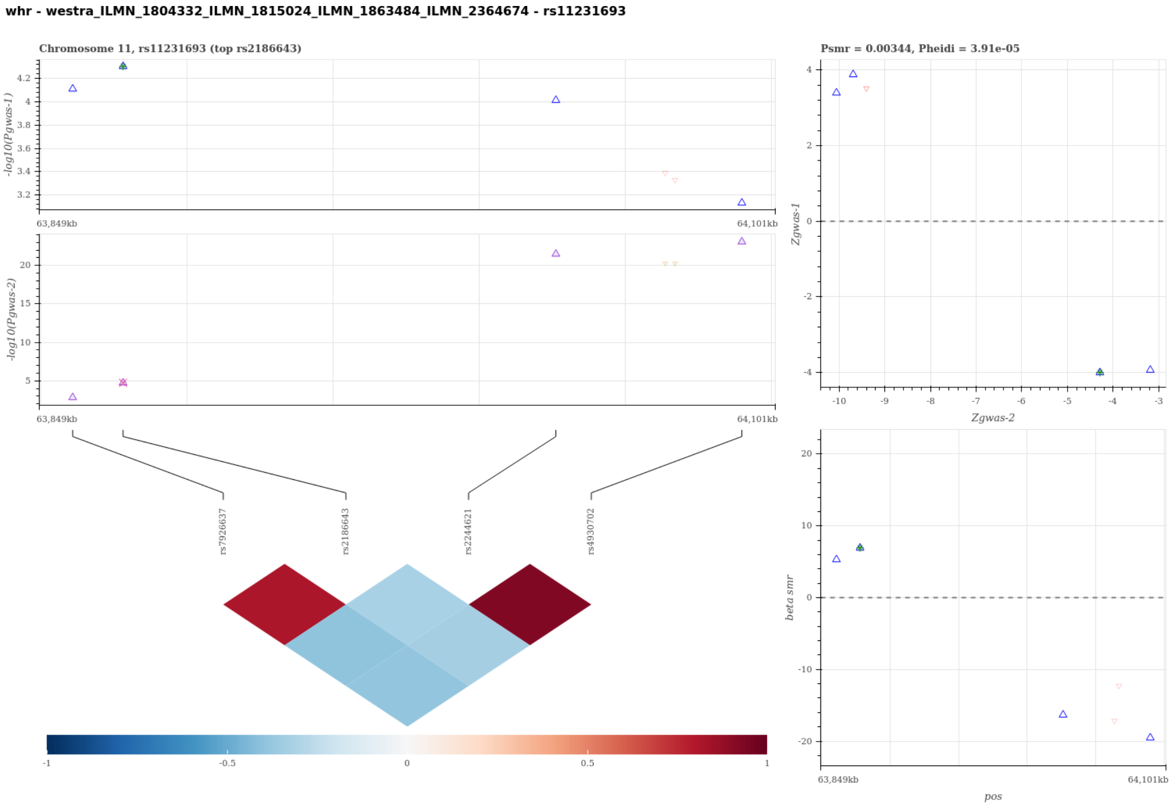


Figure S42: Visualization of the SMR-HEIDI test results to predict candidate gene for WHR using probe ILMN 1804332 ILMN 1815024 ILMN 1863484 ILMN 2364674. A statistical significant SMR test and non-significant HEIDI test suggest that the gene expression of the probe-tagged gene has a cis-eQTL signal (gwas-2) sharing the same single causal variant with WHR (gwas-1). The top left panel compares the regional association results in -log scale p-values between WHR and the eQTL scan. The top right panel compares the regional Z-statistics of the two association scans for the SMR test, and the bottom right panel visualizes the SMR causal effect estimates, i.e. the ratio of the genetic effect on WHR to that on the gene expression tagged by probe ILMN 1804332 ILMN 1815024 ILMN 1863484 ILMN 2364674, for the heterogeneity HEIDI test. The Bottom left panel visualizes the linkage disequilibrium structure of the region. In the plots, each triangle represents a SNP, where the pruned SNP for the SMR-HEIDI test are shown in bigger blue triangles and the top gwas-1 SNP is marked with extra colors.

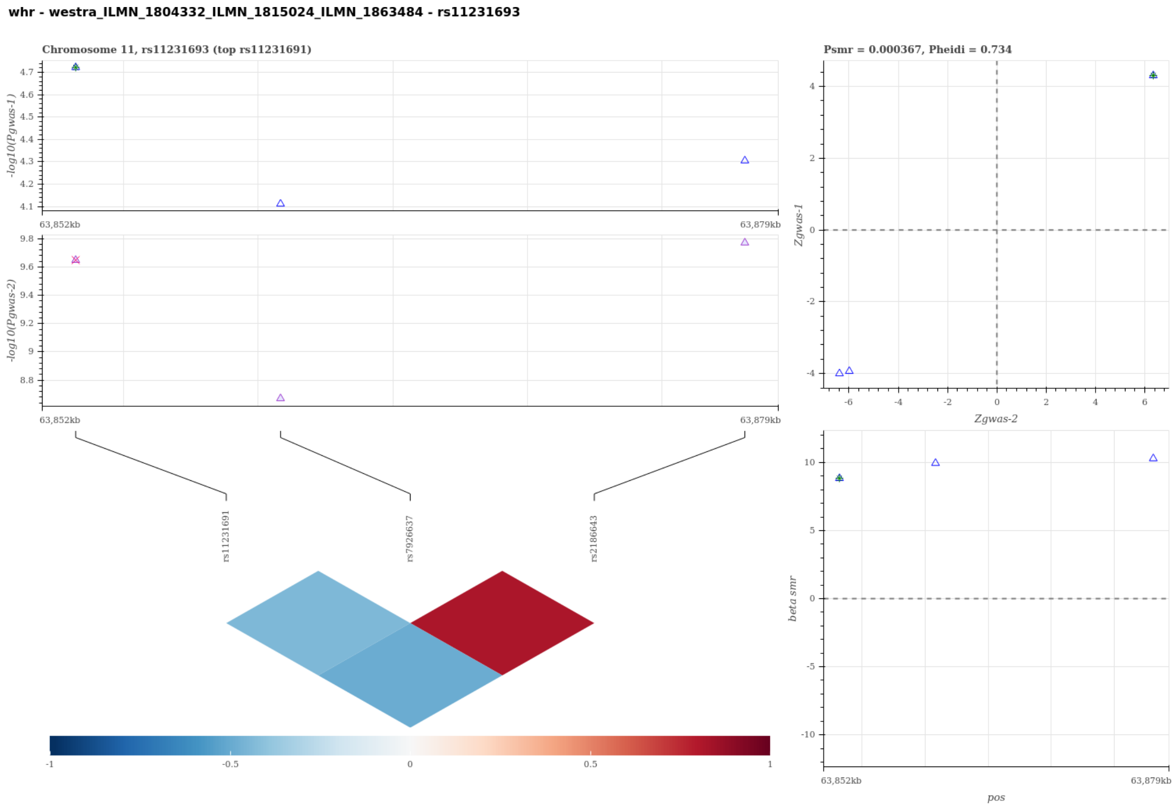


Figure S43: Visualization of the SMR-HEIDI test results to predict candidate gene for WHR using probe ILMN 1804332 ILMN 1815024 ILMN 1863484. A statistical significant SMR test and non-significant HEIDI test suggest that the gene expression of the probe-tagged gene has a cis-eQTL signal (gwas-2) sharing the same single causal variant with WHR (gwas-1). The top left panel compares the regional association results in -log scale p-values between WHR and the eQTL scan. The top right panel compares the regional Z-statistics of the two association scans for the SMR test, and the bottom right panel visualizes the SMR causal effect estimates, i.e. the ratio of the genetic effect on WHR to that on the gene expression tagged by probe ILMN 1804332 ILMN 1815024 ILMN 1863484, for the heterogeneity HEIDI test. The Bottom left panel visualizes the linkage disequilibrium structure of the region. In the plots, each triangle represents a SNP, where the pruned SNP for the SMR-HEIDI test are shown in bigger blue triangles and the top gwas-1 SNP is marked with extra colors.

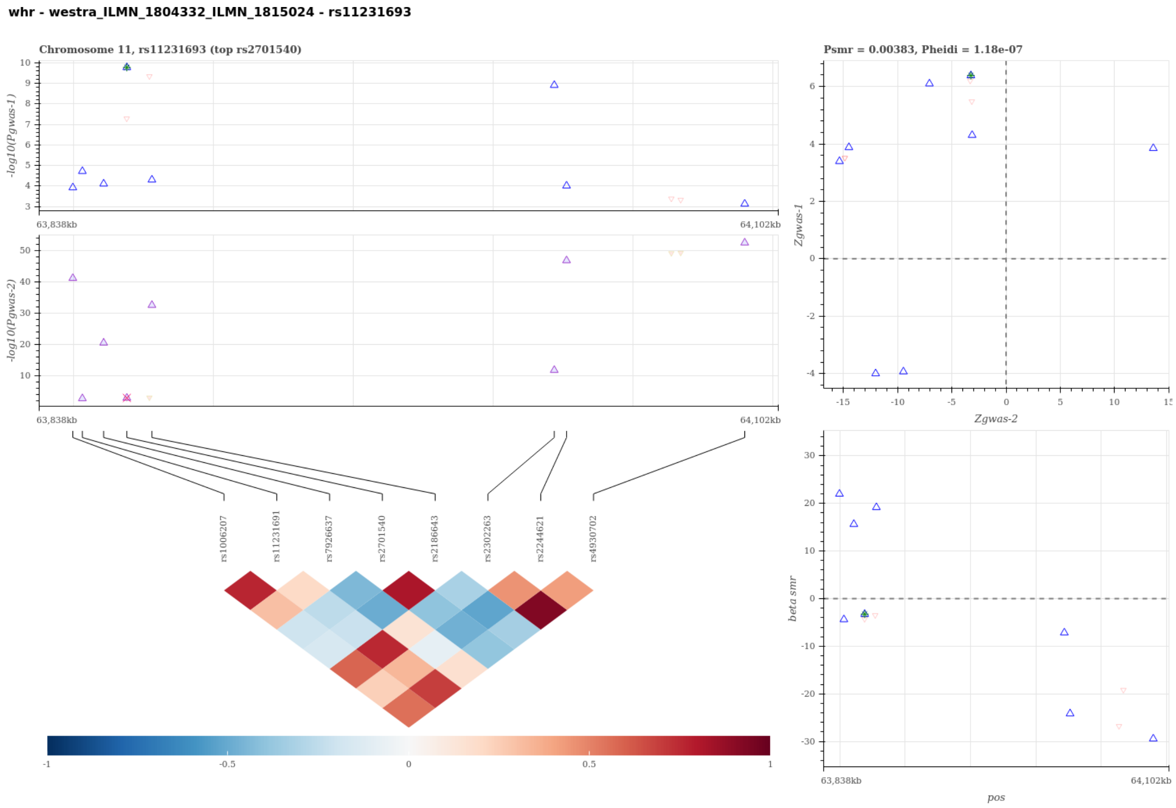


Figure S44: Visualization of the SMR-HEIDI test results to predict candidate gene for WHR using probe ILMN 1804332 ILMN 1815024. A statistical significant SMR test and non-significant HEIDI test suggest that the gene expression of the probe-tagged gene has a cis-eQTL signal (gwas-2) sharing the same single causal variant with WHR (gwas-1). The top left panel compares the regional association results in -log scale p-values between WHR and the eQTL scan. The top right panel compares the regional Z-statistics of the two association scans for the SMR test, and the bottom right panel visualizes the SMR causal effect estimates, i.e. the ratio of the genetic effect on WHR to that on the gene expression tagged by probe ILMN 1804332 ILMN 1815024, for the heterogeneity HEIDI test. The Bottom left panel visualizes the linkage disequilibrium structure of the region. In the plots, each triangle represents a SNP, where the pruned SNP for the SMR-HEIDI test are shown in bigger blue triangles and the top gwas-1 SNP is marked with extra colors.

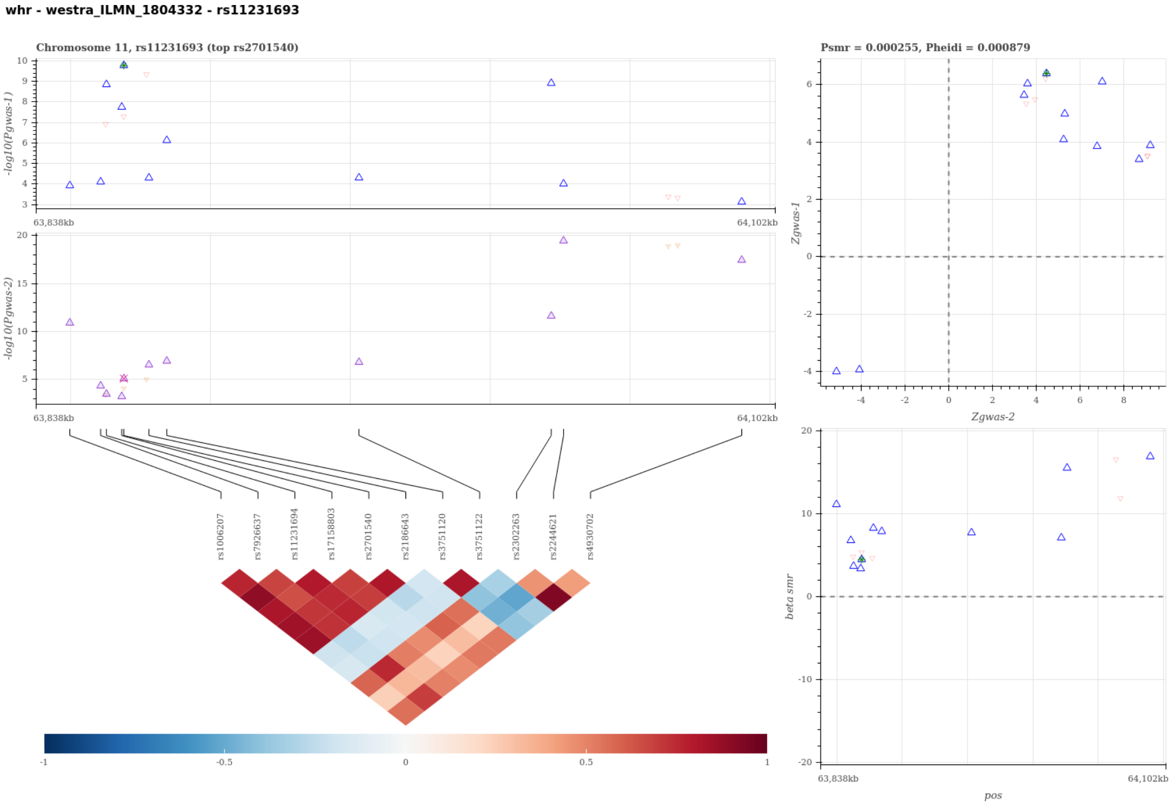


Figure S45: Visualization of the SMR-HEIDI test results to predict candidate gene for WHR using probe ILMN 1804332. A statistical significant SMR test and non-significant HEIDI test suggest that the gene expression of the probe-tagged gene has a cis-eQTL signal (gwas-2) sharing the same single causal variant with WHR (gwas-1). The top left panel compares the regional association results in -log scale p-values between WHR and the eQTL scan. The top right panel compares the regional Z-statistics of the two association scans for the SMR test, and the bottom right panel visualizes the SMR causal effect estimates, i.e. the ratio of the genetic effect on WHR to that on the gene expression tagged by probe ILMN 1804332, for the heterogeneity HEIDI test. The Bottom left panel visualizes the linkage disequilibrium structure of the region. In the plots, each triangle represents a SNP, where the pruned SNP for the SMR-HEIDI test are shown in bigger blue triangles and the top gwas-1 SNP is marked with extra colors.

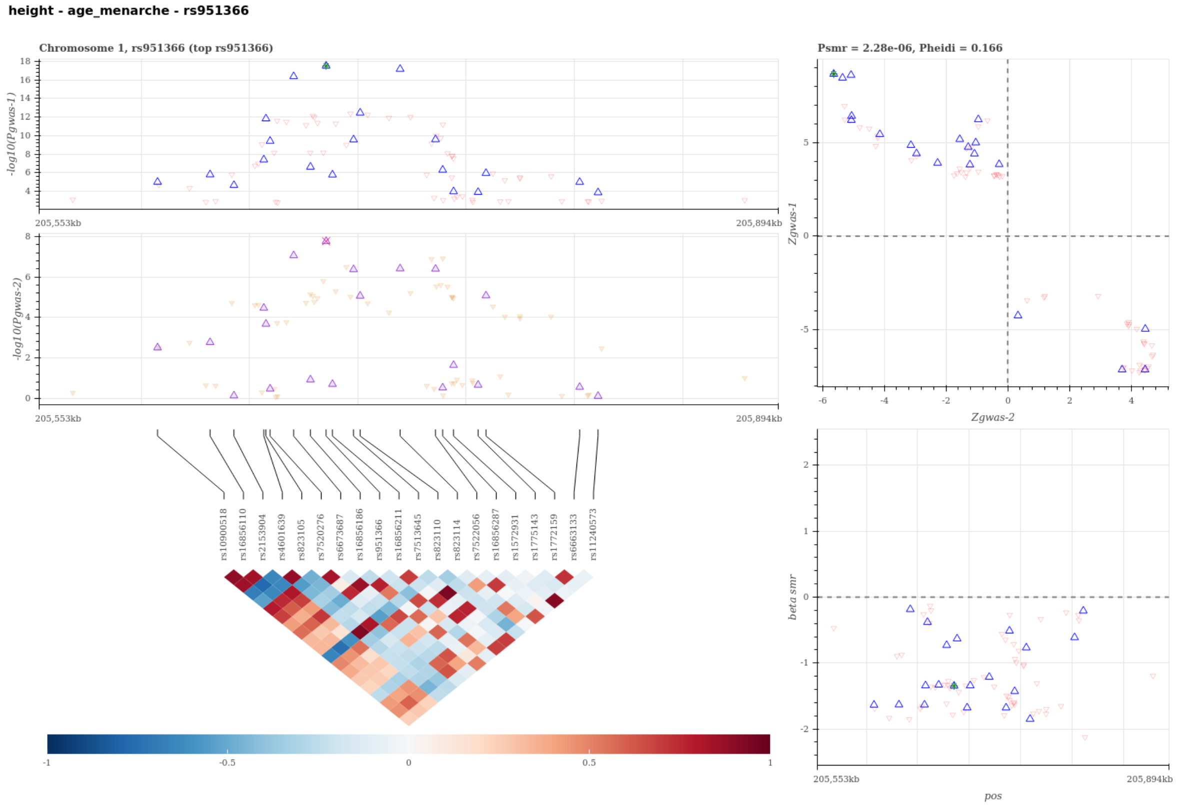


Figure S46: Visualization of the SMR-HEIDI test results for a single shared causal variant between HEIGHT and AgeAtMenarche. A statistical significant SMR test and non-significant HEIDI test suggest that AgeAtMenarche (gwas-2) shares the same single causal variant with HEIGHT (gwas-1). The top left panel compares the regional association results in -log scale p-values between HEIGHT and AgeAtMenarche. The top right panel compares the regional Z-statistics of the two association scans for the SMR test, and the bottom right panel visualizes the SMR causal effect estimates, i.e. the ratio of the genetic effect on HEIGHT to that on AgeAtMenarche, for the heterogeneity HEIDI test. The Bottom left panel visualizes the linkage disequilibrium structure of the region. In the plots, each triangle represents a SNP, where the pruned SNP for the SMR-HEIDI test are shown in bigger blue triangles and the top gwas-1 SNP is marked with extra colors.

**height - bw - rs11708067**

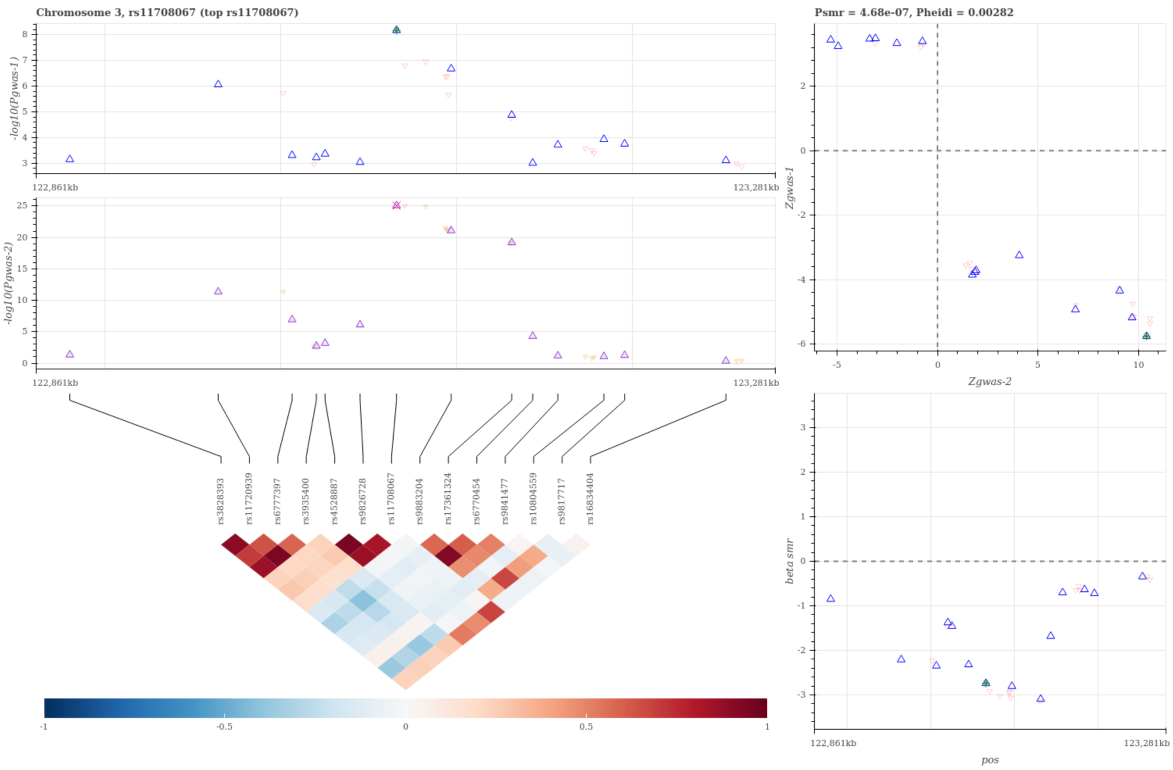


Figure S47: Visualization of the SMR-HEIDI test results for a single shared causal variant between HEIGHT and BirthWeight. A statistical significant SMR test and non-significant HEIDI test suggest that BirthWeight (gwas-2) shares the same single causal variant with HEIGHT (gwas-1). The top left panel compares the regional association results in -log scale p-values between HEIGHT and BirthWeight. The top right panel compares the regional Z-statistics of the two association scans for the SMR test, and the bottom right panel visualizes the SMR causal effect estimates, i.e. the ratio of the genetic effect on HEIGHT to that on BirthWeight, for the heterogeneity HEIDI test. The Bottom left panel visualizes the linkage disequilibrium structure of the region. In the plots, each triangle represents a SNP, where the pruned SNP for the SMR-HEIDI test are shown in bigger blue triangles and the top gwas-1 SNP is marked with extra colors.

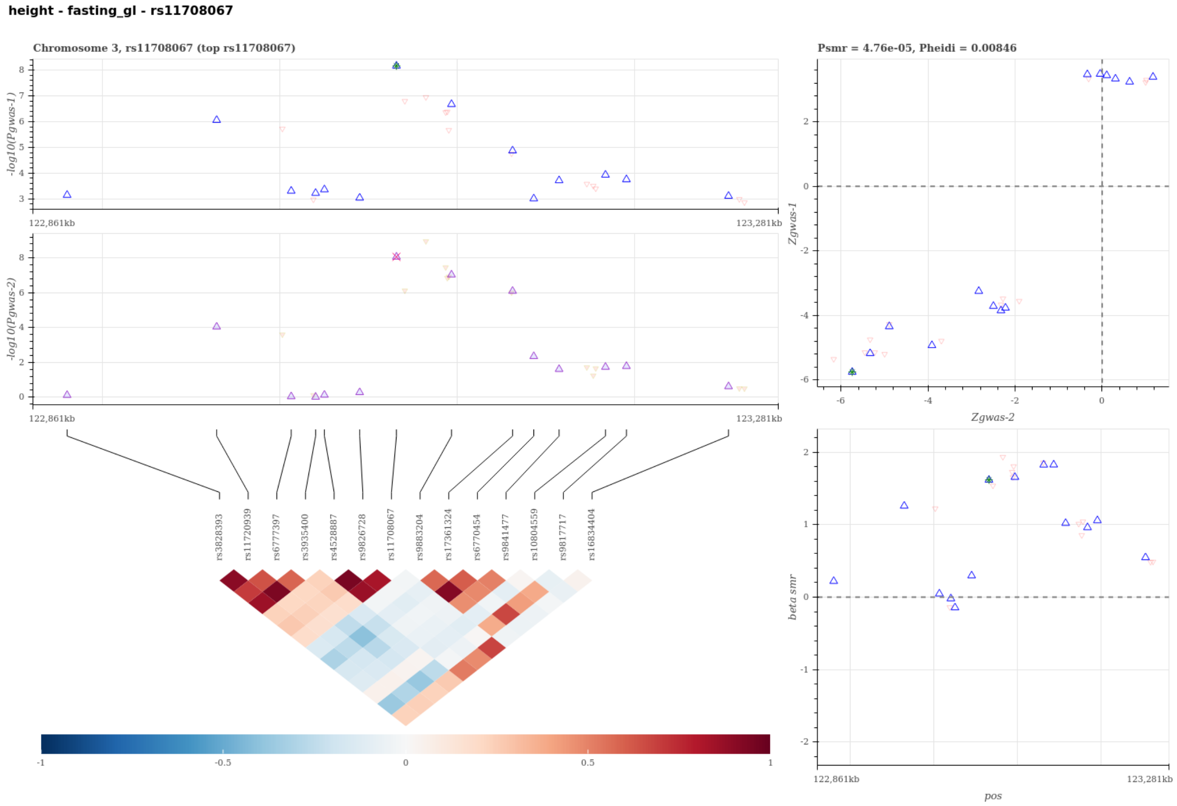


Figure S48: Visualization of the SMR-HEIDI test results for a single shared causal variant between HEIGHT and FastingGlucose. A statistical significant SMR test and non-significant HEIDI test suggest that FastingGlucose (gwas-2) shares the same single causal variant with HEIGHT (gwas-1). The top left panel compares the regional association results in -log scale p-values between HEIGHT and FastingGlucose. The top right panel compares the regional Z-statistics of the two association scans for the SMR test, and the bottom right panel visualizes the SMR causal effect estimates, i.e. the ratio of the genetic effect on HEIGHT to that on FastingGlucose, for the heterogeneity HEIDI test. The Bottom left panel visualizes the linkage disequilibrium structure of the region. In the plots, each triangle represents a SNP, where the pruned SNP for the SMR-HEIDI test are shown in bigger blue triangles and the top gwas-1 SNP is marked with extra colors.

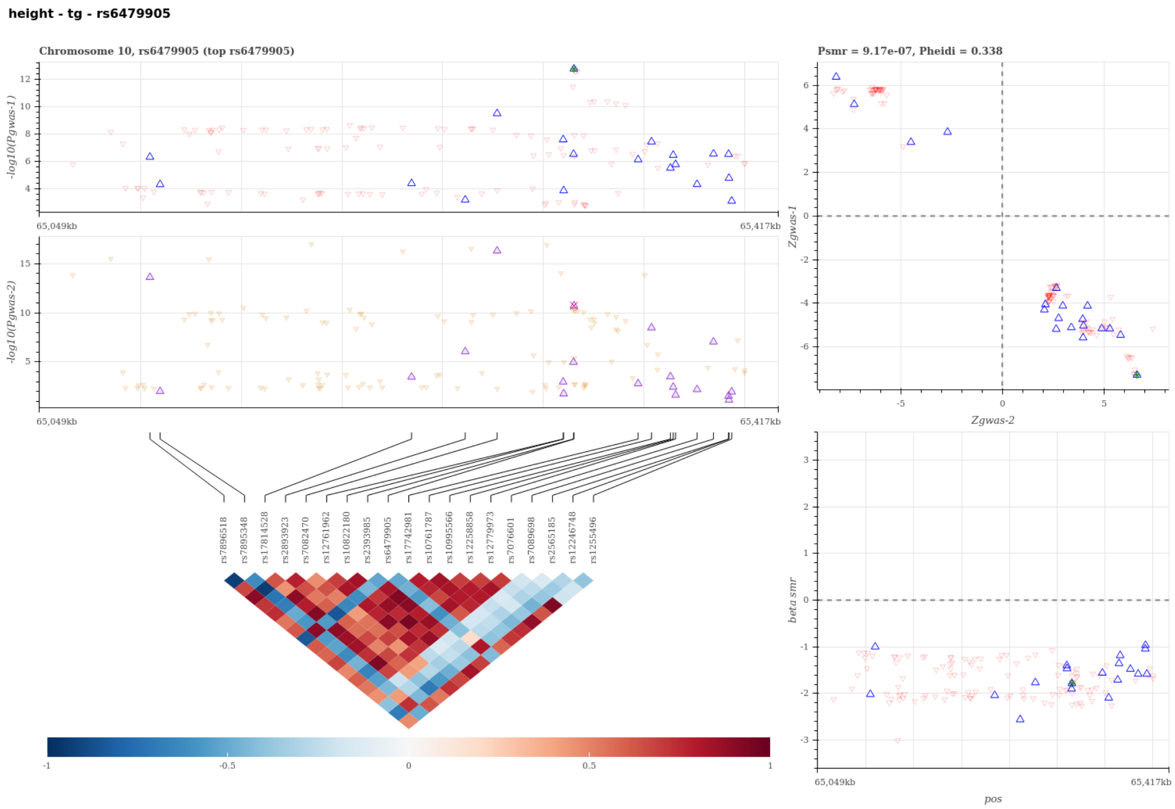


Figure S49: Visualization of the SMR-HEIDI test results for a single shared causal variant between HEIGHT and Triglycerides. A statistical significant SMR test and non-significant HEIDI test suggest that Triglycerides (gwas-2) shares the same single causal variant with HEIGHT (gwas-1). The top left panel compares the regional association results in -log scale p-values between HEIGHT and Triglycerides. The top right panel compares the regional Z-statistics of the two association scans for the SMR test, and the bottom right panel visualizes the SMR causal effect estimates, i.e. the ratio of the genetic effect on HEIGHT to that on Triglycerides, for the heterogeneity HEIDI test. The Bottom left panel visualizes the linkage disequilibrium structure of the region. In the plots, each triangle represents a SNP, where the pruned SNP for the SMR-HEIDI test are shown in bigger blue triangles and the top gwas-1 SNP is marked with extra colors.

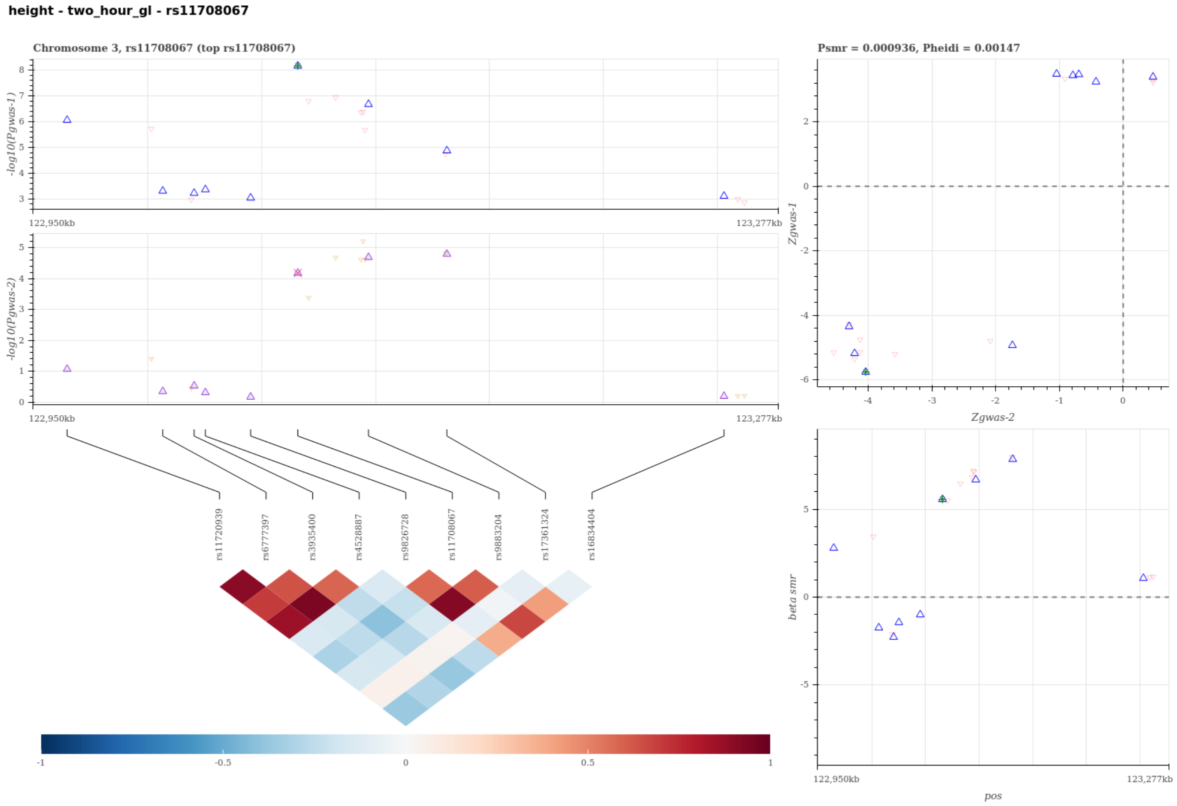


Figure S50: Visualization of the SMR-HEIDI test results for a single shared causal variant between HEIGHT and TwoHourGlucose. A statistical significant SMR test and non-significant HEIDI test suggest that TwoHourGlucose (gwas-2) shares the same single causal variant with HEIGHT (gwas-1). The top left panel compares the regional association results in -log scale p-values between HEIGHT and TwoHourGlucose. The top right panel compares the regional Z-statistics of the two association scans for the SMR test, and the bottom right panel visualizes the SMR causal effect estimates, i.e. the ratio of the genetic effect on HEIGHT to that on TwoHourGlucose, for the heterogeneity HEIDI test. The Bottom left panel visualizes the linkage disequilibrium structure of the region. In the plots, each triangle represents a SNP, where the pruned SNP for the SMR-HEIDI test are shown in bigger blue triangles and the top gwas-1 SNP is marked with extra colors.

**height - ed_years - rs6479905**

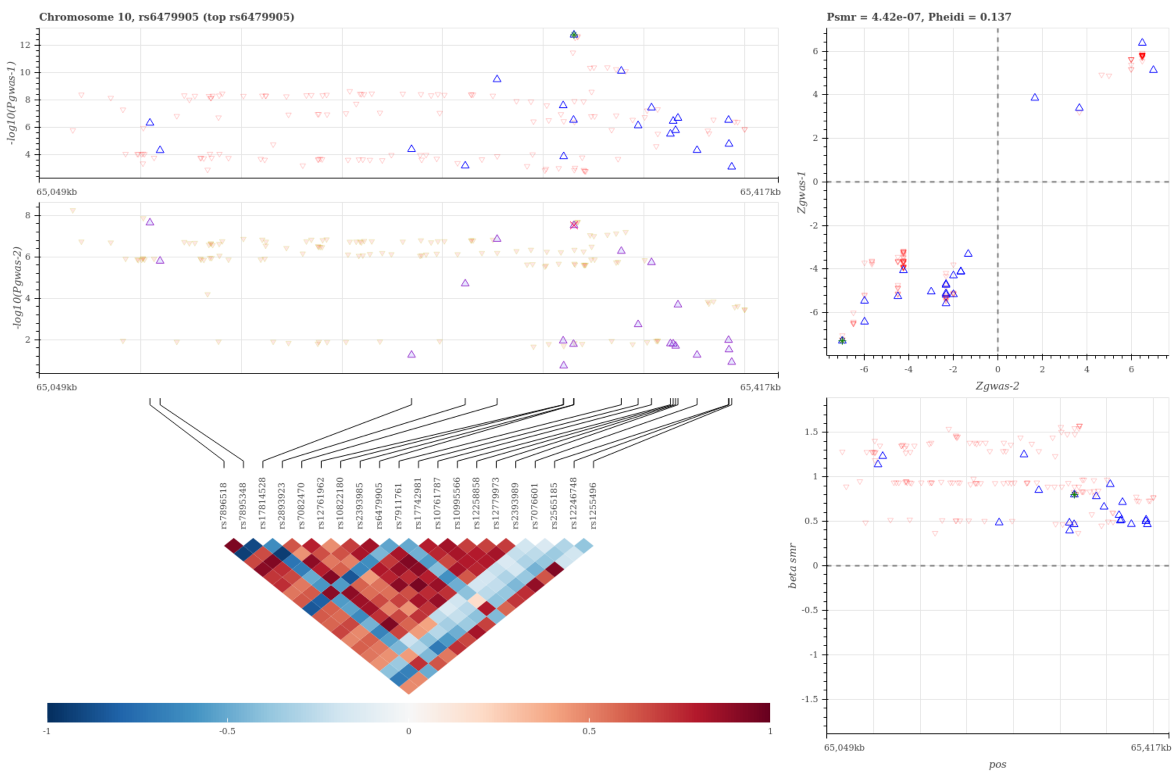


Figure S51: Visualization of the SMR-HEIDI test results for a single shared causal variant between HEIGHT and YearsOfEducation. A statistical significant SMR test and non-significant HEIDI test suggest that YearsOfEducation (gwas-2) shares the same single causal variant with HEIGHT (gwas-1). The top left panel compares the regional association results in -log scale p-values between HEIGHT and YearsOfEducation. The top right panel compares the regional Z-statistics of the two association scans for the SMR test, and the bottom right panel visualizes the SMR causal effect estimates, i.e. the ratio of the genetic effect on HEIGHT to that on YearsOfEducation, for the heterogeneity HEIDI test. The Bottom left panel visualizes the linkage disequilibrium structure of the region. In the plots, each triangle represents a SNP, where the pruned SNP for the SMR-HEIDI test are shown in bigger blue triangles and the top gwas-1 SNP is marked with extra colors.

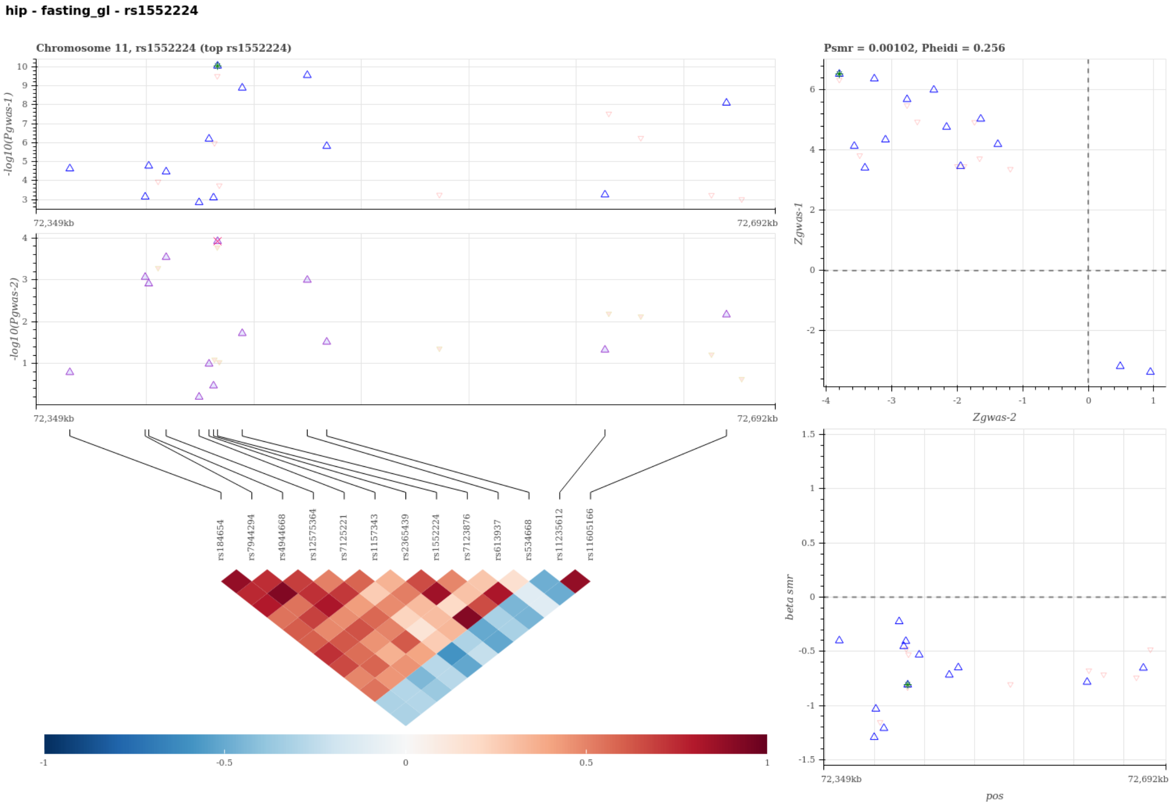


Figure S52: Visualization of the SMR-HEIDI test results for a single shared causal variant between HIP and FastingGlucose. A statistical significant SMR test and non-significant HEIDI test suggest that FastingGlucose (gwas-2) shares the same single causal variant with HIP (gwas-1). The top left panel compares the regional association results in -log scale p-values between HIP and FastingGlucose. The top right panel compares the regional Z-statistics of the two association scans for the SMR test, and the bottom right panel visualizes the SMR causal effect estimates, i.e. the ratio of the genetic effect on HIP to that on FastingGlucose, for the heterogeneity HEIDI test. The Bottom left panel visualizes the linkage disequilibrium structure of the region. In the plots, each triangle represents a SNP, where the pruned SNP for the SMR-HEIDI test are shown in bigger blue triangles and the top gwas-1 SNP is marked with extra colors.

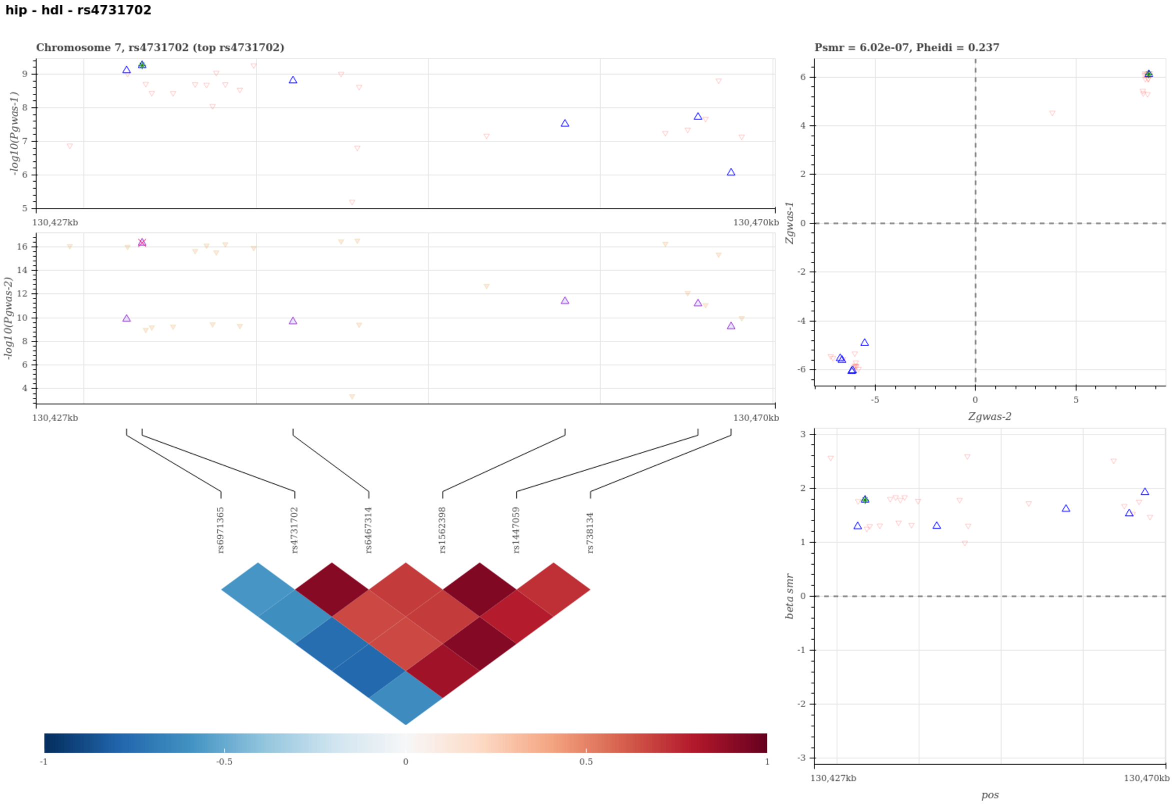


Figure S53: Visualization of the SMR-HEIDI test results for a single shared causal variant between HIP and HDL. A statistical significant SMR test and non-significant HEIDI test suggest that HDL (gwas-2) shares the same single causal variant with HIP (gwas-1). The top left panel compares the regional association results in -log scale p-values between HIP and HDL. The top right panel compares the regional Z-statistics of the two association scans for the SMR test, and the bottom right panel visualizes the SMR causal effect estimates, i.e. the ratio of the genetic effect on HIP to that on HDL, for the heterogeneity HEIDI test. The Bottom left panel visualizes the linkage disequilibrium structure of the region. In the plots, each triangle represents a SNP, where the pruned SNP for the SMR-HEIDI test are shown in bigger blue triangles and the top gwas-1 SNP is marked with extra colors.

**hip - tg - rs972283**

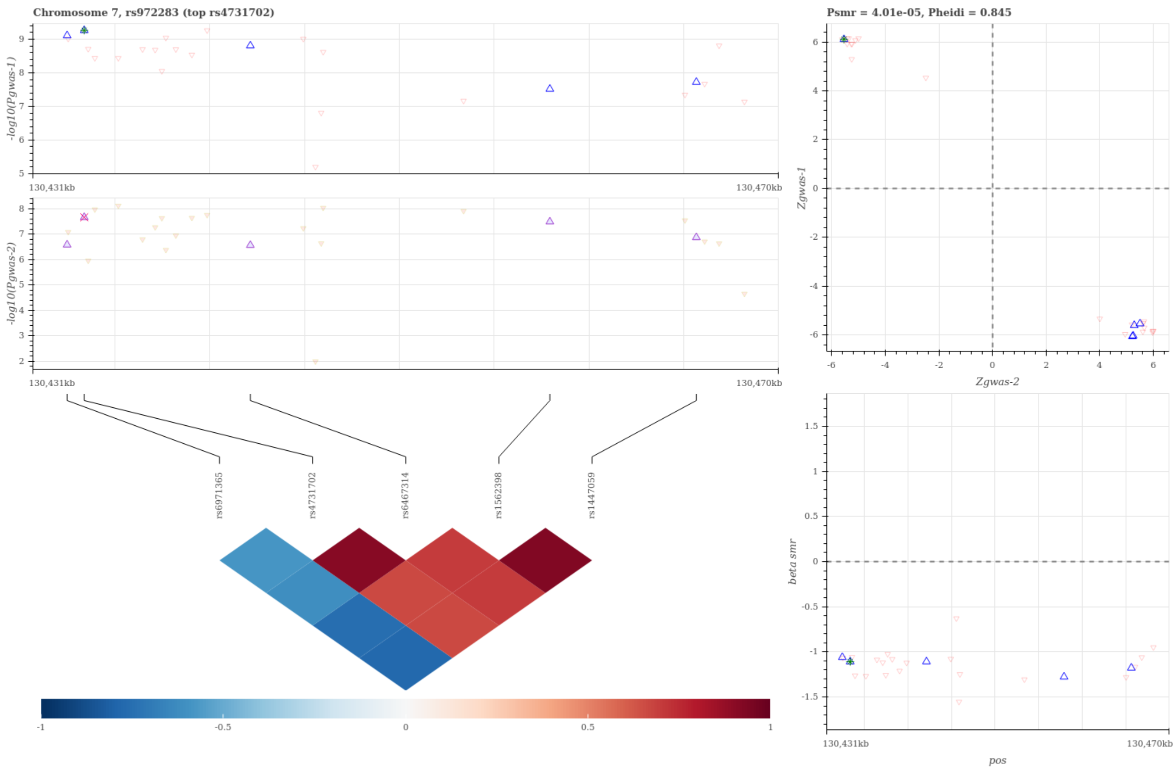


Figure S54: Visualization of the SMR-HEIDI test results for a single shared causal variant between HIP and Triglycerides. A statistical significant SMR test and non-significant HEIDI test suggest that Triglycerides (gwas-2) shares the same single causal variant with HIP (gwas-1). The top left panel compares the regional association results in -log scale p-values between HIP and Triglycerides. The top right panel compares the regional Z-statistics of the two association scans for the SMR test, and the bottom right panel visualizes the SMR causal effect estimates, i.e. the ratio of the genetic effect on HIP to that on Triglycerides, for the heterogeneity HEIDI test. The Bottom left panel visualizes the linkage disequilibrium structure of the region. In the plots, each triangle represents a SNP, where the pruned SNP for the SMR-HEIDI test are shown in bigger blue triangles and the top gwas-1 SNP is marked with extra colors.

#### Supplemental Tables

**Table S1: MV analysis results for GIANT2013 data with replication on GIANT2015 data, UKB data, and GIANT2015+UKB meta-analysis data (Excel file).**

**Table S2: Extended results for 49 SNPs discovered using MV-only approach in GIANT2015 data (Excel file).**

**Table S3: Correlation matrix for GIANT 2013, GIANT2015 and UKB (Excel file).**

**Table S4: Pleiotropy database records with the anthropometric traits according to PhenoScanner (FDR $<$ 5%) for the 49 novel loci (Excel file).**

**Table S5: SMR-HEIDI test results for the prediction of candidate gene and detection of shared genetic basis across complex traits (Excel file).**

### References


1. Olson, C. L. On choosing a test statistic in multivariate analysis of variance. *Psychological Bulletin* **83**, 579 (1976).

2. Zhu, X. *et al.* Meta-analysis of correlated traits via summary statistics from GWASs with an application in hypertension. *American journal of human genetics* **96**, 21–36 (2015).

3. Yang, J. *et al.* Conditional and joint multiple-SNP analysis of GWAS summary statistics identifies additional variants influencing complex traits. *Nature genetics* **44**, 369–75– S1–3 (2012).

4. Ning, Z. *et al.* A selection operator for summary association statistics reveals allelic heterogeneity of complex traits. *The American Journal of Human Genetics* **101**, 903–912 (2017).

5. Staley, J. R. *et al.* Phenoscanner: a database of human genotype–phenotype associations. *Bioinformatics* **32**, 3207–3209 (2016).

6. Zhu, Z. *et al.* Integration of summary data from GWAS and eQTL studies predicts complex trait gene targets. *Nature genetics* **48**, 481–487 (2016).

7. Westra, H.-J. *et al.* Systematic identification of trans eqtls as putative drivers of known disease associations. *Nature genetics* **45**, 1238 (2013).
